# TIGER: The gene expression regulatory variation landscape of human pancreatic islets

**DOI:** 10.1101/2021.05.26.445616

**Authors:** Lorena Alonso, Anthony Piron, Ignasi Morán, Marta Guindo-Martínez, Sílvia Bonàs-Guarch, Goutham Atla, Irene Miguel-Escalada, Romina Royo, Montserrat Puiggròs, Xavier Garcia-Hurtado, Mara Suleiman, Lorella Marselli, Jonathan L.S. Esguerra, Jean-Valéry Turatsinze, Jason M. Torres, Vibe Nylander, Ji Chen, Lena Eliasson, Matthieu Defrance, Ramon Amela, MAGIC, Hindrik Mulder, Anna L. Gloyn, Leif Groop, Piero Marchetti, Decio L. Eizirik, Jorge Ferrer, Josep M. Mercader, Miriam Cnop, David Torrents

## Abstract

GWAS have identified more than 700 genetic signals associated with type 2 diabetes (T2D). To gain insight into the underlying molecular mechanisms, we created the Translational human pancreatic Islet Genotype tissue-Expression Resource (TIGER), aggregating >500 human islet RNA-seq and genotyping datasets. We imputed genotypes using 4 reference panels and meta-analyzed cohorts to improve coverage of expression quantitative trait loci (eQTL) and developed a method to combine allele-specific expression across samples (cASE). We identified >1 million islet eQTLs (56% novel), of which 53 colocalize with T2D signals (60% novel). Among them, a low-frequency allele that reduces T2D risk by half increases *CCND2* expression. We identified 8 novel cASE colocalizations, among which an *SLC30A8* T2D associated variant. We make all the data available through the open-access TIGER portal (http://tiger.bsc.es), which represents a comprehensive human islet genomic data resource to elucidate how genetic variation affects islet function and translate this into therapeutic insight and precision medicine for T2D.

## Introduction

Diabetes is a complex metabolic disease, characterized by elevated blood glucose levels, that affects more than 463 million people worldwide ^1^. Type 2 diabetes (T2D) accounts for >85% of diabetes cases and is strongly related to age, obesity and sedentary lifestyles. Epidemiologic studies forecast a 40% increase in prevalence by 2030 ^2–4^. This makes the study and understanding of diabetes a top research and healthcare priority. Progressive pancreatic islet dysfunction is central to the majority of diabetes forms and hereby key to gain insights into the disease pathophysiology.

Great efforts have been dedicated to uncover the link between genetic variation and complex disease susceptibility through large-scale genetic studies. For T2D, >700 genetic loci have been identified to date ^5–8^. The vast majority of variants in these loci do not disrupt protein coding sequences ^9,10^. Thus, the mechanisms by which these variants influence predisposition to disease remain to be elucidated. As the number of newly identified risk variants keeps increasing, their functional interpretation constitutes the main bottleneck to gain insights into the underlying molecular mechanisms and thus, to develop more effective and targeted preventive and therapeutic strategies ^11^.

To provide functional interpretation of non-coding variation, large international efforts have generated and integrated genomic, transcriptomic and epigenomic data from a large variety of healthy and diseased samples to build comprehensive and genome-wide maps of functional annotations. Among others, the Genotype-Tissue Expression (GTEx) project uses expression quantitative trait loci (eQTL) analysis to link genetic variation with gene expression across 54 different human tissues ^12^. The Roadmap Epigenomics Mapping project ^13^ and the International Human Epigenome project ^14^ also provide a broad characterization of epigenomic signatures in a variety of tissues and cell types.

The functional interpretation of genetic variants, which are usually associated with moderate or small effect sizes, requires tools and resources that focus on cells and tissues that are impacted in the disease of interest. The islets of Langerhans, which are clusters of specialized endocrine cells that are essential to maintain glucose homeostasis, play a central role in the etiology of T2D ^15,16^. Because human islets are difficult to obtain ^17–19^, large multi-tissue resources such as GTEx do not contain islet data and at best use whole pancreas as a proxy, despite the fact that >97% of the pancreatic tissue consists of exocrine cells that mask islet signals ^20^. Hence, the development of publicly available resources and tools that include data on islet tissue is essential to translate T2D genetic signals into molecular and physiological mechanisms.

The first studies of eQTL in human islets pinpointed genes that might be influenced by genetic variants and thus possibly mediate T2D risk ^21,22^. Despite the small number of samples, they identified a few loci linked to differential expression of islet genes, which were enriched in genome-wide association study (GWAS) signals for T2D and related traits. More recently, a multinational consortium effort, InsPIRE, generated a largest islet eQTL study with a sample size of 420 islet donors, which identified 46 T2D GWAS signals that colocalize with islet eQTL ^23^.

To further expand the understanding of human islet regulatory genomics, and its role in T2D the Horizon 2020 T2DSystems consortium (https://www.t2dsystems.eu/) gathered the most extensive collection to date of human islet samples with gene expression, epigenomic data, genotypic and phenotypic information, with a total of 514, from which 207 samples were analyzed by the InsPIRE consortium. In this study, we discovered 40 T2D risk signals that colocalize with eQTL or ASE signals by improving genotype imputation methods and analyses and by developing a new method to combine allele-specific expression (cASE) across samples, knowledge previously unknown.

Importantly, the results from this study are made publicly available to the community through the Translational human pancreatic Islet Genotype tissue-Expression Resource (TIGER, http://bsc.tiger.es) portal (Figure 1A). This portal integrates the newly generated data with publicly available T2D genomic and genetic resources to facilitate the translation of genetic signals into their functional and molecular mechanisms.

**Figure 1:**
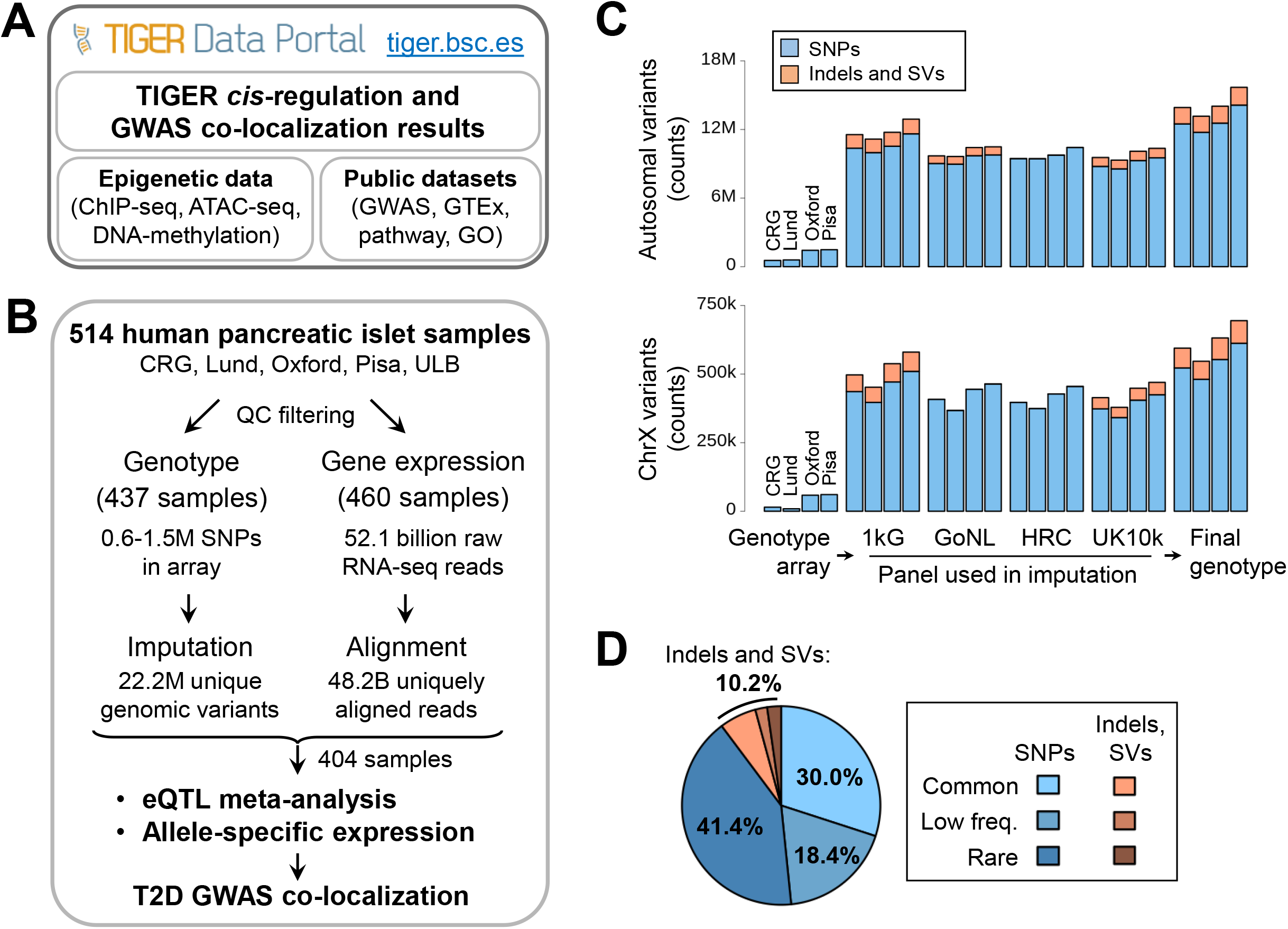
Project overview and genotype imputation. **A)** Overview of the TIGER data portal. **B)** Datasets of the T2DSystems consortium and project workflow. **C)** Multi-panel genotype imputation identified 13.1-15.7M autosomal variants (top) and 550-700k chrX variants (bottom), with **D)** a large proportion of low frequency (MAF 1-5%) and rare (<1%) variants, including 10.2% of Structural Variants (SVs), including small indels and large SVs.

## Results

### A catalogue of genetic variation and gene expression in human pancreatic islets

To study gene expression and the effects of genetic variation in human pancreatic islets, we obtained newly generated and published human islet data from 514 organ donors of European background, distributed across five cohorts (Center for Genomic Regulation, Lund University, University of Oxford/University of Alberta, Edmonton, Università di Pisa and Université Libre de Bruxelles).

The DNA of 307 new samples was isolated, sequenced and genotyped (Suppl. Table S1, Suppl. Methods) and aggregated to be harmonized with existing data from 207 samples. After quality control, filtering of RNA-seq and genotyping array data (Suppl. Methods), 404 human islet samples remained with high quality genotypes and RNA-seq data (Figure 1B).

To fully characterize the genetic variation present in the samples, genotype imputation was performed separately for each cohort using four different reference panels as previously described ^7,24^ (1000 genomes ^25^, GoNL ^26^, the Haplotype Reference Consortium ^27^ and UK10K ^28^). The results were integrated by selecting, for each variant, the imputed genotypes from the reference panel that achieved the best imputation quality (IMPUTE2 info score > 0.7, Suppl. Methods). This allowed imputation of >22 million unique high-quality genetic variants across all samples, 10% of which were indels and small structural variants, and more than 1.05 million variants in chromosome X (Figure 1C) (Suppl. Table S2, Suppl. Methods). Notably, this strategy allowed accurate imputation of 4 million low-frequency (minor allele frequency (MAF) between 0.05 and 0.01) and 10 million rare (0.01>MAF>0.001) variants (Figure 1D).

Additionally, we performed RNA-seq in 514 samples 460 of which were retained after stringent quality control, including >52 billion raw short reads. We uniquely aligned more than 48 billion reads (median of 93 million per sample) (Suppl. Table S3), which allowed us to observe >22K genes expressed at >0.5 transcripts per million (TPM) (Suppl. Methods).

### An atlas of eQTLs in human pancreatic islets

To explore the association between genetic variation and gene expression, we performed an eQTL meta-analysis across four cohorts. First, we performed a *cis*-eQTL analysis, using data from each cohort independently (Suppl. Methods). For each analysis, we corrected for known covariates (age, sex and body mass index (BMI)), genetically derived principal components, and PEER factors for hidden confounding factors ^29^. The eQTL results from each of the four cohorts were then meta-analyzed (Figure 2A). This resulted in >1.11 million significant eQTLs in more than 21,115 eGenes (12,802 protein coding genes, 8,313 non-coding) at 5% false discovery rate (FDR) after Benjamini-Hochberg correction for multiple testing ^30^ (Figure 2B). The quantile-quantile plot showed no baseline inflation in the results (Suppl. Figure S1). More than 12% of all significant eQTLs were small indels or larger structural variants, and this type of variation was the top associated variant for 14% of all genes. This is in line with what has been observed in primary human immune cell types in which indels comprised 12.5 % of the variants in the 95% credible sets for eQTLs in human immune cell types ^31^, and in GTEx, where it was observed that SVs have a stronger effect than SNVs ^32^.

**Figure 2:**
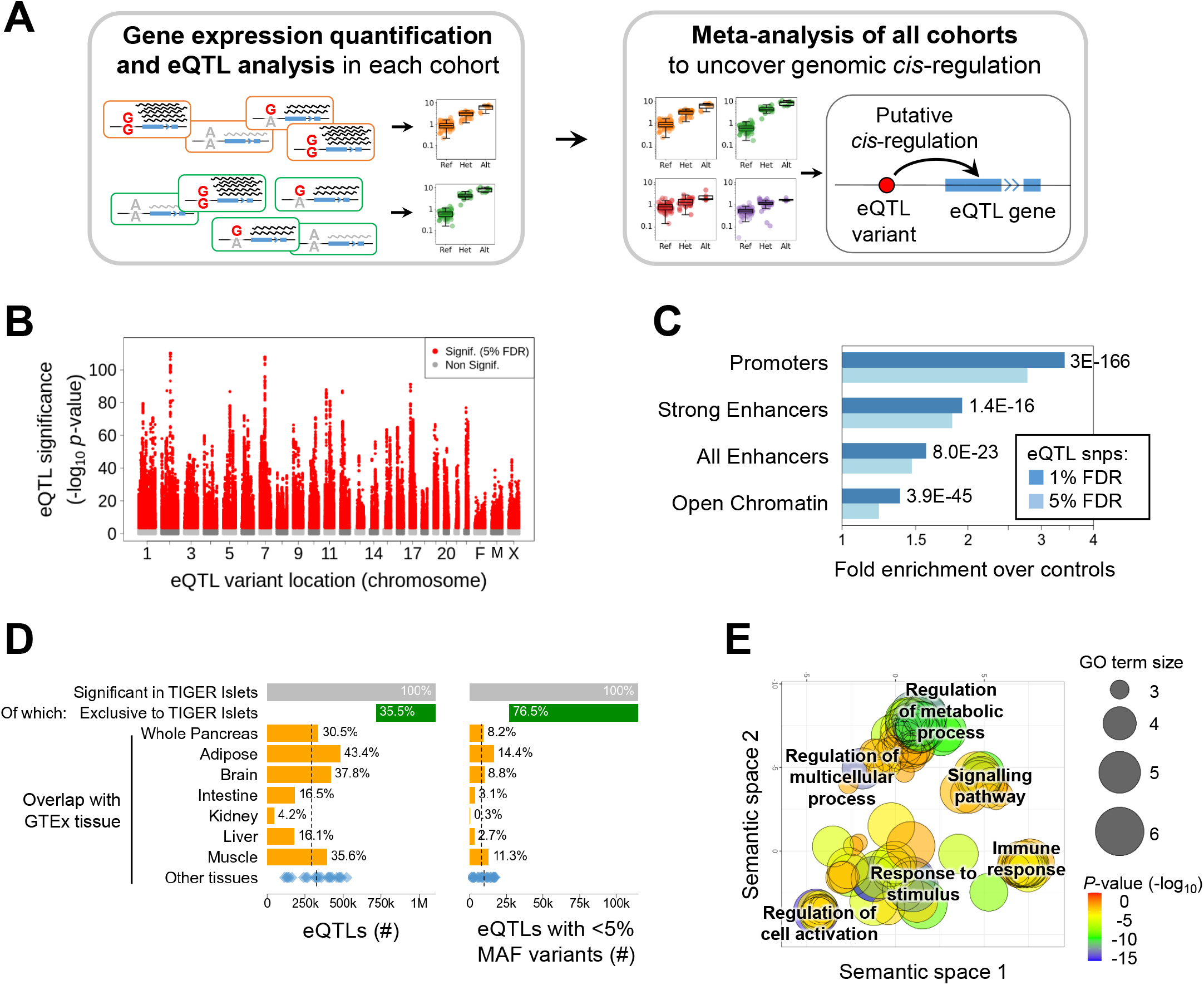
*Cis*-eQTL meta-analysis in human pancreatic islets. **A)** Overview of the metaanalysis. **B)** Manhattan plot of all eQTLs including chrX, analyzed with female-only (F) or male-only (M) samples, and jointly (X). **C)** Fold enrichment over controls of significant eQTL variants, in islet regulatory chromatin regions. *P*-values for 1% FDR eQTL enrichments are shown. **D)** Proportion of eQTLs novel in TIGER human islets (green) and previously found in GTEx project: tissues related to T2D aetiology (orange), other tissues (blue); means in dashed lines. Right panel restricted to low minor allele frequency (MAF) variants only. **E)** Gene ontology analysis of the genes of TIGER-specific eQTLs.

To assay the potential functional impact of the identified eQTL variants, we tested for their enrichment in human islet regulatory regions, defined by a variety of pancreatic islet chromatin assays ^10^. We observed that eQTL variants overlapped with gene promoters with very strong fold enrichment when compared with a control set of genetic variants (3.1-fold for 1% FDR eQTL variants, *p*=3×10^-166^) (Suppl. Methods), as well as with strong enhancers ^10^ (2-fold, *p*=1.4×10^-16^), and open-chromatin regions (1.4-fold, *p*=3.9×10^-45^) (Figure 2C, Suppl. Figure S2). These results are consistent with eQTL studies in other tissues ^12^.

Next, we contrasted the TIGER human islet results with the latest GTEx eQTL datasets, which analyzed 54 human tissues including whole pancreas, but not islets ^12^. Of all significant human islet eQTLs, 64.7% were also significant in at least one other GTEx tissue, whereas 35.3% were exclusive to human islets (Figure 2D, left panel). Only 30.5% of human islet eQTLs were also significant in whole pancreas in GTEx, an overlap similar to the rest of GTEx tissues (26% mean overlap with T2D related tissues, 29% with other tissues), highlighting that whole pancreas is not a better proxy for pancreatic islets compared to other tissues. In addition, when considering rare and low-frequency variants, the proportion of TIGER islet exclusive eQTLs increased to 76.5% (Figure 2D, right panel). These observations highlight again the importance of assaying human islets, since a sizeable proportion of the eQTLs cannot be found in other tissues. Interestingly, these observations also held true when we compared the TIGER results with the recently published eQTL analysis of 420 islet samples ^23^. Overall, 56.2% of the significant eQTLs were exclusive to our analysis (not assayed or non-significant in the InsPIRE study ^23^). Identification of eQTLs driven by low-frequency or rare variants may be more clinically impactful as significant low-frequency variants tend to have larger effects on disease risk and gene expression ^33^. Notably, the proportion of TIGER exclusive eQTLs increased to 74.6% for low-frequency variants, despite similar sample sizes between the studies. Overall, we identified 125,918 low-frequency eQTLs compared to 113,285 low-frequency eQTLs identified in the InsPIRE study (Suppl. Figure S3).

Gene ontology analysis of the significant human islet eQTL genes revealed signaling (including G-protein coupled receptor signaling) and metabolic regulation terms, albeit with moderate significance (Suppl. Figure S4). In contrast, comparing TIGER-specific eQTL genes against those also present in GTEx tissues revealed strong enrichment for these terms as well as “response to stimulus” or “regulation of cell activation”, and immune system related terms (including “lymphocyte/T-cell activation” and “regulation of immune system process”) (Figure 2E). This suggests that these novel eQTLs affected genes relevant to β-cell physiology, including some related to immune processes with potential relevance for type 1 diabetes ^34^.

### Islet eQTLs colocalize with T2D GWAS signals

To assess whether the identified eQTLs can help to identify effector transcripts for T2D risk variants, we investigated the intersection between *cis*-eQTLs and known T2D associations ^5–7^, by performing colocalization analyses using *COLOC* method^35^ (Suppl. Methods).

This analysis uncovered 49 eQTL variants associated with expression of 53 genes that significantly colocalized with T2D GWAS loci (Suppl. Table 4), of which 32 are novel (Table 1, Suppl. Figure S5). Interestingly, we identified three low-frequency variants, which may have large effect sizes, that colocalized with gene expression, suggesting a target gene and direction of effect, i.e,. whether the genetic variant is associated with increased or decreased gene expression. Among the 49 colocalizing signals (Suppl. Figure S5), rs77864822 (MAF=0.07) minor allele (G) was associated with higher *RMST* expression and decreased T2D risk (OR=0.93, *p*=2.2×10^-8^). By interrogating the latest GWAS study on glycemic traits ^36^, we observed that the protective allele was associated with decreased fasting glucose (beta=-0.024, *p*=4×10^-11^), reduced HbA1c (beta=-0.087, *p*=4.6×10^-4^), and reduced 2 hours glucose in an oral glucose tolerance test (beta=-0.064, *p*=2.4×10^-4^; Suppl. Table 4). The variant rs1531583 colocalized with *CPLX1* expression (Figure 3A-C). Interestingly, the same variant was associated with *PCGF3* but not with *CPLX1* gene expression in whole pancreas in GTEx (Figure 3B), demonstrating once again the importance of performing eQTL in the relevant tissue. A detailed analysis of enhancer chromatin marks in human islets showed that rs73221115 (r^2^=0.978 with rs1531583) and rs73221116 (r^2^=0.98 with rs1531583) had allele-specific H3K27ac binding suggesting that these two variants are the most likely causal variants of the *CPLX1* locus (Figure 3D-E). We also identified significant colocalization between the low-frequency variant rs76895963, known to reduce T2D risk by half ^37^, and increased *CCND2* expression in islets (Figure 3F-G). This variant was also associated with reduced fasting glucose (beta=-0.033, *p*=0.0017), HbA1c (beta=-0.042, *p*=3.6×10^-8^) and reduced 2 hours glucose in oral glucose tolerance test (beta=-0.095, *p*=0.01, Suppl. Table 4).

**Table 1.**
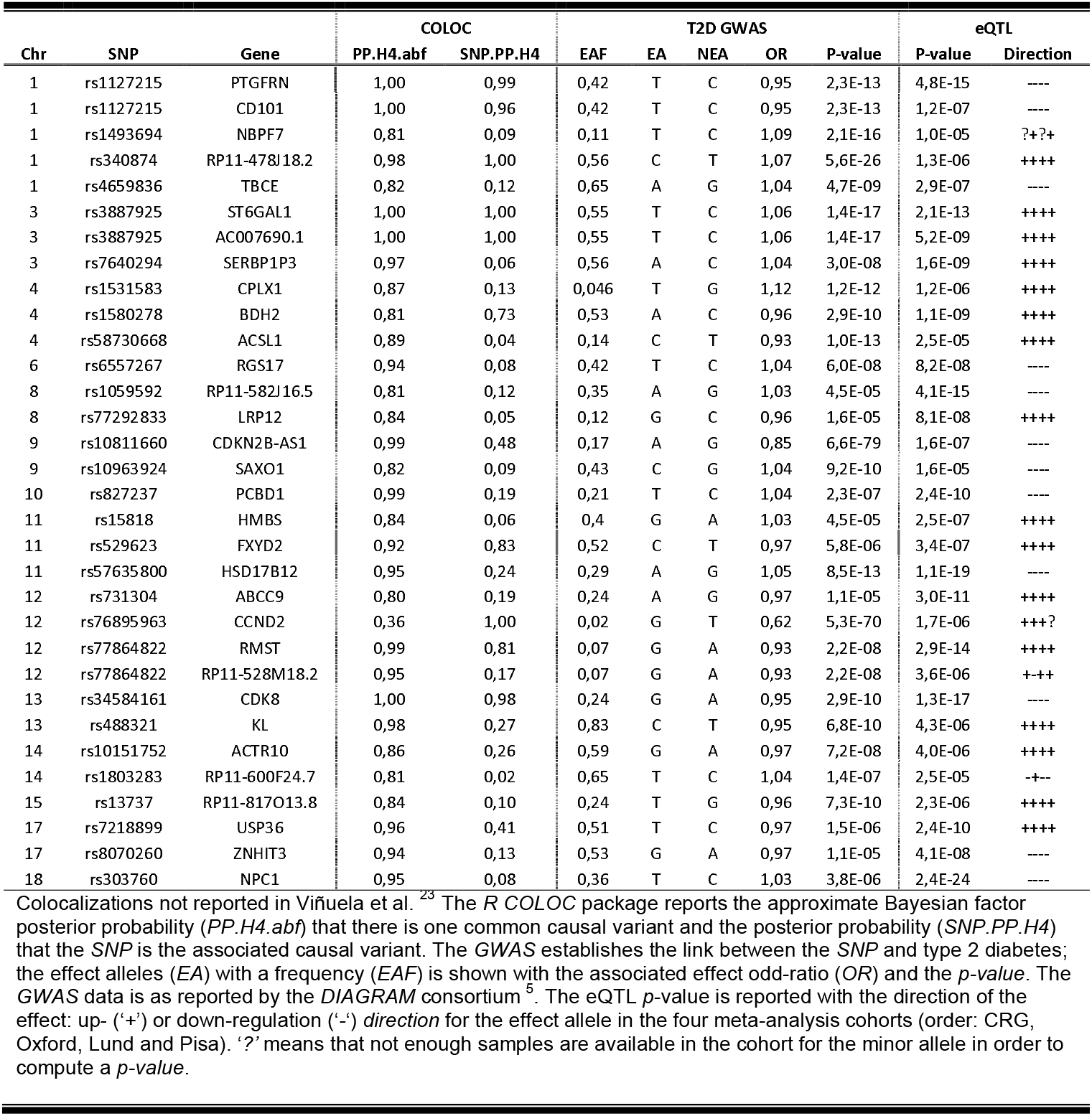
Novel human pancreatic islet colocalization of expression quantitative trait loci metaanalysis (eQTL) with type 2 diabetes (T2D) genome-wide association studies (GWAS).

**Figure 3:**
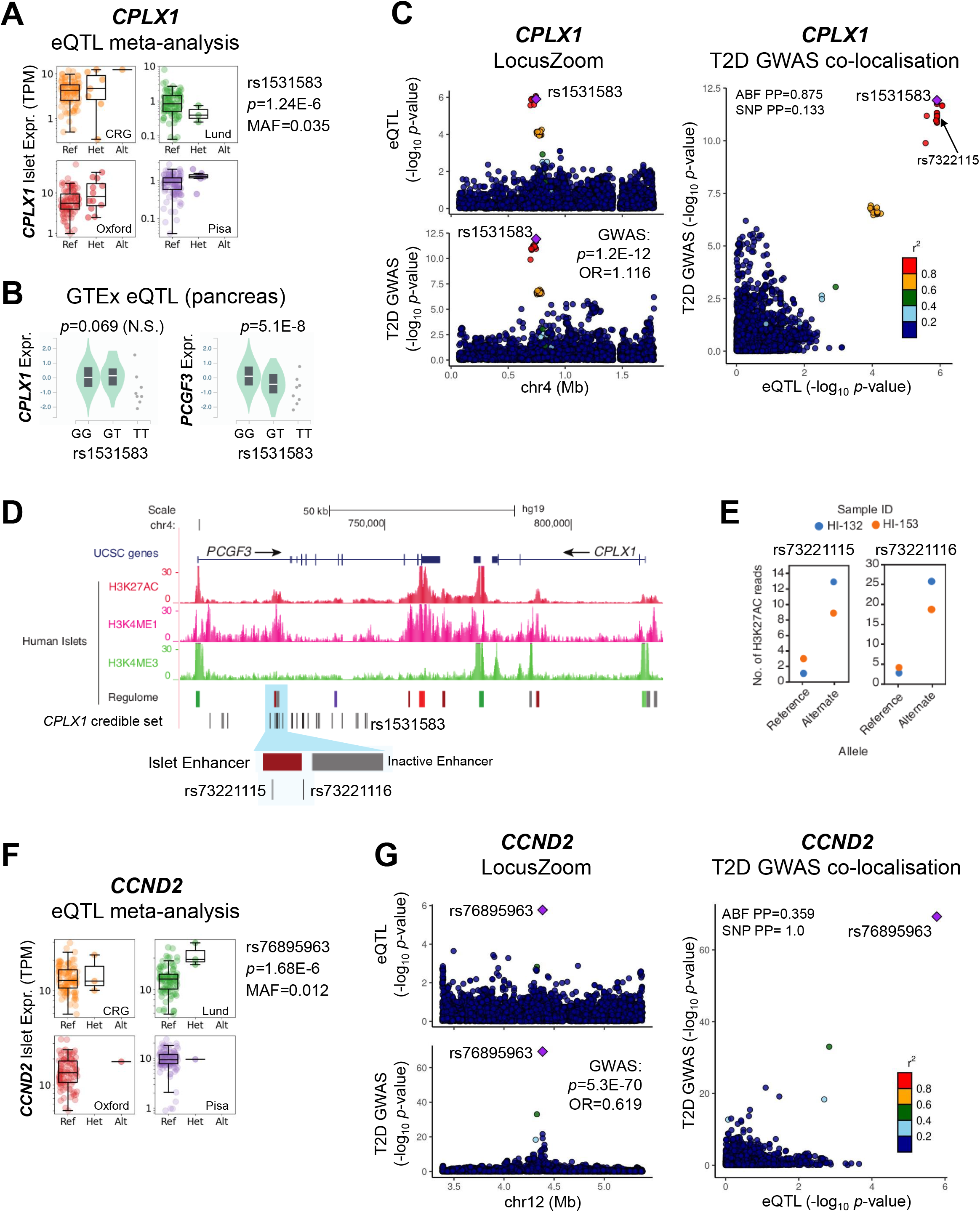
Examples of co-localization of pancreatic islets eQTLs with T2D GWAS. **A)** Boxplots representing expression of CPLX1 across different genotypes of variant rs1531583 in each of the cohorts and final meta-analysis results. **B)** rs1531583 was not significant in GTEx whole pancreas for *CPLX1*, but instead it was for *PCGF3* (bottom). **C)** LocusZoom plots of islet eQTL (top) and T2D GWAS (bottom) signals for rs1531583-*CPLX1*, and their co-localization (right). ABF: Approximate Bayes Factor, PP: Posterior Probability. **D)** An islet enhancer overlaps with rs73221115 and rs73221116, part of the *CPLX1* credible set of SNPs. **E)** Two human islet samples heterozygous for rs73221115 and rs73221116 showed allelic imbalance in their H3K27ac enhancer chromatin marks. **F)** eQTL meta-analysis of *CCND2* and the low frequency *cis*-regulatory variant rs76895963. **G)** Co-localization plots for rs76895963-*CCND2*, as in B).

### An atlas of cASE in human pancreatic islets

Preferential expression of mRNA copies containing one of the two alleles of a genetic variant (allele-specific expression, ASE) can result from *cis*-regulation. However, ASE can occur while the overall amount of expression of a gene remains constant, and therefore this type of regulation cannot be identified by conventional eQTL analysis.

We implemented a cASE pipeline for the analysis of ASE replicated across multiple samples that differ in age, gender, BMI and environmental factors, thereby likely to stem from *cis*-regulatory genetic variants (Figure 4A). cASE analysis complements eQTL analysis, and additionally controls for: a) environmental and batch effects, which are important confounding factors in eQTL studies ^38–43^, b) sample heterogeneity, which is prevalent in human islet samples ^44^, and c) *trans* effects, since these would affect the two alleles in the same manner and thus cannot result in ASE. cASE combines ASE from each sample into a single Z-score statistic that summarizes the overall ASE across the cohort of samples (Suppl. Methods, Suppl. Figure S6) ^45^. Variants that preferentially express the reference allele result in a positive Z-score and vice versa (Figure 4A).

**Figure 4:**
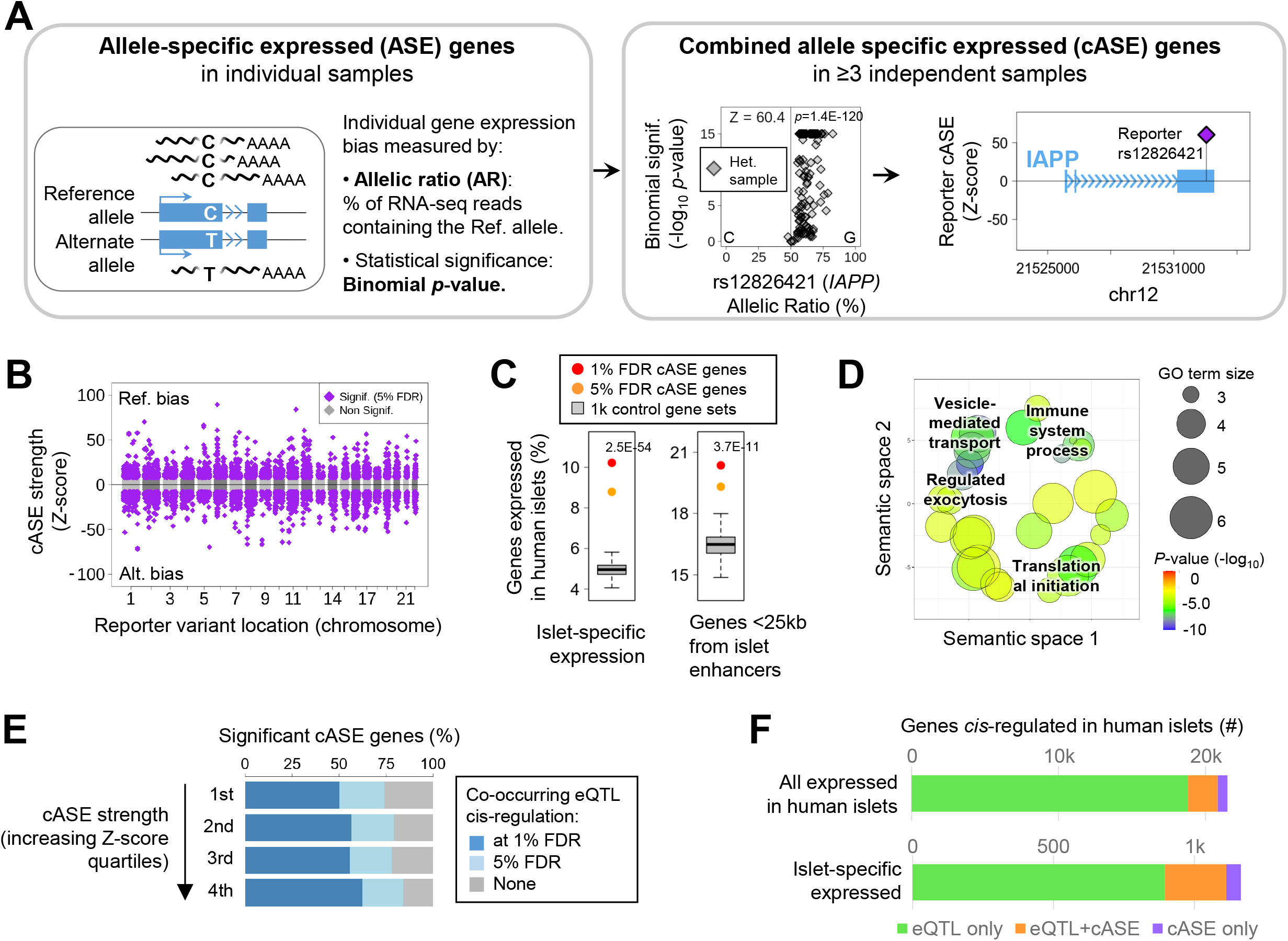
Combined ASE analysis in human islets. **A)** Overview of the cASE analysis, with *IAPP* as example of a gene with an imbalanced reporter variant, rs12826421. **B)** Manhattan plot of cASE, positive values refer to Reference-biased genes, negative to Alternate. **C)** Significant cASE genes are enriched for islet-specific expression and proximity to islet-regulatory regions. *P*-values for 1% FDR eQTL enrichments are shown. **D)** Gene ontology analysis of cASE significant genes. **E)** In genes with significant cASE, the proportion of also eQTL significant increased with increasing cASE magnitude. **F)** Total number of *cis*-regulated genes (top) and of islet-specific expressed (bottom), identified only by the eQTL analysis (green), cASE (purple), and by both (orange).

Using this strategy, we identified 2,707 genes with 5,271 reporter variants showing cASE in human islets, at 5% FDR (Figure 4B). The similar number of reference and alternate imbalanced variants (2,606 and 2,589, respectively) showed that alignment biases towards the reference allele were successfully controlled (see also Suppl. Figure S6B-E).

When comparing cASE genes against a set of non-significant genes (matched by gene expression level, Suppl. Methods), we observed that cASE genes were enriched for islet-specific expression (2.1-fold, *p*=2.5×10^-54^ at 1% FDR) and preferentially located near islet regulatory regions (1.23-fold, *p*=3.7×10^-11^) (Figure 4C). Gene ontology analysis (Suppl. Methods) revealed islet-specific terms such as “vesicle-mediated transport” and “regulated exocytosis”, (Figure 4D), related to insulin production and secretion in β-cells. As a notable example, the islet amyloid polypeptide gene (*IAPP*) was among the most imbalanced cASE genes. *IAPP* had 7 independent reporter SNPs at 1% FDR (Figure 4A, right panel; Suppl. Figure S7), all of which with strong imbalance towards the reference allele in the >100 independent samples that were heterozygous for the variants. Notably, there were no significant eQTLs for this gene, highlighting the complementarity between the two methods to identify regulatory variation. These findings highlight the potential of cASE to identify genes involved in regulating pancreatic islet physiology.

Given that eQTL and cASE analyses are complementary methods to detect genes affected by *cis*-regulation, we assessed the concordance between each of them. We first interrogated the proportion of genes with significant eQTL of all cASE genes across absolute Z-score quartiles (strength of imbalance), and observed that the proportion of eQTL genes increased with increasing Z-scores (Figure 4E), indicating that stronger cASE effects were more likely to be also identified in eQTL analysis, and showing a correlation between the two effects.

Of 2,707 cASE significant genes, 2,052 (75.8%) were detected in eQTL analysis, whereas 655 (24.2%) were detected uniquely through cASE (Figure 4F, top panel). The same trend was observed when considering only islet-specific expressed genes. Among 270 islet-specific significant eGenes detected by cASE, 218 were also detected by eQTL analysis, while the remaining 52 were exclusively found by cASE and not eQTL analysis (Figure 4F, bottom panel).

### Mapping distal cASE variants allows cASE colocalization analysis and implicates additional T2D effector genes

We next developed an approach to identify distal putative cASE regulatory variants by interrogating all variants within the same topologically associated domain as the reporter variant (i.e. the variant located in the transcribed gene region). For each candidate regulatory variant, we stratified samples between heterozygous and homozygous for the variant. We then recomputed cASE of the reporter variant (i.e., the transcribed variant) for each of the groups (Figure 5A). This approach allowed us to prioritize the candidate variant that had the highest reporter cASE when the candidate regulatory variant was also heterozygous, compared to when the regulatory variant was homozygous (Figure 5B, see Suppl. Methods).

**Figure 5:**
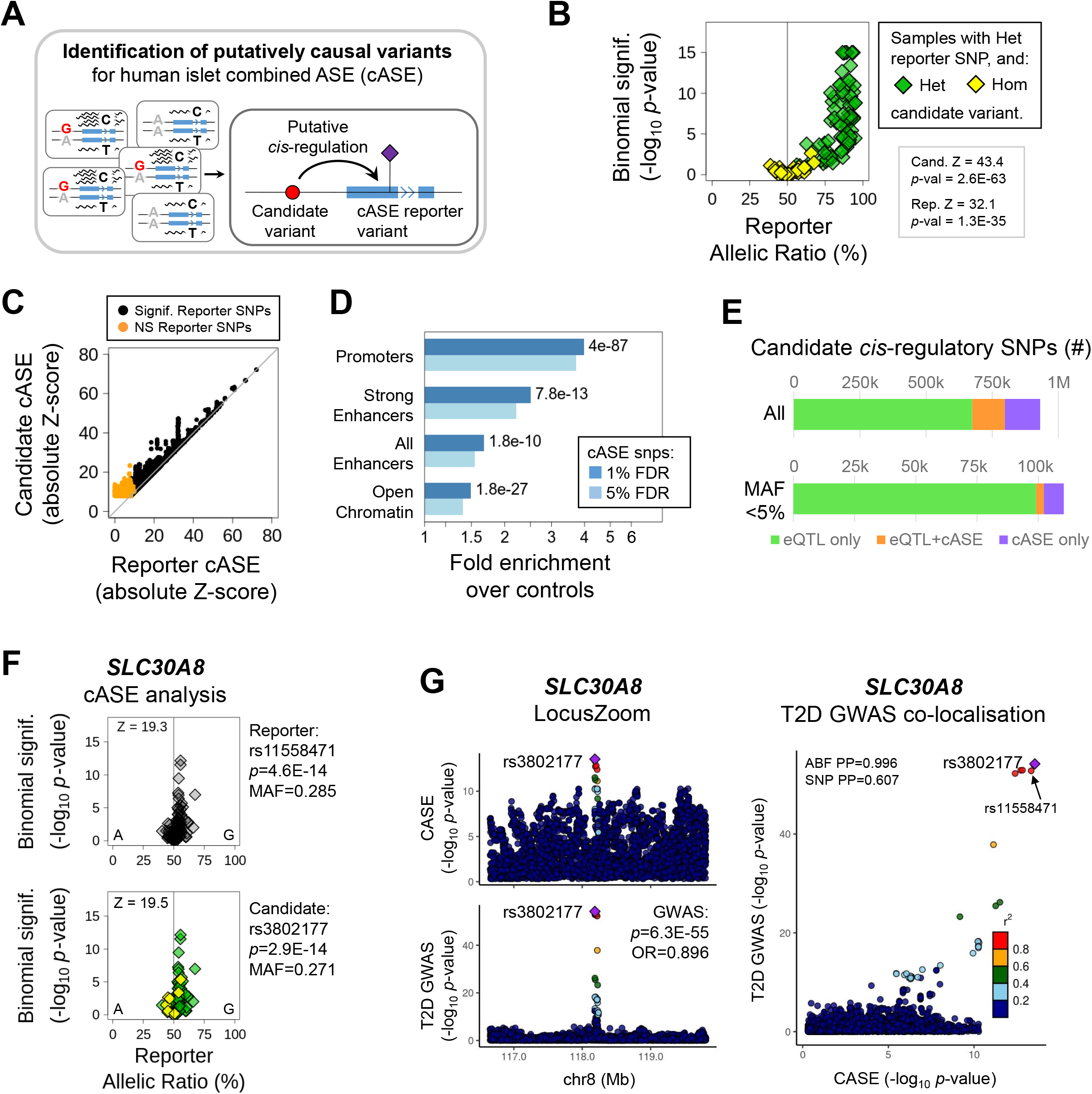
Identification of *cis*-regulatory variants in Combined ASE. **A)** Overview of the analysis. **B)** An example of *cis*-regulatory variant analysis; the samples Het for the candidate variant (green) have a higher cASE Z-score for the reporter SNP, while samples that are Hom for the candidate (yellow) do not show significant imbalance for the reporter SNP. **C)** Candidate variants often have stronger Z-scores than the reporters, including some reporter variants that were non-significant by themselves (orange). **D)** Fold enrichment over controls of significant cASE variants, in islet regulatory chromatin regions. *P*-values for 1% FDR cASE enrichments. **E)** Total number of candidate cis-regulatory variants (top) and low-frequency variants (bottom) identified by only the eQTL analysis (green), cASE (purple), and by both (orange). **F)** cASE analysis for *SLC30A8*, its best reporter SNP (top) and best candidate variant (bottom). **G)** LocusZoom plots of islet cASE (top) and T2D GWAS (bottom) signals for rs3802177-*SLC30A8*, and their co-localization (right). ABF: Approximate Bayes Factor, PP: Posterior Probability.

This analysis uncovered 256,981 putative regulatory variants for 3,425 genes, including 570 genes that had no significant reporter variants by themselves, but that did reach significance upon stratifying by genotype of regulatory variants (Figure 5C, orange points, see Suppl. Figure S8 for examples). To assay the potential functional impact of the identified reporter variants, we tested for their enrichment in human islet regulatory regions ^10^, observing overlap with gene promoters with very strong fold enrichment when compared with a control set of genetic variants (4-fold for 1% FDR eQTL variants, *p*=4×10^-87^) (Suppl. Methods), as well as with strong enhancers ^10^ (2.5-fold, *p*=7.8×10^-13^), and open-chromatin regions (1.5-fold, *p*=1.8×10^-27^) (Figure 5D). When comparing these *cis*-regulatory variants with the 1.11M eQTLs, we found 123,748 variants significant by both methods (3,138 with MAF<5%), and a further 133,233 (9,190 with MAF<5%) that were identified only by cASE (Figure 5E), showcasing the relevance of this analysis for enriching genetic *cis*-regulatory discovery.

Assigning statistical significance to cASE distal regulatory variants allowed us to test for colocalization between cASE regulatory variants and T2D GWAS variants. For each T2D GWAS locus, we assessed all regulatory variants for all imbalanced genes in the region and identified 14 colocalized locus-gene pairs (Table 2, Suppl. Figure S9). Of these, 6 had also been identified in eQTL/T2D GWAS colocalization analyses, showing consistency between the two methods. Interestingly, the 8 colocalizations identified by cASE alone suggested that these T2D variants may mediate disease risk by causing an imbalance in allelic expression, rather than altering overall gene expression. A notable example was the highly significant cASE observed in *SLC30A8* (rs11558471; *p*=2.9×10^-14^), which showed colocalization with a well-established T2D-associated variant (Figure 5F-G) (Suppl. Table 5) for which there was no eQTL colocalization. Thus, our novel cASE analysis uncovered additional disease-relevant genomic regulation and provides a potential biological mechanism underlying the association.

**Table 2.**
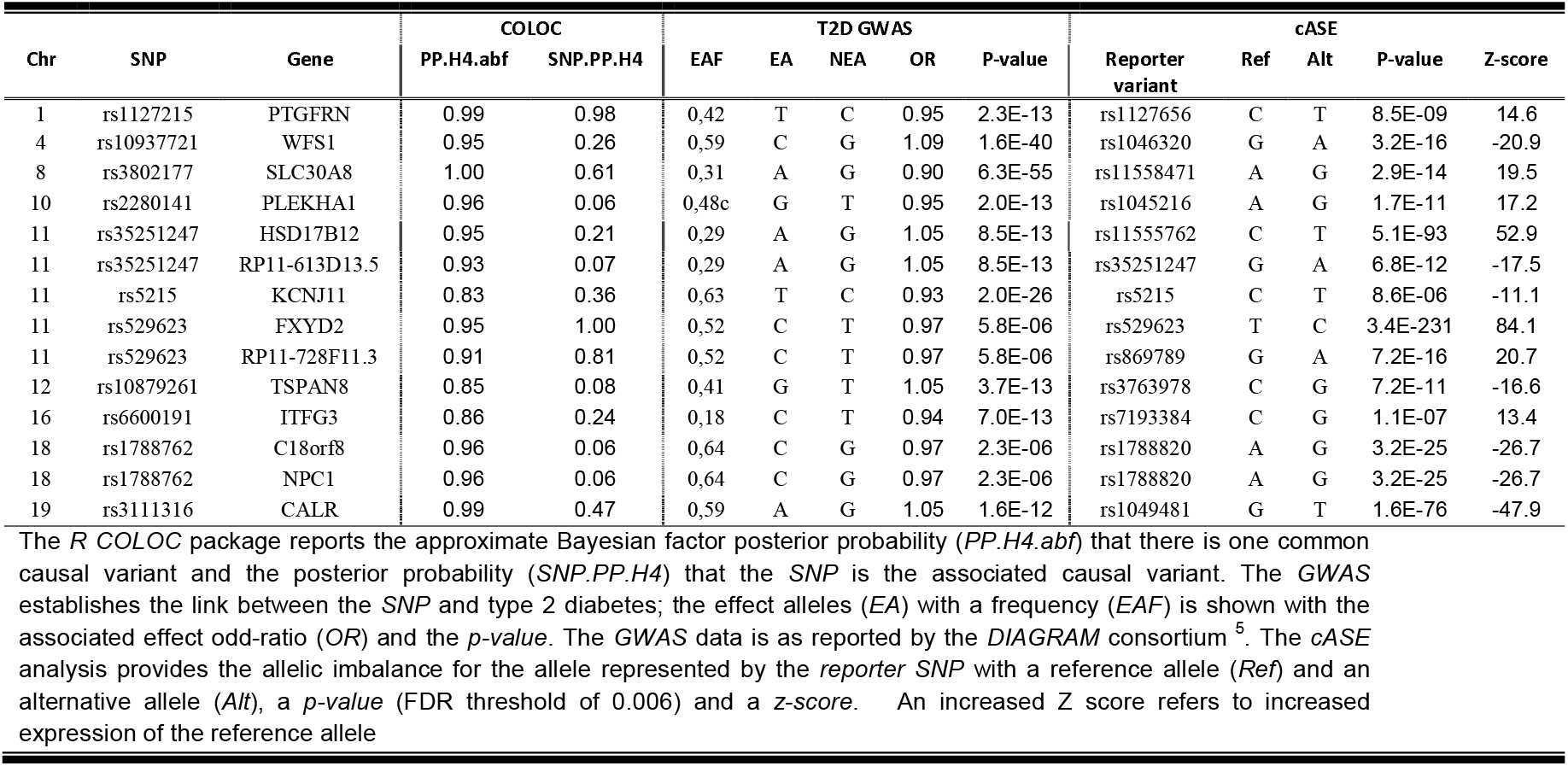
Colocalization of allele specific expression (*cASE*) with type 2 diabetes (T2D) genome-wide association study (GWAS).

### A web portal to explore regulatory variation and genomic pancreatic islet information

Finally, to provide the research community with a user-friendly open access tool to explore these findings and mine the molecular basis of complex diseases influenced by pancreatic islet biology, we created TIGER (http://tiger.bsc.es) (Figure 6). This portal integrates the results obtained in this study with other public genomic, transcriptomic and epigenomic pancreatic islet resources, as well as T2D GWAS meta-analysis summary statistics (Suppl. Methods).

**Figure 6:**
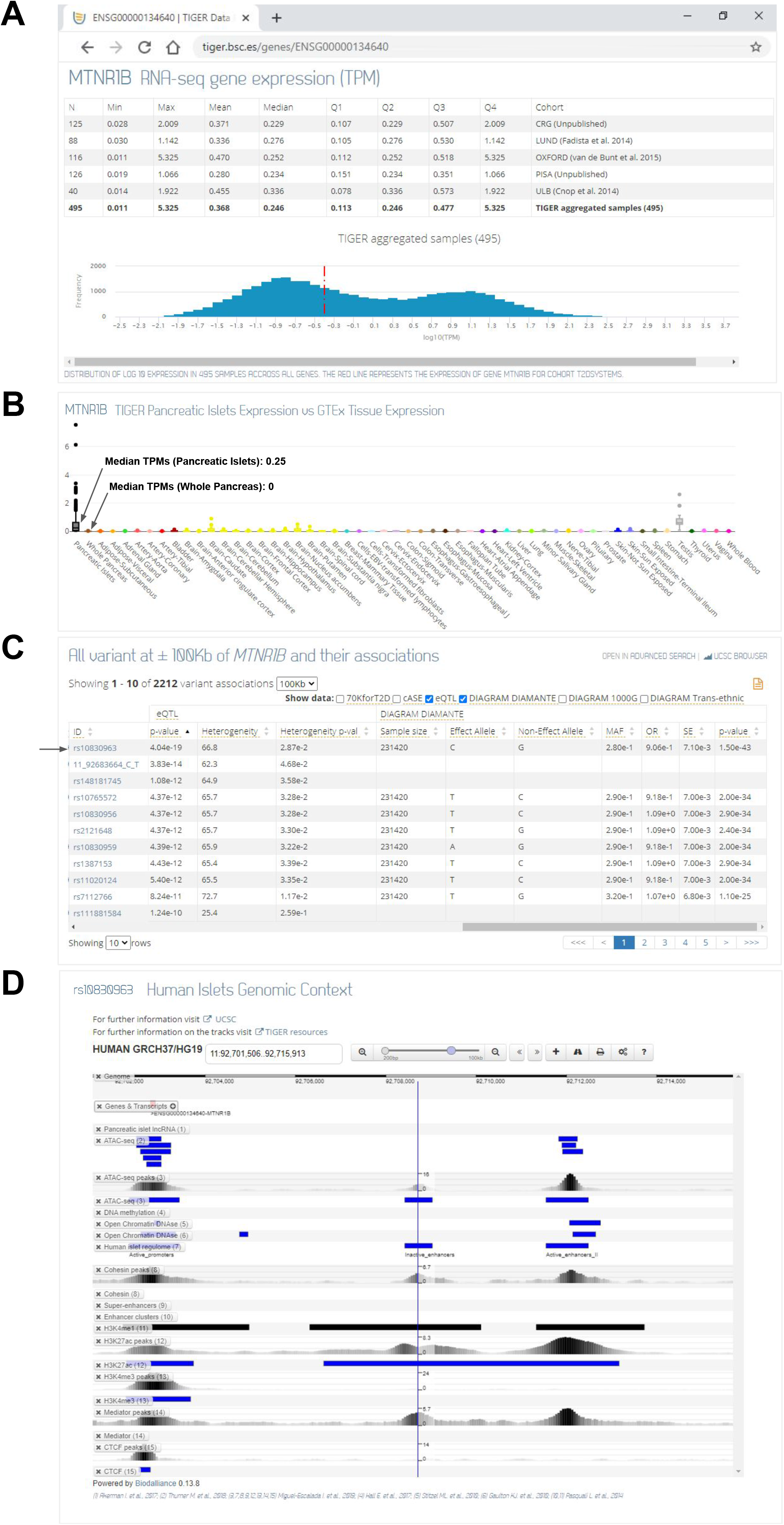
TIGER platform example. A) *MTNR1B* normalized log10(TPM) expression in islets; table (top) displays *MTNR1B* normalized TPM expression in each cohort and across the cohorts (bold); histogram (bottom) shows log10(TPM) gene expression distribution in 495 human islets samples, the red dashed line corresponds to *MTNR1B* log10(TPM) expression. B) *MTNR1B* normalized TPM expression in islets vs other GTEx tissues where each boxplot represents one tissue; *MTNR1B* has higher expression in pancreatic islets (black) compared to the whole pancreas (brown), which has almost no expression. C) Table showing the list of variants in a 100Kb window around *MTNR1B* and displaying results from either eQTL or DIAMANTE GWAS data sorted by ascending eQTL *p*-value; the eQTL variant rs10830963 (*p*=4.04×10^-19^) colocalizes with DIAMANTE (*p*=1.50×10^-43^). D) 15Kb human islet genomic context of variant rs10830963 (chr11:92708710); islet significant regions (black/blue boxes) and peaks are represented in each track, the blue line corresponds to rs10830963 position.

The TIGER website represents homogeneous gene expression levels from 446 RNA-seq pancreatic islet samples corrected for batch and covariate effects (Suppl. Figure S10), and enables comparison with GTEx expression data ^12^ (Suppl. Methods).

In addition to the eQTL and cASE results and to provide further functional assessment, we gathered islet regulatory information ^9,10,45^, methylation marks ^47,48^ and chromatin modification datasets ^49–51^. Further, to enable the translation of genetic variation to disease risk, we also integrated the latest T2D GWASs meta-analysis summary statistics ^5,7,52,53^ (Figure 1A).

The TIGER database currently contains expression and molecular data for >59K Gencode genes (version gencode.v23lift37 ^54^) and >27M variants. The portal allows users to perform both variant and gene centric queries. The results are displayed in a set of graphical tools and a genomic browser that will help visualize and interpret the molecular context of the query, as well as download the data. As a result of these efforts, the TIGER resource has already been used in recent studies ^55–57^.

As an example, we present the visualization of *MTNR1B*, a gene associated with type 2 diabetes and impaired insulin secretion ^58^. Although, this gene is lowly expressed in pancreatic islets (median 0.25 TPM) by comparison with other GTEx tissues it only shows low expression in testis (median 0.61 TPM) and brain (median 0.06 TPM) but none expression in whole pancreas and other tissues (median 0 TPM), thus highlighting the utility of this resource for studying human islet-specific expression (Figure 6A-B). A T2D risk associated locus has been previously described and fine-mapped ^5^ to a single variant (rs10830963, *p*=4.8×10^-43^, PP=0.99, Figure 6C, Suppl. Figure S5). Notably, this variant is located within islet H3K27ac peaks, suggesting potential regulatory implications of this variant (Figure 6D). In summary, the close lookup at this locus emphasizes that the TIGER portal can be easily used to interrogate gene expression, epigenomic and genomic variation regulatory landscape, providing an invaluable resource to the research community for the study of complex diseases affecting pancreatic islets.

## Discussion

By analyzing the largest dataset to date of pancreatic islets with gene expression and dense genotyping information we have uncovered one million significantly associated variant-gene pairs. Of all the associations we found, 35.3% were islet-specific, highlighting the importance of performing tissue-specific eQTL studies (Figure 2D). Remarkably, 17 human islet eQTLs that colocalized with T2D GWAS signals were not associated with gene expression in any GTEx tissue, including whole pancreas, which emphasizes the fact that pancreas cannot be used as proxy for pancreatic islets and vice-versa.

We compared our findings with those obtained in the InsPIRE islet eQTL study that comprised 420 samples ^23^, of which 207 were also included in our study. We observed that 18 (34%) of the 53 eQTLs that colocalized with T2D GWAS signals were also identified in InsPIRE study (Suppl. Table 4). The improved power in our study obtained by the use of integrative approaches, such as combined reference panels genotype imputation and meta-analysis allowed us to detect lower MAF eQTL signals (10.4% with <5% MAF), representing a 7-fold increment of low frequency eQTL variants compared to this previous large islet eQTL study. Importantly, the meta-analyses also allow us to compare the heterogeneity of the associations between cohorts and filter out signals that are not consistent across cohorts, thereby avoiding false positives.

We detected 32 novel T2D colocalizations with low MAF variants, including variants associated with expression of *CCND2, RMST*, and *CPLX1*. The variant rs76895963 (MAF 0.02) that upregulates *CCND2*, halves the risk of T2D ^37^ and is potentially implicated in the perinatal development of human β-cells ^59^. While the posterior probability of the colocalization was below the threshold of 0.8, the SNP had a clear eQTL with the gene, and a convincing colocalization (see Locus Compare plots, Figure 3G). The variant rs77864822 (MAF=0.07) upregulates *RMST* expression and decreases T2D risk. *RMST* (rhabdomyosarcoma 2 associated transcript) is a reportedly neuron-specific long noncoding RNA involved in neurogenesis ^60^; it is well expressed in human islet cells ^61^ but its function in β-cells is unknown. The variant rs1531583, with the minor T allele associated with increased T2D risk ^5^, upregulates *CPLX1*, encoding complexin-1, again a reportedly neuron-specific gene. Complexin-1 plays a role in Ca^2+^ dependent insulin exocytosis in rodent β-cells, although it is intriguing that both *CPLX1* silencing and overexpression impaired insulin secretion ^62^. GWAS often report as a target the gene closest to the variant, in this case *PCGF3*, for which eQTLs exist in many GTEx tissues. Notably, rs1531583 lies in an intronic region of *PCGF3*, and is an eQTL for this gene in several GTEx tissues. However, we demonstrate here that it is specifically associated with *CPLX1* expression in human islets and not with *PCGF3*, challenging the hypothesis that the closest gene is often the most likely target gene (Figure 3A-E).

The imputation with four reference panels allowed us to analyze different sources of genetic variation, including indels and structural variants. In our study, 12.6% of the eQTL are indels. This stresses the fact that indels are a significant part of the genetic background influencing RNA expression. Unfortunately, the largest available T2D GWAS dataset ^5^ did not consider indels, and so we could not include them in our colocalization analysis. In the near future, this approach could be used to fine-map the contribution to disease risk of indels and structural variants.

Capitalizing on this valuable pancreatic islet resource, we also analyzed *cis*-regulation via ASE for the first time. We developed a novel method named cASE, which combines ASE across samples, maximizing the power to detect variants associated with ASE. We identified variants associated with allelic imbalanced expression while not changing the overall gene expression, and thus undetectable by eQTL. We extended the cASE results in colocalization analysis and identified 14 T2D colocalizations. While 6 of them were detected in the eQTL/T2D GWAS colocalization, 8 were novel signals, including *WFS1, SLC30A8, KCNJ11, TSPAN8, C18orf8* and *CALR*. For these, the lead SNP causes allelic imbalance but no overall gene expression change. These findings suggest that a subset of regulatory genetic variants confer disease risk by causing imbalance in allelic expression of their target genes, a novel mechanism for which knowledge is lacking. A particular locus of interest was the colocalization for common variant rs3802177 associated with *SLC30A8*. rs3802177 is in strong linkage disequilibrium with rs13266634 T2D associated variant, widely discussed in the literature ^63–66^. In our study both variants had nearly identical *p-values* (*p*=2.9×10^-14^ for rs3802177 and *p*=3.3×10^-14^ for rs13266634), showing that any or both of those SNPs could induce allelic imbalance. Rare loss-of-function variants in *SLC30A8* strongly reduce T2D risk ^67^ by enhancing insulin secretion ^68^. However, the direction of effect of the common coding variants is not known. Our cASE results suggest that imbalanced expression towards the rs13266634-T allele is protective for T2D. Since *SLC30A8* loss-of-function decreases risk, these results suggest that the rs13266634-T allele may cause reduced *SLC30A8* function.

In summary, we generated the largest to date expression regulatory variation resource in human pancreatic islets, a tissue with a central pathogenic role in most if not all types of diabetes. All these results are available through the TIGER web portal, which constitutes a user-friendly visualization tool that facilitates the exploration of the datasets, democratizing human islet genomic information to all islet researchers and clinicians.

We expect that this resource, in combination with the growing number of large-scale genetic and functional studies will represent a critical step forward towards understanding the molecular underpinnings of complex diseases that impact pancreatic islet biology and provide a path for the identification of novel and personalized drug targets.

## Supporting information

Appendix MAGIC Authors

## Data and code availability

The eQTL and cASE results are available for browsing at TIGER (http://tiger.bsc.es) and the full summary statistics will also be available for download upon publication.

The cASE code is available through https://github.com/imoran-BSC/TIGER_cASE.

Source data used for this study supporting all findings are available within the article and its Supplementary Information files or from the appropriate repositories. Already published genotype, sequence, methylation and expression data was obtained from the European Genome-phenome Archive (EGA; https://www.ebi.ac.uk/ega/) under the following accession numbers: EGAD00001001601; EGAD00001003946; EGAD00001003947 and Gene Expression Omnibus (GEO; https://www.ncbi.nlm.nih.gov/geo/) under the following accession numbers: GSE50244; GSE76896; GSE53949; GSE35296. Samples used to generate Islet regulome annotations, ChIP-seq and ATAC-seq were taken from EGA repository, under the accession numbers: EGAD00001005203, EGAD00001005202, EGAD00001005204, EGAD00001005201 and their corresponding processed files are available through https://www.crg.eu/es/programmes-groups/ferrer-lab#datasets. New genotype, RNA-sequencing and associated metadata from Pisa and CRG samples are being deposited in EGA (EGA number pending).

## Acknowledgments

This work has been supported by the European Union’s Horizon 2020 research and innovation program T2Dsystems under grant agreement No 667191. L.Alonso was supported by the grant BES-2017-081635 of Severo Ochoa Program, awarded by the Spanish Government. I. Moran was supported by the FJCI-2017-31878 Juan de la Cierva grant, awarded by the Spanish Government. Work in the M.Cnop and D.Eizirik labs was further supported by the Fonds National de la Recherche Scientifique (FNRS), the Brussels Region Innoviris project DiaType and the Walloon Region SPW-EER Win2Wal project BetaSource, Belgium. Eizirik is also supported by a grant from the Welbio–FNRS (Fonds National de la Recherche Scientifique), Belgium, and start-up funds from the Indiana Biosciences Research Institute (IBRI), USA. J.M.Mercader is supported by American Diabetes Association Innovative and Clinical Translational Award 1-19-ICTS-068. J.Chen is supported by an Expanding excellence in England award from Research England. H.Mulder, J.L.S.Esguerra and L.Eliasson are supported by the Swedish Strategic Research Foundation (IRC15-0067).A.L.Gloyn is a Wellcome Trust Senior Fellow in Basic Biomedical Science. This work was funded in Oxford & Stanford by the Wellcome Trust (095101 [ALG], 200837 [A.L.G.], 106130 [A.L.G], 203141 (A.L.G.] and NIH (U01-DK105535; U01-DK085545) [M.I.M., A.L.G.]. The research was funded by the National Institute for Health Research (NIHR) Oxford Biomedical Research Centre (BRC) [A.L.G.]. I.Miguel-Escalada was supported by EFDS/Novo Nordisk Rising Star Programme. Work in J.F. lab was supported by Imperial College London Research Computing Service, NIHR Imperial Biomedical Research Centre (BRC), and CRG genomics facility, and grants from Ministerio de Ciencia e Innovación (BFU2014-54284-R, RTI2018-095666-B-I00), Medical Research Council (MR/L02036X/1), Wellcome Trust Senior Investigator Award (WT101033), European Research Council Advanced Grant (789055). The views expressed are those of the author(s) and not necessarily those of the NHS, the NIHR or the Department of Health.

The technical support group from the Barcelona Supercomputing Center is gratefully acknowledged. Finally, we thank the entire Computational Genomics group at the BSC for their helpful discussions and valuable comments on the manuscript. We also acknowledge Cristian Opi for designing the T2DSystems logo and Laia Codó for the technical support with the website allocation, Isabelle Millard and Anyishaï Musuaya from the ULB Center for Diabetes Research for excellent technical and experimental support.

## Data Sources

The database generated in this project has made use from the following list of publicly available resources: The Genotype Tissue Expression (GTEx) Project was supported by the Common Fund of the Office of the Director of the National Institutes of Health, and by NCI, NHGRI, NHLBI, NIDA, NIMH, and NINDS. The data used for the analyses described in this manuscript were obtained from the GTEx Portal [GTEx Analysis V7 - Transcript TPMs] on 04/09/19. FastDMA proble full annotation: Wu D, Gu J, Zhang MQ (2013) FastDMA: An Infinium HumanMethylation450 Beadchip Analyzer. PLoS ONE 8(9): e74275. ENCODE (2012-2016) Open Chromatine Dnase: The ENCODE Project Consortium. An integrated encyclopedia of DNA elements in the human genome, Nature (2012). ENCODE Data Coordination Center (DCC). The ENCODE Consortium and the ENCODE production laboratory(s) generating the datasets. Gene Ontology: Ashburner et al. Gene ontology: tool for the unification of biology (2000) Nat Genet 25(1):25-9. Online at Nature Genetics. GO Consortium, Nucleic Acids Res., 2017. DIAGRAM 1000G GWAS meta-analysis Stage 1 Summary statistics, Trans-ethnic T2D GWAS meta-analysis and DIAMANTE T2D GWAS meta-analysis. DIAGRAM Consortium. Reactome Pathway database: Reactome https://reactome.org/download-data/ (Jul 2017). DisGeNET, May 2017. Janet Piñero, Àlex Bravo, Núria Queralt-Rosinach, Alba Gutiérrez-Sacristán, Jordi Deu-Pons, Emilio Centeno, Javier García-García, Ferran Sanz, and Laura I. Furlong. DisGeNET: a comprehensive platform integrating information on human disease-associated genes and variants. Nucl. Acids Res. (2016). doi:10.1093/nar/gkw943. GWAS Catalog version 1.0 release 2020-12-02. MacArthur J, Bowler E, Cerezo M, Gil L, Hall P, Hastings E, Junkins H, McMahon A, Milano A, Morales J, Pendlington Z, Welter D, Burdett T, Hindorff L, Flicek P, Cunningham F, and Parkinson H. The new NHGRI-EBI Catalog of published genome-wide association studies (GWAS Catalog). Nucleic Acids Research, 2017, Vol. 45 (Database issue): D896-D901. Ensembl Variant Effect Predictor version 87.27. McLaren W, Gil L, Hunt SE, Riat HS, Ritchie GR, Thormann A, Flicek P, Cunningham F. The Ensembl Variant Effect Predictor. Genome Biology Jun 6;17(1):122. (2016). doi:10.1186/s13059-016-0974-4. RefSeq BUILD.37.3: O’Leary NA, Wright MW, Brister JR, Ciufo S, Haddad D, McVeigh R, Rajput B, Robbertse B, Smith-White B, Ako-Adjei D, Astashyn A, Badretdin A, Bao Y, Blinkova O, Brover V, Chetvernin V, Choi J, Cox E, Ermolaeva O, Farrell CM, Goldfarb T, Gupta T, Haft D, Hatcher E, Hlavina W, Joardar VS, Kodali VK, Li W, Maglott D, Masterson P, McGarvey KM, Murphy MR, O’Neill K, Pujar S, Rangwala SH, Rausch D, Riddick LD, Schoch C, Shkeda A, Storz SS, Sun H, Thibaud-Nissen F, Tolstoy I, Tully RE, Vatsan AR, Wallin C, Webb D, Wu W, Landrum MJ, Kimchi A, Tatusova T, DiCuccio M, Kitts P, Murphy TD, Pruitt KD. Reference sequence (RefSeq) database at NCBI: current status, taxonomic expansion, and functional annotation. Nucleic Acids Res. 2016 Jan 4;44(D1):D733-45. Gencode v23 lift 37 annotation: Frankish A, et al (2018) GENCODE reference annotation for the human and mouse genomes. Nucleic Acids Res 2018: Oct24. doi:10.1093/nar/gky955. gnomAD version 2.0.2: The authors would like to thank the Genome Aggregation Database (gnomAD) and the groups that provided exome and genome variant data to this resource. A full list of contributing groups can be found at http://gnomad.broadinstitute.org/about. Data on glycaemic traits have been contributed by MAGIC investigators (www.magicinvestigators.org.). Members of MAGIC are provided in Appendix S1.

## Author Contribution

L.A., A.P., I.M., J.F., J.M.M., M.C. and D.T. conceived and planned the main analyses. J.F. provided unpublished allelic ChIP-seq and RNA-seq datasets, and supervised cASE, which was developed and implemented by I.M. during his PhD in IDIBAPS and Imperial College London. I.M. further applied cASE in the TIGER dataset with collaboration of L.A., M.G-M., S.B-G., M.P., R.A. and J.M.M. A.P. performed eQTL and colocalization analyses with collaboration of L.A., M.G-M., S.B-G., M.D., R.A. and J.M.M. L.A. developed the TIGER portal with collaboration of R.R and J.M.M. and performed expression analysis with collaboration of I.M. and A.P. and J.M.M. I.M., A.P., L.A., J.M.M., D.T. and M.C. wrote and edited the manuscript. G.A. and I.M-E. contributed with islet regulatory data and analysis. I.M., S.B-G. and J.F. contributed with Imperial and CRG data and analysis. J.L.S.E., L.E., H.M. and L.G. contributed with Lund data and analysis. J-V.T., D.L.E. and M.C. contributed with ULB data and analysis. M.S., L.M. and P.M. contributed with Pisa data and analysis. M.S., L.M., P.M. contributed with Pisa islet samples. J.L.S.E. contributed with Pisa sample sequencing. V.N. contributed with Pisa sample genotyping. J.M.T., V.N. and A.L.G. contributed with Oxford data and analysis and the genotyping of Pisa samples. X.G-H prepared chromatin immuno-precipitaton, RNA and DNA samples and managed CRG data generation. A.L.G., J.L.S.E., P.M., D.L.E, J.F., J.M.M., M.C. and D.T. provided guidance in the design and during the development of the project. D.L.E., M.C. and D.T. worked on the creation of TIGER. J.C. and MAGIC contributed with MAGIC data and analysis. J.M.M., M.C. and D.T. supervised the study.

## Notes

### Competing Interest Statement

The authors have declared no competing interest.

