## Supplemental Methods for "TIGER: The gene expression regulatory variation landscape of human pancreatic islets"

**Online Methods**

### Islet sample collection and genotyping

TIGER data consists of 514 RNA-seq and 485 genotyped array data of deidentified cadaveric human pancreatic islet samples from five research centers: 1) Centre for Genomic Regulation, 2) Lund University Diabetes Centre, 3) University of Oxford/University of Alberta, 4) Department of Endocrinology and Metabolism, University of Pisa and 5) ULB Center for Diabetes Research, Universite Libre de Bruxelles (Suppl. Table S1).

**Centre for Genomic Regulation (CRG).** The DNA of 127 CRG samples was isolated, sequenced and genotyped using Illumina's Human OmniExpress 12 v1 and 2.5-8 v1.1 chips, as described in Miguel-Escalada I. et al 2019 ^1^. Genotype array was done in 125 samples with Illumina’s Genome Studio software providing information on a total of 624k SNPs.

**Lund University Diabetes Centre (Lund).** The DNA of 89 Lund samples from cadaver donors of European ancestry provided by the Nordic Islet Transplantation Programme was isolated as described in Fadista J. et al. 2014 ^2^. The samples were genotyped using Illumina’s HumanOmniExpress 12v1 C chips passing standard quality control metrics providing information on a total of 609k SNPs.

**University of Oxford/University of Alberta (Oxford).** The DNA of 118 Oxford samples was isolated from either spleen or the exocrine fraction of the islet isolation using the Tissue DNA Purification Kit. When no other tissue was available, DNA was extracted from human islets using the Trizol fraction remaining after extraction of RNA as described in van de Bunt M. et al 2015 ^3^. The samples were genotyped using Illumina’s Human Omni 2.5 exome array following the Illumina Infinium protocol providing information on a total of 2.5M SNPs.

**University of Pisa (Pisa).** The DNA of 154 Pisa samples was isolated according to previously described in Marselli et al. 2020 ^4^ and sequenced. Genotype calling was done in 153 samples with Illumina’s Human Omni 2.5 exome array providing information on a total of 2.6M SNPs.

**ULB Center for Diabetes Research (ULB).** The 43 ULB samples were isolated in Pisa using collagenase digestion and density gradient purification from beating-heart organ donors with no medical history of diabetes or metabolic disorders. Following islet shipment to Brussels, mRNA was extracted and processed following the RNeasy Qiagen protocol as described in Cnop M. et al. 2014 ^5^.

### Genotyping quality control

PLINK v1.9 ^6^ was used to do standard quality control of the genotype data, at the variant and sample level. At the variant level, we discarded rare variants (Minor Allele Frequency MAF<0.01) and applied Hardy-Weinberg equilibrium test filtering (*p*<=1×10^-6^) ^7,8^. Further, we filtered the variants below a missingness threshold of 0.05. At the sample level, we discarded samples presenting a gender discordance between the reported gender in the metadata and the genetic sex, as well as the subjects with at least a 3rd degree of relatedness, those below a missingness threshold of 0.02 and, finally, individuals not clustering within the 4 standard deviations of the first four principal components from the multidimensional scale analysis. The ancestry of the individuals was assessed by principal components analysis comparisons with phase3 1,000 Genomes Project populations ^9^.

After QC this resulted in a total of: 1) 103 individuals, 559,083 SNPs in the CRG cohort, 2) 88 individuals, 596,273 SNPs in the Lund cohort, 3) 102 individuals, 1,487,651 SNPs in the Oxford cohort and 4) 144 individuals, 1,542,765 SNPs in the Pisa cohort.

### Genotype phasing and imputation

The autosomal genotypes were phased with Eaglev3 ^10,11^ using the Human Reference Consortium Project reference panel ^12^. The X chromosome was phased without reference panel with SHAPEIT ^13^ (Suppl. Figure S11). GUIDANCE ^14^ integrating IMPUTE2 ^15^ was used for imputation, using 4 reference panels: the 1000 Genomes Project phase 3 ^9^, the Genome of the Netherlands Project ^16^, the Haplotype Reference Consortium Project ^12^ and the UK10K Project ^17^, with an IMPUTE2 info score threshold of ≥ 0.7. This resulted in a total of 13.7-16.3M SNPs for each cohort separately, that were merged considering the best info score obtained across all panels, resulting in 22,983,795 genotyped and imputed genetic variants with MAF>0.001.

### RNA-seq read mapping

RNA from 514 human donor islet samples was isolated and purified, and was used to construct RNA-seq libraries. These RNA-seq assays generated a total of >72 billion pair-ended fragments of 75, 76, 100, 101, 125 bp read lengths.

To perform eQTL analysis, we aligned all samples against the transcriptome reference gencode.v23lift37^18^ with STAR v2.4.0 ^19^, using

--paired-end –p 8

An alternative mapping strategy was used for RNA-seq read mapping to be used for cASE. Given that the standard reference genome contains only one allele in polymorphic sites, standard RNA-seq read mapping can produce reference-biased alignments, leading to false positives in the study of ASE. To align RNA-seq datasets in an allele unbiased manner, two modified reference genomes were built, defined as a ‘masked’ and an ‘enhanced’ genome. The ‘masked' reference genome was built by substituting with an ‘N’ the nucleotide position of each common SNP in dbSNP142 ^20^ (MAF >1%), using the vcf2diploid.jar ^21^ tool. To construct the ‘enhanced' reference genome, we modified the scripts developed by Satya et al. ^22^ to accommodate RNA-seq reads, which added artificial contigs to the reference genome containing all possible SNP allele combinations. For this step, we used the subset of 4M common SNPs located within gene coordinates in the Ensembl ^23^, RefSeq ^24^ and UCSC ^25^ annotations, or within previously identified human islet lncRNAs ^26^ (Suppl. Figure S6).

STAR v2.2.0 ^19^ was used to align the RNA-seq datasets against the masked genome, using

--outFilterMultimapNmax 1 --outFilterMismatchNmax 10

--outSAMstrandField intronMotif --outSAMattributes All

in order to allow up to 10% of nucleotide mismatches, suppress multimapped reads, and make the output compatible with downstream software. Bowtie v2.0.5 ^27^ was used to align the RNA-seq data against the enhanced genome, using

--n-ceil L,0,0.03 --score-min C,-14,0 -N 1 -X 50000

to allow up to 3 nucleotide mismatches evenly distributed within the read, and long range read pairs. Bowtie2 ^28^ was chosen because it does not map the RNA-seq spliced reads, (only the reference allele-containing spliced sequences were present in the enhanced genome) which prevents the generation of allelic alignment bias.

After mapping the RNA-seq datasets to the two modified reference genomes, the outputs of both alignments were combined into one non-redundant set of reads, using the read merging C++ scripts available in our github repository (https://github.com/imoran-BSC/TIGER_cASE , scripts 02 and 03). Reads that aligned to the same genomic positions by both methods were kept, as well as reads mapped only by one of the two methods. In addition, all reads that mapped partially to intronic regions were discarded. The resulting set of reads was named ‘unbiased alignment’ (Suppl. Figure S6A). This method successfully eliminated alignment bias in heterozygous positions (Suppl. Figure S6B), and mapped 86.2% of all RNA-seq reads. When comparing this alignment with one using the standard reference genome and STAR v2.2.0 using a subset of the samples, we recovered an extra 8.5% more reads using the unbiased alignment method (Suppl. Figure S6C).

### Sample concordance verification between genotype and gene expression

To avoid mislabeled samples leading to mismatching errors between genotype-phenotype samples, and to discard samples with poor quality or possible contamination, we used verifyBamID v1.1.3 ^29^ with “--best”, applied to the RNA-seq alignments sorted and indexed with samtools v1.1 ^30^, and comparing with their genotypes. After these steps, 404 samples with good quality genotype and RNA-seq data and concordance remained for further analysis.

### eQTL analysis

The cis-eQTL analysis of 404 human pancreatic islets for which both RNA-seq and genotyping data remained after QC was performed by cohort with fastQTL v2.0 tool ^31^. The analysis was run for regions one million base pairs up- or downstream from the transcription start site of each gene using *gencode.v23lift37* ^18^ version. For each cohort, we corrected for known covariates (age, sex and BMI), 7 genomic ancestry principal components, and 15 PEER v1.3 ^32^ factors in order to account for hidden confounding factors. For the X chromosome, we used 5 PEER factors and 4 genomic ancestry principal components and the *cis*-eQTL analysis was performed stratified by sex and combined. The full command for *fastQTL* is

fastQTL --log 'chr1.log' --vcf 'chr1.bcf' --bed 'rsem.bed' -C ‘covariates.tsv’ --threshold '0.01' --out 'chr1.fastQTL.gz'

Age and BMI missing metadata was imputed using the cohort mean.

The by-cohort fastQTL ^31^ results were then meta-analyzed with METAL ^33^ using the sample size strategy and computing heterogeneity. For the X chromosome, the meta-analysis was run over the 4 cohorts for both sexes together and over the 8 eQTL analysis (4 cohorts, 2 sexes). The full configuration files for METAL are given by:

SEPARATOR WHITESPACE

MARKER ensg.snp

ALLELE a0 a1

EFFECT slope

PVALUE pval

WEIGHT N

PROCESS cohort_CRG

PROCESS cohort_OXFORD

PROCESS cohort_LUND

PROCESS cohort_PISA

OUTFILE metal .tsv

ANALYZE HETEROGENEITY

QUIT

### Identifying variant regulatory enrichments using GREGOR

To test the eQTL and cASE variants for enrichment in islet regulatory overlaps, we used the Genomic Regulatory Elements and Gwas Overlap algoRithm (GREGOR) ^34^, designed to calculate such enrichment while controlling for linkage-disequilibrium between variants, MAF and distance to nearest gene. We used the 1% and 5% FDR set of significant eQTL variants, after selecting them by linkage disequilibrium <0.2 using PLINKv1.9 ^6^ with “--indep-pairwise 100k 5 0.2”. We tested enrichment against a set of human islet regulatory regions, including gene promoters, enhancers and open-chromatin derived from ChIP-seq experiments in human islets (Figure 2C, Suppl. Figure S2, Suppl. Figure S12) ^1^. Specifically, we used an R^2^ threshold of 0.99, a window size of 1,000,000, a min_neighbor_num of 500, and European (EUR) as the population.

**Comparison of TIGER eQTLs with the GTEx and InsPIRE datasets**

To assess the degree of concordance between the TIGER significant eQTLs and those reported in the GTEx v8 dataset ^35^, we searched for exact variant-target gene matches among the dataset of significant eQTLs in all 54 GTEx tissues. To analyze the overlap of eQTLs with low-frequency variants, we repeated the analysis, but first filtered the TIGER and GTEx eQTLs to include only those with variants with a MAF < 0.05 in the EUR population of the 1000 genomes phase-3 dataset ^9^.

To obtain a relevant comparison with the InsPIRE ^36^ dataset, we first applied the same multiple-testing correction method used in this study to the full nominal p-values of the InsPIRE dataset. The Benjamini-Hochberg corrections for 1 and 5% FDR resulted in the nominal *p*-value thresholds of *p*=8.55×10^-5^ and *p*=6.2×10^-4^, corresponding to 974,435 and 1,408,891 significant eQTLs. Two eQTLs were considered significant by both methods if they were detected at <5% FDR in both studies, and had an exact match in both variant and target gene. The low-frequency variant eQTLs were determined as described above.

### Colocalization analysis

COLOC 4.0 ^37^ R package was used for the colocalization analysis of cis-eQTL and T2D GWAS. We used the coloc.abf method which implements a variation of the Approximate Bayes Factor computations ^38^. The coloc.abf function was called with two R lists, one for the eQTL and one for the GWAS:

list(pvalues=…, N=…, MAF=…, snp=…, type="quant")

with a vector of *p*-values, N the sample size, MAF the minor allele frequency and snp the rsid of the variant.

In order to select regions for colocalization analyses, we selected genes associated with at least one significant eQTL SNP which had been previously reported as a GWAS lead variant ^39–41^. The significant eQTL SNPs were determined based on a 0.05 threshold Benjamini-Hochberg FDR ^42^. Similarly, we used the *p*-values of the cASE analysis to perform colocalization, considering loci with an at least 5% FDR significant signal. The colocalization was run over regions ranging from one million base pairs downstream to one million upstream from the cis-regulatory target gene transcription start site.

The colocalization plots were generated by the locuscompare R package v1.0.0 ^43^ (Suppl. Figure S5, Suppl. Figure S9).

### Generation of an unbiased set of ASE reporter variants

To identify loci under mappability related allelic biases, a C++ script available in the github repository (https://github.com/imoran-BSC/TIGER_cASE, script 01) was used to generate all possible reads containing both alleles of all possible reporter SNPs. A splice junction database was created using the Ensembl ^23^, RefSeq ^24^, UCSC ^25^ and human islet lncRNA ^26^ gene annotations, to take splice junctions into account.

The resulting dataset, consisting of 240M artificial reads, was aligned using the unbiased mapping strategy described above, and the allelic ratios (i.e., the percentage of reference-allele carrying reads) were quantified. Since the same number of reads were purposely generated carrying both alleles, any observed allelic imbalance would derive exclusively from mapping biases. SNPs whose allelic ratio was not between 49-51% were blacklisted. Additionally, all SNPs located within 100 bps of a common or low-frequency indel present in dbSNP142 ^20^ were also blacklisted.

The remaining curated set of 3.97M SNPs were used as bona-fide SNPs for reporting ASE.

### Identification of ASE

The number of reads containing the reference and alternate alleles RNA-seq reads overlapping each reporter SNP were quantified using the mpileup command of samtools v1.1 ^30^, with the flags “-A -B -d 20000”, and the ComputePileupFreqs.pl script ^22^. Sample-specific ASE was assessed calculating the allelic ratio, i.e., the fraction of reads containing the reference allele over the total number of reads. We selected the set of SNPs with at least 3 heterozygous samples with ≥15 RNA-seq reads (of which ≥10 non-clonal), resulting in a set of >170k informative reporter SNPs.

A binomial test ^44^ was used to assess the significance of ASE for all reporter SNPs, using the number of reads carrying the reference and alternate alleles. To account for any possible remaining alignment bias in the datasets, the median allelic ratio for each possible bi-allelic SNP (AC, AG, AT, CG, CT, GT) across the genome was calculated and used as null, instead of the theoretical 50%. Similarly, the allelic ratios were proportionally adjusted using the sample and nucleotide-pair specific median value.

The resulting *p*-values were used to calculate a sample-specific 1% and 5% FDR Benjamini-Hochberg ^42^ thresholds, to correct for multiple testing.

### Assessing cASE using Stouffer's Z-score

To assess cASE in a given heterozygous variant in many independent samples, the Stouffer's Z-score ^45^ method was used. This method combines independently obtained p-values into a Z statistic, which increases in absolute value with significance. The method allows for weighting of independent *p*-values and, additionally, it accounts for a positive or negative direction in the magnitude associated with the p-values. Thus, this method allows to differentiate between significant reference and alternate reporter variants, as well as providing a way to account for the variance inherent to differing numbers of informative RNA-seq reads in each reporter.

For each reporter, a Z-score was calculated as follows:

$$Z=\frac{\sum w_{i}Z_{i}}{\sqrt{\sum w_{i}^{2}}}$$

where $w_{i}$ was the total read coverage of sample *i*, and $Z_{i}$ was the transformed binomial p-value $p_{i}$:

$$Z_{i}=\pm\theta^{-1}\left( 1-\frac{p_{i}}{2} \right)$$

where the sign was positive if the value of the allelic ratio was >50%, zero if exactly 50%, and negative otherwise, and $\theta^{-1}$ was the inverse of the standard normal cumulative distribution function, calculated using the qnorm function in R. A threshold of 10^-15^ was imposed as the minimum possible binomial p-value, in order to prevent single events with very significant *p*-values from dominating the Z-score value, while still maintaining their relevance. Therefore, Stouffer’s Z-score ^45^ method accounted for consistency in the overall reference or alternate direction of the allelic bias across samples, and considered all p-values into account, regardless of their sample-specific significance.

Z-scores were only calculated if the reporter SNP was heterozygous in 3 or more samples, and only samples with a read coverage of ≥15 RNA-seq reads, of which ≥10 non-clonal, were used in the calculation.

### Assessing the significance of cASE Z-scores

To assess the significance of the obtained Z-scores, we performed 1,000 permutations of the reference/alternate read counts between heterozygous SNPs, and calculated their binomial p-values and resulting control Z-scores (https://github.com/imoran-BSC/TIGER_cASE, script 04). To account for the differences in gene expression, all reporter SNPs were distributed in 5 bins: one containing all SNPs with a median coverage of 0 reads, and 4 more bins containing the remaining SNPs according to their read coverage quartile, and the read counts of heterozygous SNPs were only shuffled within their bins. By permuting only the values of the heterozygous SNPs while keeping the reference and alternate homozygous values invariant, the distribution of the number of samples in heterozygosity for each SNP was kept constant.

The resulting null distribution of Z-scores was therefore attributable only to stochasticity, and so for each empiric Z-score, a p-value was calculated from this null distribution. The Benjamini-Hochberg method ^42^ was then used to obtain *q*-values from these *p*-values and thus correct for multiple testing.

### Regulatory enrichment of cASE significant genes

To calculate these regulatory enrichments, we first generated a null distribution of control genes that were non-significant for cASE but had similar expression levels. First, we separated the cASE significant genes in 4 bins of expression, and randomly selected the same number of non-significant genes of the same expression quartile, 1,000 times. We then calculated, in the 1% and 5% FDR cASE genes and in each of the 1,000 control sets, the proportion of genes that were in the islet-specifically expressed genes list ^1^ (Figure 4C, left). The same procedure was performed to calculate the enrichment for proximity to islet enhancers, by calculating the proportion of genes located at less than 25kb from islet enhancers ^1^. The p-values were obtained by approximating these permuted control distributions as Gaussian distributions and deriving a *p*-value using the pnorm R function.

### Gene ontology analyses and islet-specific expression

Gene ontology terms in the analyses of eQTL and cASE genes were obtained using the PANTHER (Protein ANalysis THrough Evolutionary Relationships) ^46,47^ classification system.

For eQTL, we analyzed all 5% FDR significant genes versus a background list of all genes expressed in islets (Suppl. Figure S4), and the list of TIGER exclusive eQTL genes versus a background of all eQTL genes shared with GTEx (Figure 2F).

For cASE, we studied 5% FDR cASE genes versus a background dataset of all genes for which the calculated cASE was NON-SIGNIFICANT (Figure 4D). The visualization of the syntactic terms was obtained using the REVIGO web tool ^48^.

### Identifying candidate SNPs putatively leading to cASE

We aimed to characterize the set of SNPs putatively causal of cASE (referred to as ‘candidate SNPs’). To that end, we first identified all variant pairs consisting of a cASE-significant reporter and a candidate variant, as long as both were located within the same topologically associating domain (TAD) ^49^, plus a boundary leeway of ±200kbs. Then, we separated the samples using the candidate variant genotype in two groups: those heterozygous (Het), and those homozygous (Hom). Finally, we calculated the reporter Z-score of both sample groups, and selected the candidate variants with significant Z-scores for the Het individuals, which were also non-significant for the Homs (https://github.com/imoran-BSC/TIGER_cASE, script). The underlying hypothesis was that if the candidate variant was homozygous, it was unlikely to be causal.

Putative causal variants were also interrogated for the set of non-cASE significant reporter variants, following the same procedure described above. This produced an additional 1,247 genes that reached cASE significance only after being considered with these putative causal variants.

### Scaling human islet gene expression values to allow comparisons with the GTEx expression datasets in TIGER

The RNA-seq expression of human islet samples was measured with RSEM v1.3.0 ^50^ in 60,261 transcripts from Gencode database (v23lift37 annotation) ^18^ using STAR v2.5.3.a ^19^ and BOWTIE v2.3.2 ^28^ hg19 aligned-reads as follows:

STAR --runMode genomeGenerate --genomeFastaFiles GRCh37.primary_assembly.genome.fa --sjdbGTFfile gencode.v23lift37.annotation.gtf
rsem-prepare-reference --gtf gencode.v23lift37.annotation.gtf --bowtie2 GRCh37.primary_assembly.genome.fa
rsem-calculate-expression --paired-end --star --paired-end -p 8

We obtained measures of raw counts, counts normalized by transcript length (TPM - transcripts per million) and fragment length (FPKM - fragments per kilobase). The batch effects and covariate differences between samples captured in the TPM measures were removed with limma removeBatchEffect function ^51^, using the log10 normalized expression of the genes that were expressed in at least 80% of human islet samples. The results of this normalization were evaluated with Spearman correlation, ensuring that there was a correlation above 0.8 between all the samples independently of the cohort after correction (Suppl. Figure S10).

TPM expression datasets from the 54 tissues available in GTEx ^52^ (20 samples per tissue) were collected, and a decile distribution analysis was performed excluding genes from GTEx samples that miss expression in at least 50% of the samples. Then, TIGER islet expression was scaled to fit these measures according to the following criteria:

(1) Each GTEx decile bin [D_G;i_,D_G;i+1_] has TPM values in [T_G;i_,T_G;i+1_], thus the corresponding decilic straight will be: $y_{G}=\left( T_{G;i+1}-T_{G;i} \right)x+T_{G;i}$.
(2) Each pancreatic islet decile bin [D_G;i_,D_G;i+1_] has TPM values in [T_PI;i_,T_PI;i+1_], thus the corresponding decilic straight will be: $y_{PI}=\left( T_{PI;i+1}-T_{PI;i} \right)x+T_{PI;i}$.

From equation (2) one can derive: $x=\frac{y_{PI}-T_{PI;i}}{T_{PI;i+1}-T_{PI;i}}$ (3) thus, allowing the relation between the TPM pancreatic islet values $y_{PI}$ and the TPM GTEx values $y_{G}$ by replacing (3) in (1): $y_{G}=\left( \frac{T_{G;i+1}-T_{G;i}}{T_{PI;i+1}-T_{PI;i}} \right)y_{PI}-T_{PI;i}\left( \frac{T_{G;i+1}-T_{G;i}}{T_{PI;i+1}-T_{PI;i}} \right)+T_{G;i}$ the scaling factor.

### TIGER web portal development

The TIGER web portal (http://tiger.bsc.es) is the comprehensive integration in an ElasticSearch v1.4.4 database of a) T2D GWAS variants identified in 70KforT2D ^41^, diagram DIAMANTE ^40^, diagram Trans-ethnic ^53^, diagram 1000G ^54^ T2D meta-analyses or included in the GWAS Catalog v1 release 2020-12-02 ^55^, b) variant annotation and characterization through Variant Effect Predictor v87.27 ^56^ and Gnomad v2.0.2 ^57^, c) epigenomic marks from islet DNA-methylation sites ^58,59^, chromatin accessibility ^60–62^ and CHiP-seq profiles ^1^, d) annotation from Gene Ontology ^63,64^, lncRNAs ^26^ and islet regulome ^1,65^ in a publicly available platform. Genes are referenced to Gencode annotation v23 lift 37 ^18^ and RefSeq BUILD.37.3 ^24^ and enriched with DisGeNET ^66^ (May 2017) and Reactome Pathway ^67^ database information. It contains results on gene expression integrating the results of a) gene expression from normalised islet RNA-seq counts, microarrays and the Genotype-Tissue Expression database (GTEx) ^52^, and b) computed eQTL and cASE.

The portal was built upon [[ICGC software codebase](https://github.com/icgc-dcc)], the front-end coded in angular v1.5.7 with embedded biodalliance v1.4.4 genomic browser ^68^, plotly v1.54.1 ^69^ and highcharts libraries and the back-end coded in Java.

### Supplemental Figure Legends:

**Supplementary Figure S1:** Quantile-quantile plot of eQTL meta-analysis *p*-values. A comparison is shown between the expected theoretical normal quantiles under the null hypothesis and the empirical quantiles obtained by the eQTL meta-analysis.

**Supplementary Figure S2:** Fold enrichment over controls of significant eQTL variants, in islet regulatory chromatin regions. *P*-values for 1% FDR eQTL enrichments. A) all eQTL variants, B) Top eQTL variants: common (>=5% MAF) and low-frequency variants (1%<MAF<5%).

**Supplementary Figure S3:** Comparison between TIGER and InsPIRE eQTL results. A) Venn diagram of overlap of the 5% FDR significant eQTL Genes. B) As in A) but restricted to eQTLs genes with low-frequency variants (<5% MAF). C) Venn diagram of the overlap with 5% FDR significant eQTLs in TIGER and in InsPIRE. D) As in C) but restricted to the eQTLs with low-frequency variants (<5% MAF).

**Supplementary Figure S4:** Gene ontology of the eQTL genes. A) Gene ontology analysis of the genes with significant eQTLs compared with a background of all genes expressed in human islets, visualized with the Gene Ontology enRIchment anaLysis and visuaLizAtion (GOrilla) web tool. B) Same information visualized using Revigo.

**Supplementary Figure S5:** Co-localization plots of eQTL signals. LocusCompare plots depicting all significant colocalizations between eQTL and T2D GWAS analyses. The lead variant is represented by a purple diamond. The linkage disequilibrium between the lead variant and the other variants is given as the square of the correlation coefficient r² and is indicated in a color scale. The -log10(p-values) for each variant — which are located in a region of one mega-base pair up- and downstream from the gene transcription start site — are depicted in three panels: (left) p-values of eQTL as x-axis and GWAS as y-axis, (bottom right) p-values of GWAS in the gene region and (top right) p-values of eQTL in the gene region. The title shows the gene name; MAF: minor allele frequency; PP.H4.abf: Posterior probability of colocalization; SNP.PP.H4: posterior probability of lead variant being the associated causal variant.

**Supplementary Figure S6:** Overview of the RNA-seq alignment used in cASE analysis. A) Decision tree of the RNA-seq reads to keep or discard, after alignment using both a masked and an enhanced reference genome. B) Mean allelic ratio resulting from the ‘unbiased alignment’ method compared versus an alignment using a standard reference genome. C) As in B but showing the percentage of raw reads aligned. D) Allelic ratio of reporter variants separated by their genotypes, across the four cohorts. E) Mean and median allelic ratio values for each of the cohorts.

**Supplementary Figure S7:** An example of a gene with multiple cASE reporter variants. A) *IAPP* has 7 distinct significant cASE reporters, all located in its last exon. B) The Z-score plots of the top 6 reporters show a consistent imbalance towards the reference allele across individuals.

**Supplementary Figure S8:** Examples of candidate cis-regulatory variants of genes without significant cASE reporters. Left, the Z-score plots for 6 genes, separating samples according to the genotype of the candidate variant, Heterozygous (green diamonds) or Homozygous (yellow). Right, table showing the reporter and candidate variant Z-scores and their p-values.

**Supplementary Figure S9:** Co-localization plots of cASE signals. LocusCompare plots depicting all significant colocalizations between cASE and T2D GWAS analyses. The lead variant is represented by a purple diamond. The linkage disequilibrium between the lead variant and the other variants is given as the square of the correlation coefficient r² and is indicated in a color scale. The -log10(*p*-values) for each variant — which are located in a region of one mega-base pair up- and downstream from the gene transcription start site — are depicted in three panels: (left) *p*-values of cASE as x-axis and GWAS as y-axis, (bottom right) *p*-values of GWAS in the gene region and (top right) p-values of cASE in the gene region. The title shows the gene name; MAF: the minor allele frequency; PP.H4.abf: Posterior probability of colocalization; SNP.PP.H4: posterior probability of lead variant being the associated causal variant.

**Supplementary Figure S10:** Gene expression homogenization. Heatmaps with gene expression spearman correlation (1-rho) between each pair of samples before (left) and after (right) batch effects and covariates correction. Sample colors represent the different cohorts included in the study.

**Supplementary Figure S11:** Phasing. Summary of the analysis performed to evaluate the effect of using a reference panel or not in the phasing step. The barplots represent the number of SNPs imputed with good quality (IMPUTE2 info score>=0.7) for each chromosome. The dark blue bars sum the number of variants obtained after imputation with HRC reference panel in the phasing step. The light blue bars sum the number of variants obtained after imputation without a reference panel in the phasing step. Chromosome X was split in males.

**Supplementary Figure S12:** Fold enrichment over controls of significant cASE variants, in islet regulatory chromatin regions. *P*-values for 1% FDR eQTL enrichments. A) all cASE variants, B) Top cASE variants: common (>=5% MAF) and low-frequency variants (1%<MAF<5%).
