## Supplementary figures and images for "TIGER: The gene expression regulatory variation landscape of human pancreatic islets"

### Supplemental Figure 1

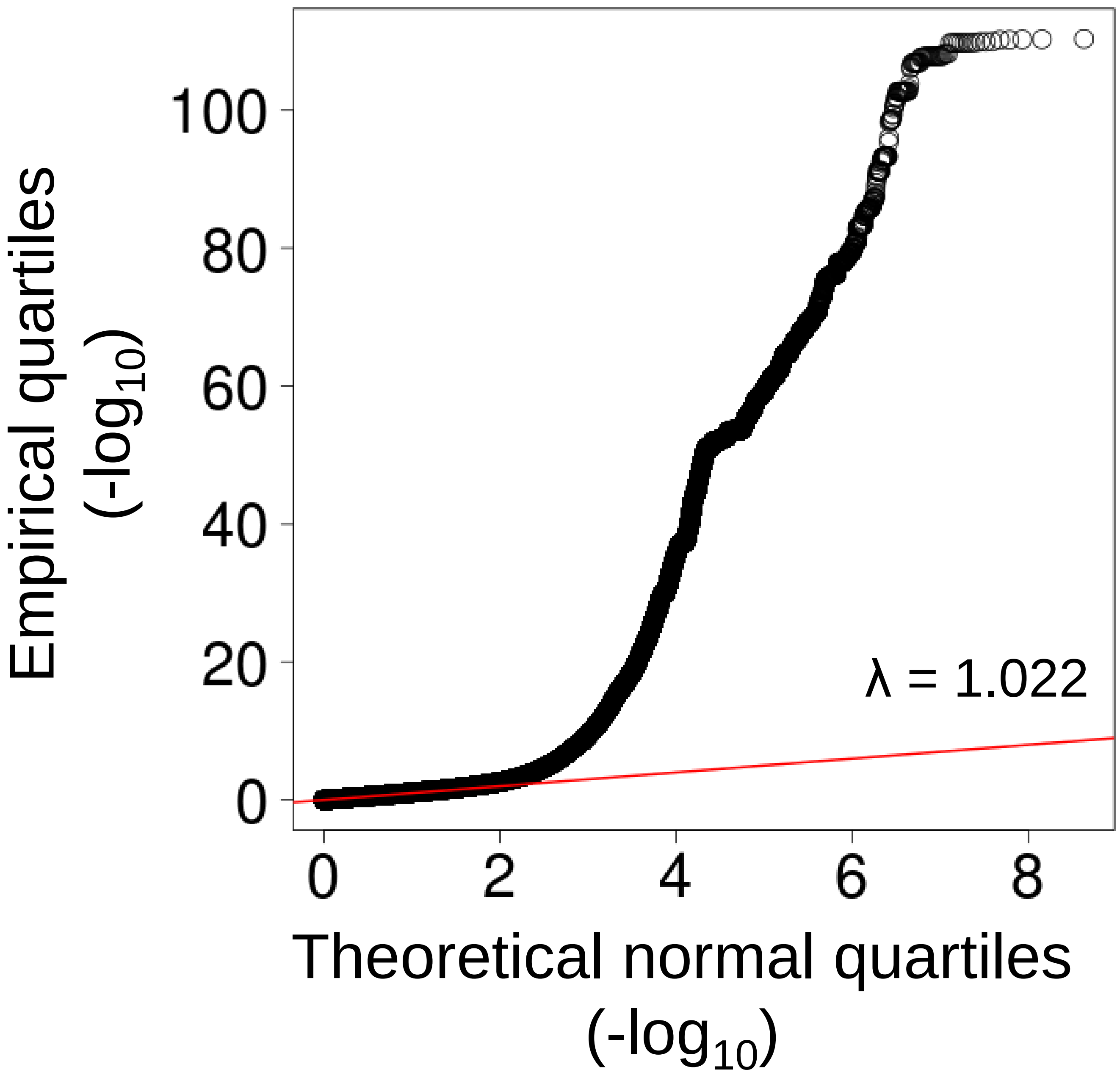

### Supplemental Figure 2

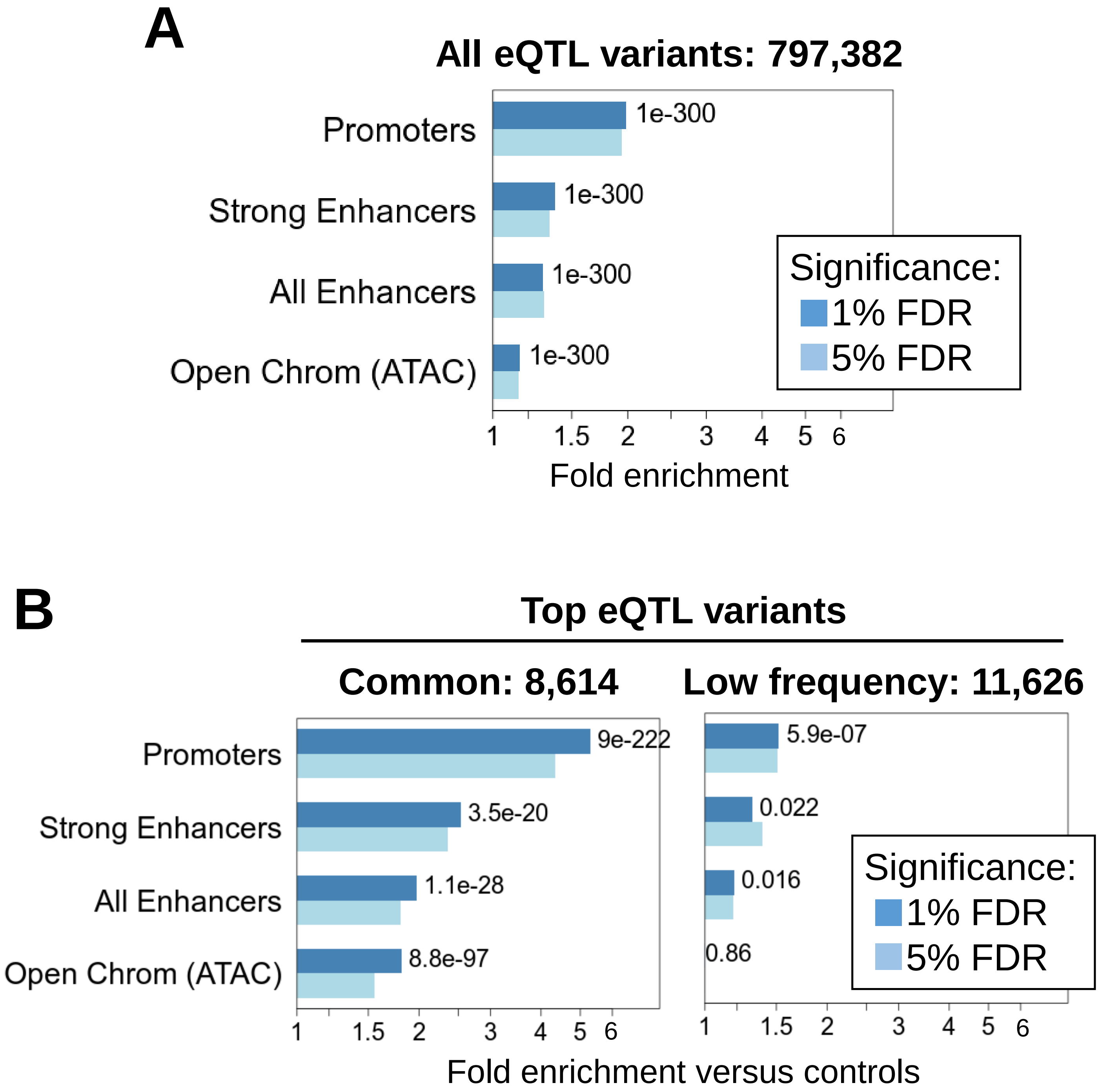

### Supplemental Figure 3

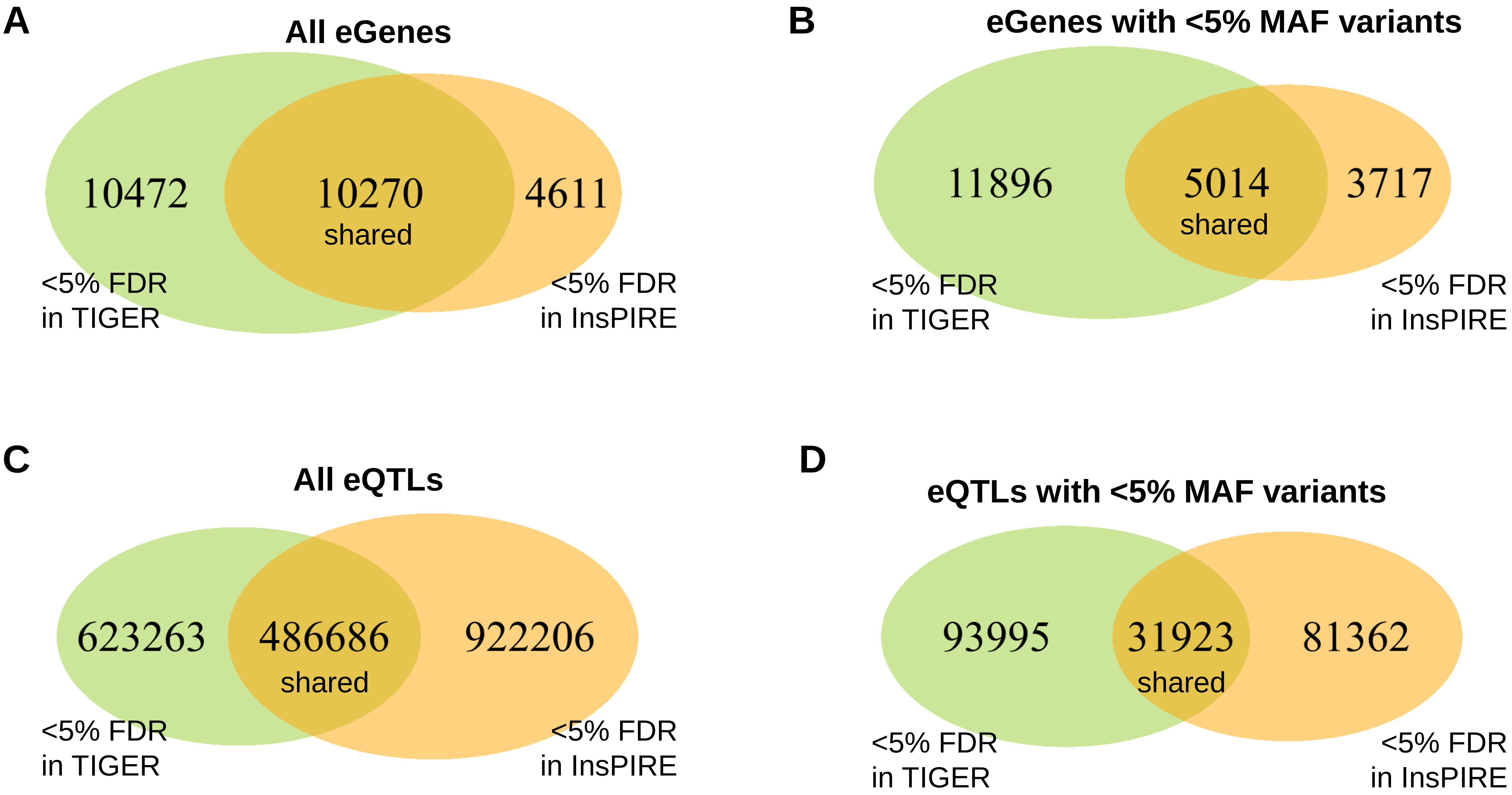

### Supplemental Figure 4

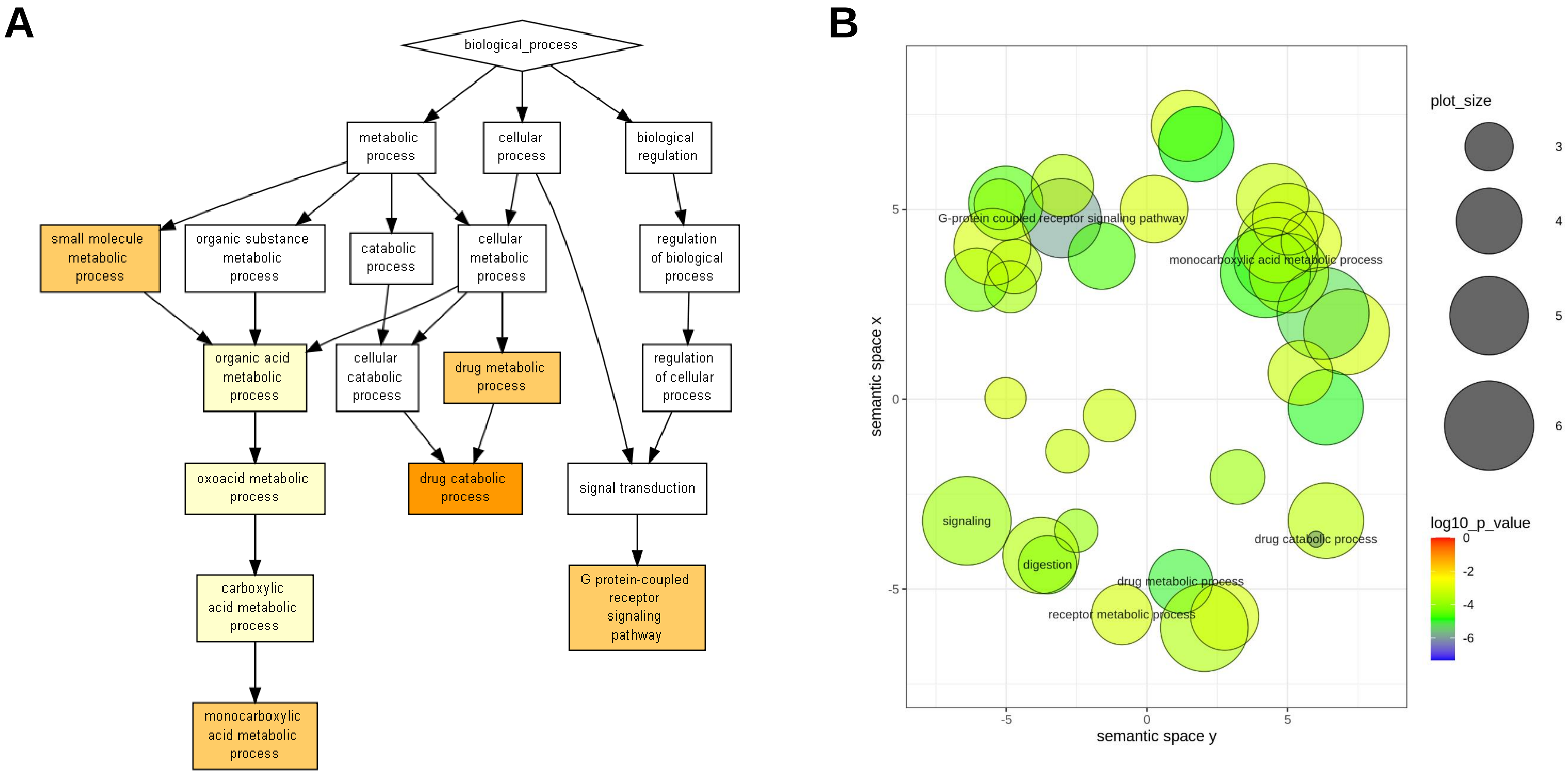

### Supplemental Figure 6

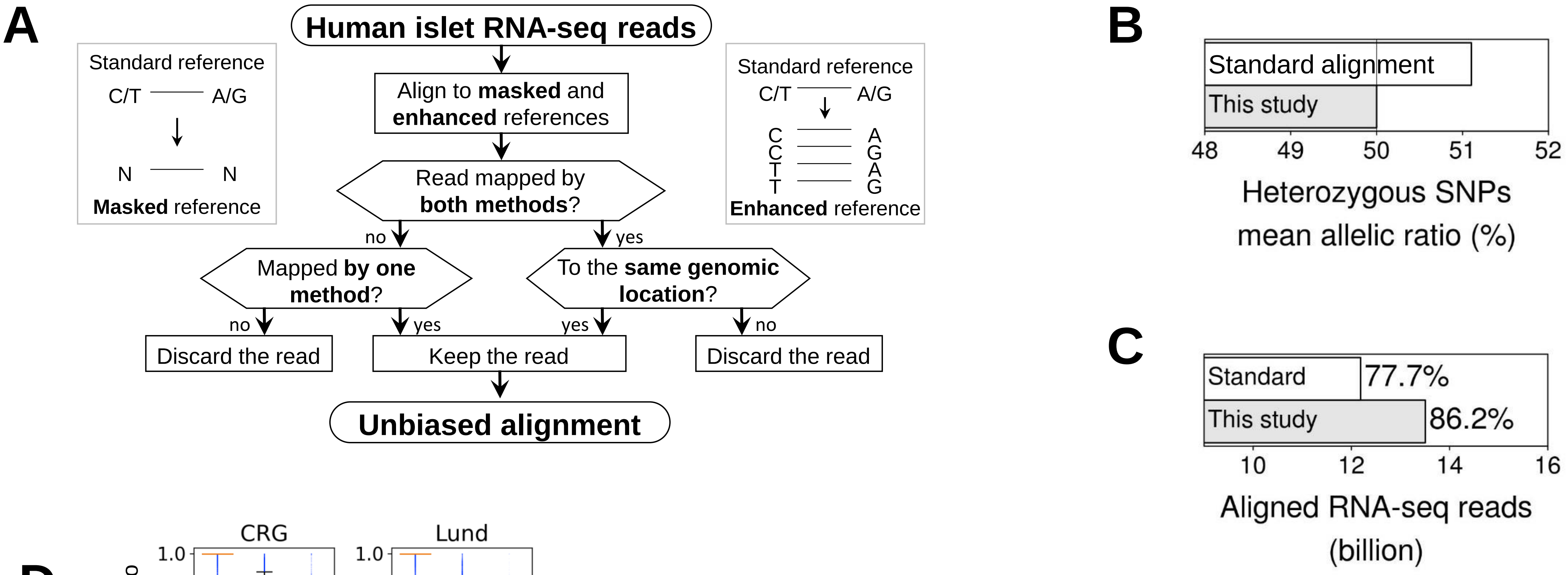


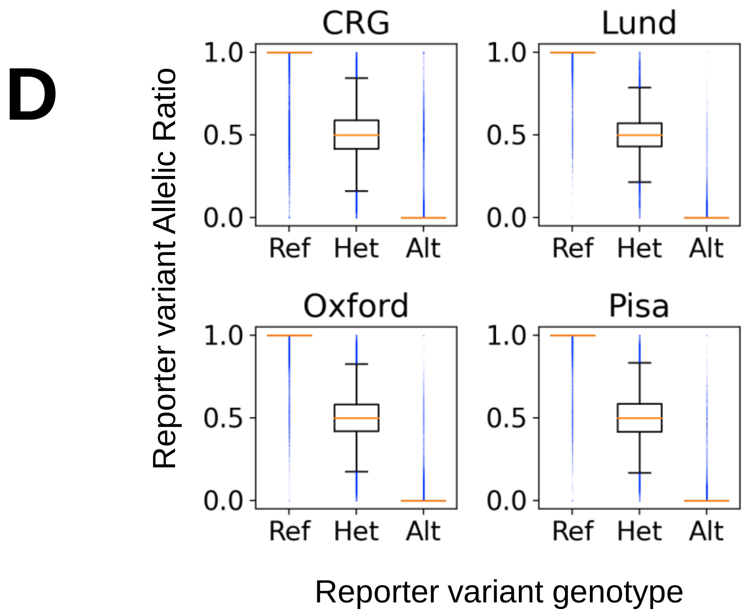


**E**

**Het reporters (Allelic Ratio)**

| Cohort | Mean | Median |
|---|---|---|
| CRG | 0.504 | 0.500 |
| Lund | 0.501 | 0.500 |
| Oxford | 0.500 | 0.500 |
| Pisa | 0.501 | 0.500 |

### Supplemental Figure 7

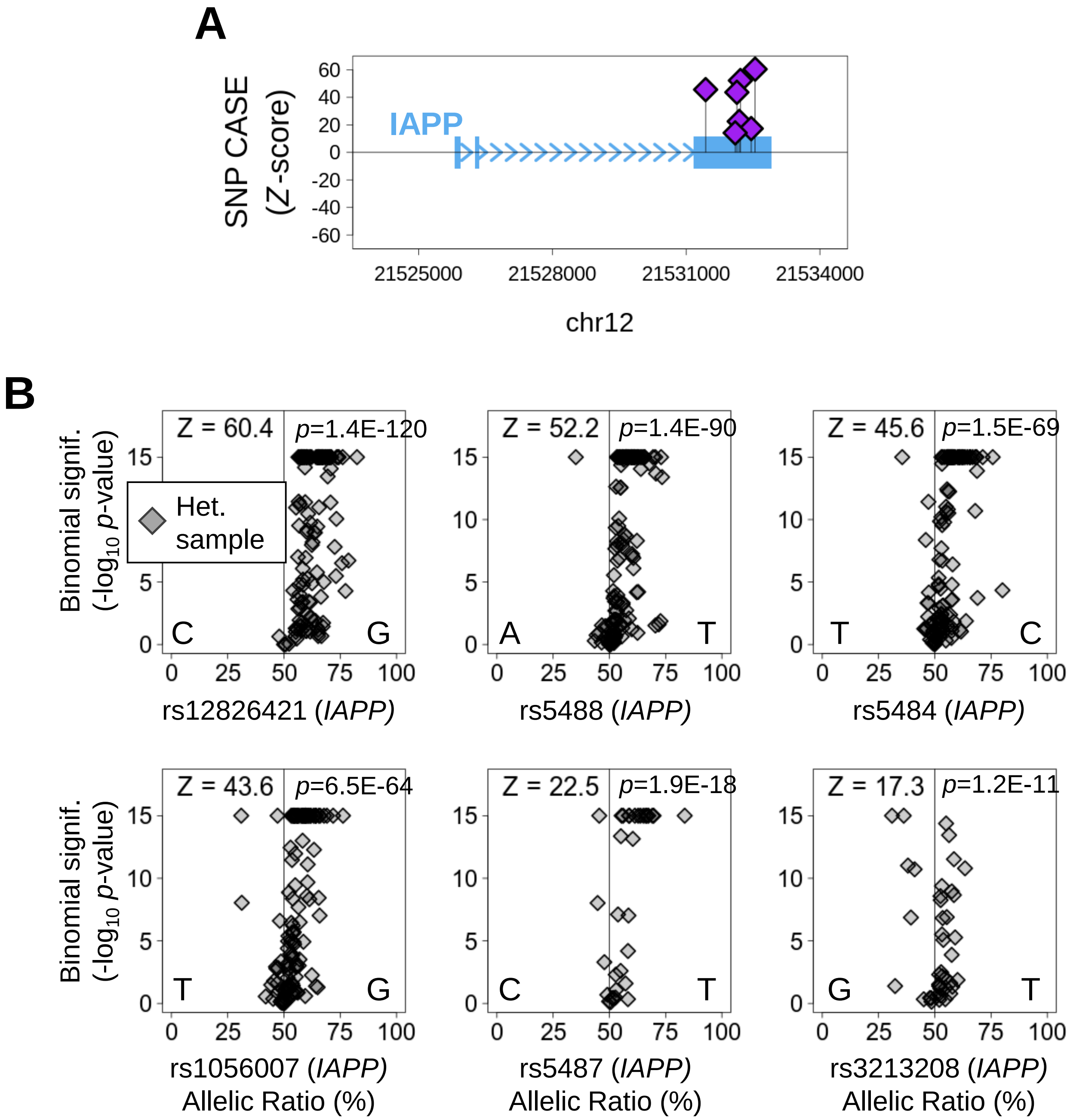

### Supplemental Figure 11

#### Number of good quality imputed genomic variants by chromosome

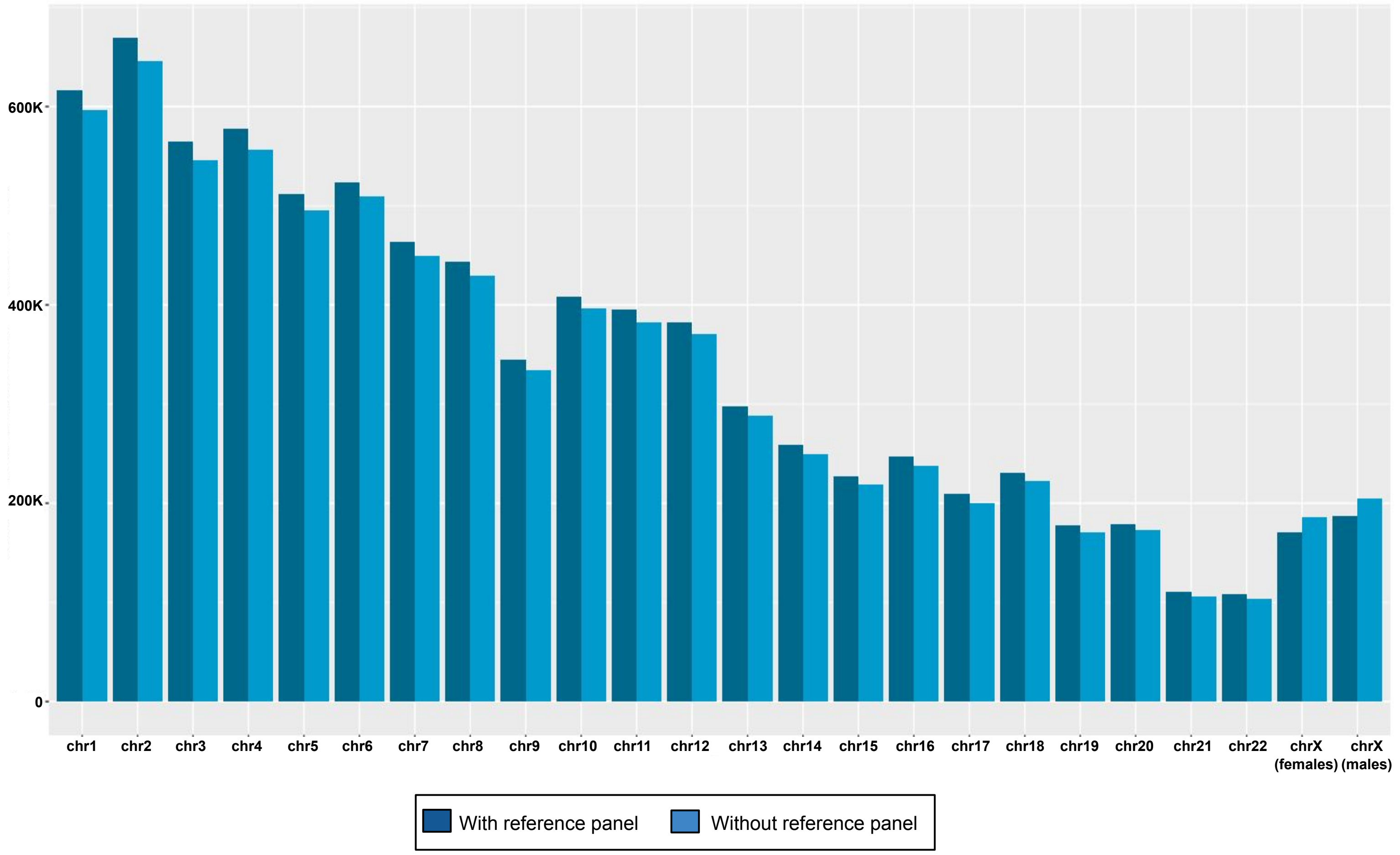

### Supplemental Figure 12

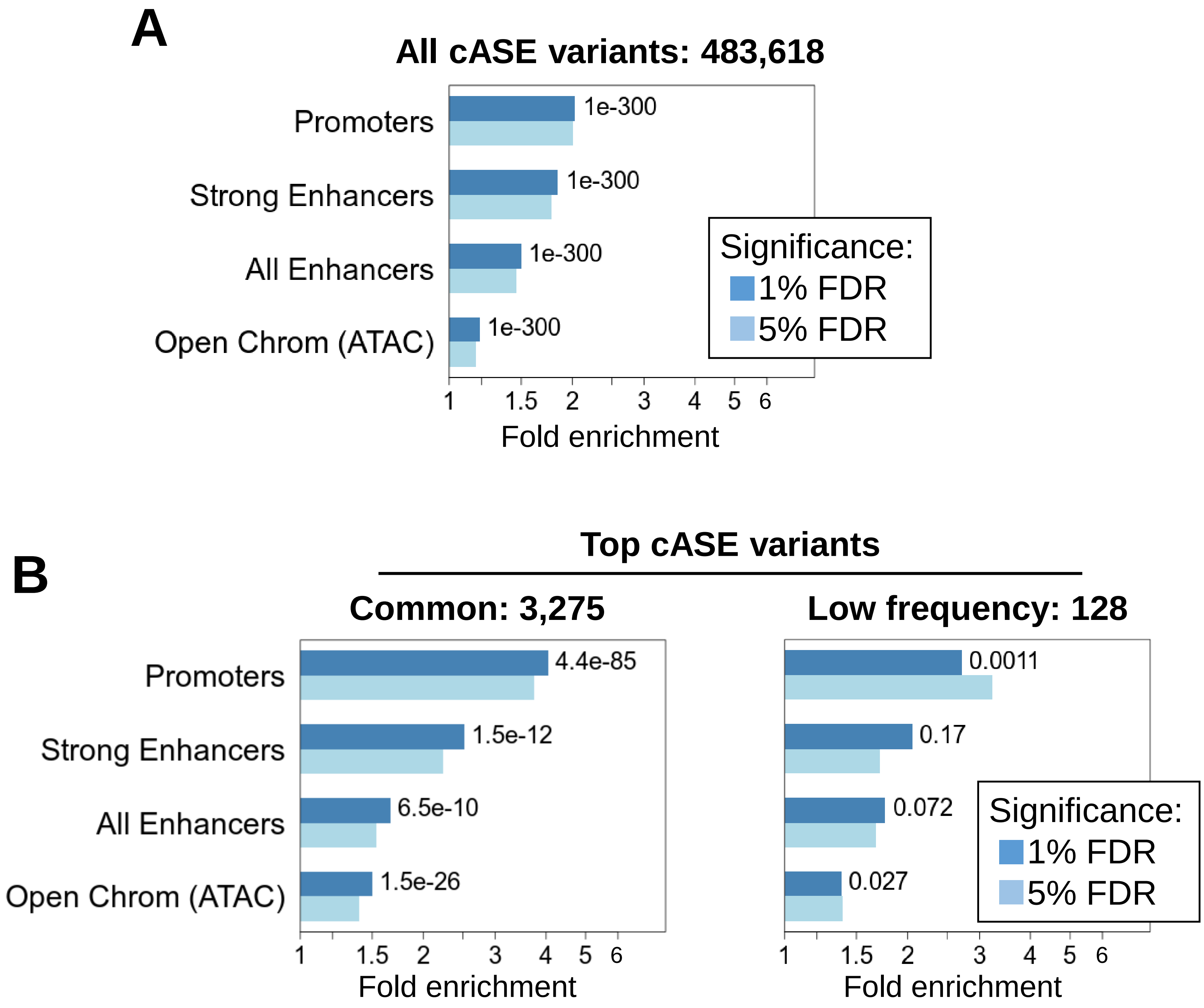
