## Supplemental Figure 5 for "TIGER: The gene expression regulatory variation landscape of human pancreatic islets"

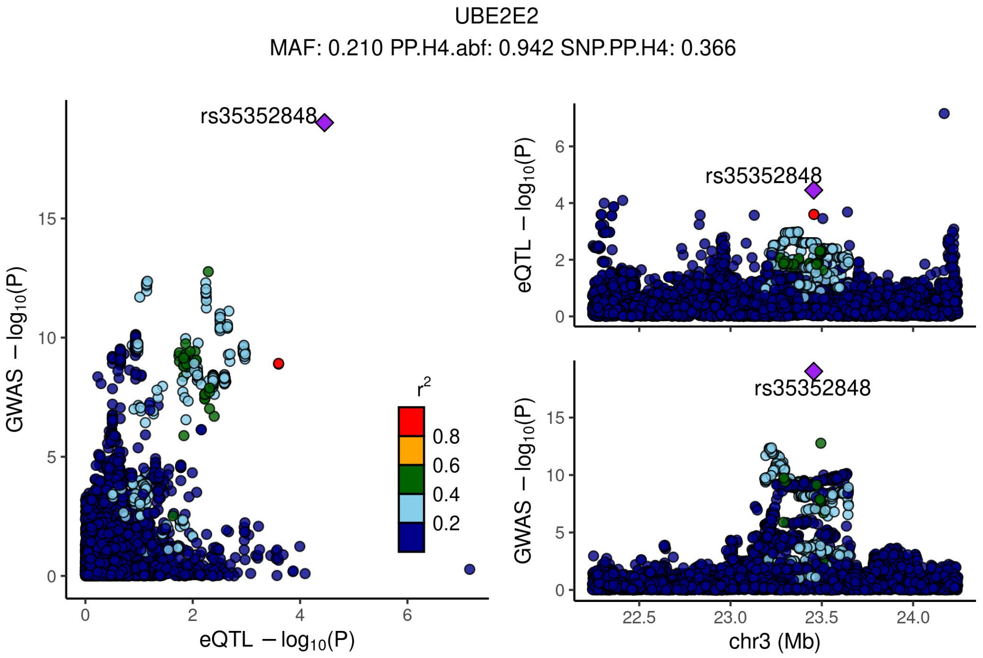


Supplementary Figure 4.1 - Locus compare plots 1Mbp up or down the TSS of UBE2E2.

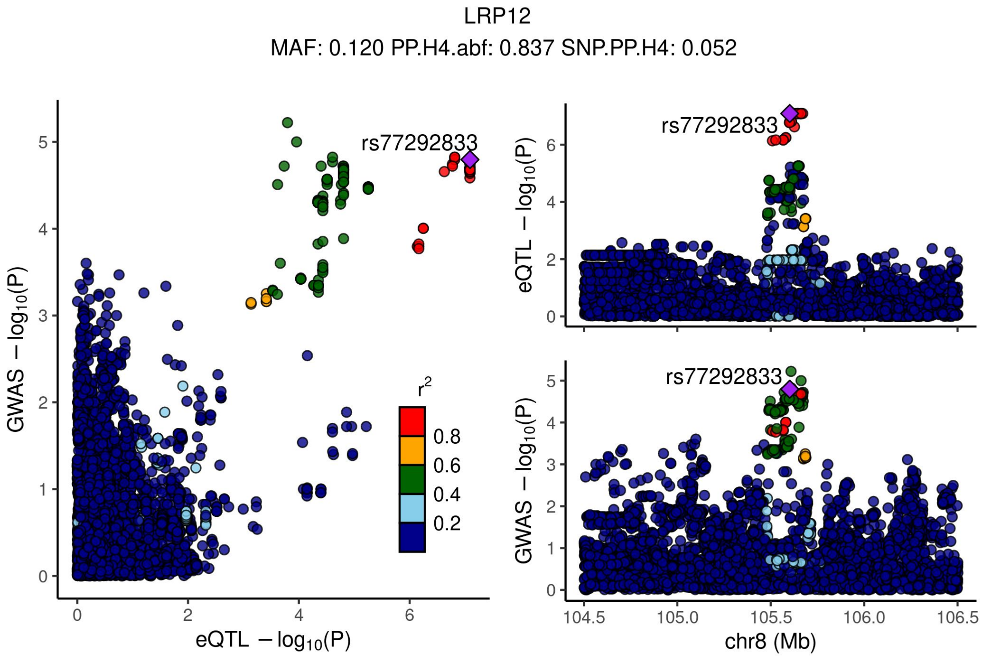


Supplementary Figure 4.2 - Locus compare plots 1Mbp up or down the TSS of LRP12.

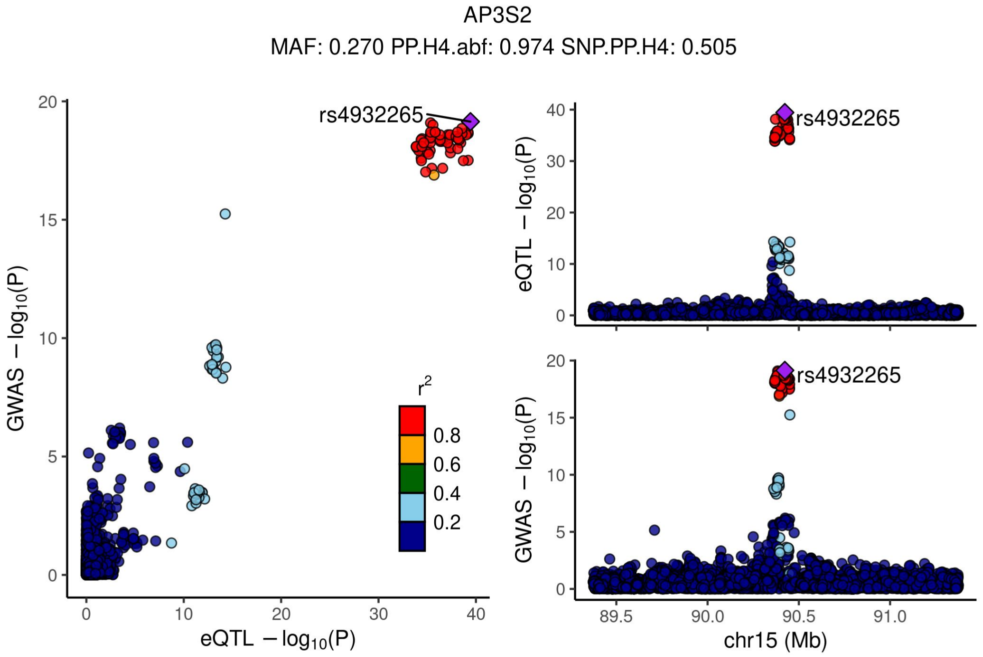


Supplementary Figure 4.3 - Locus compare plots 1Mbp up or down the TSS of AP3S2.

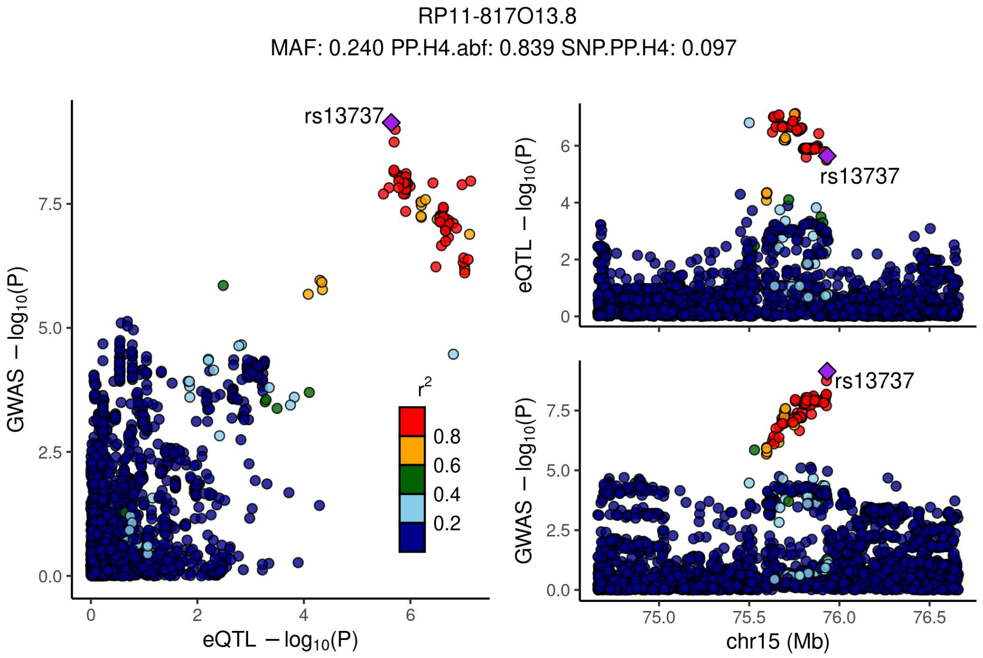


Supplementary Figure 4.4 - Locus compare plots 1Mbp up or down the TSS of RP11-817O13.8.

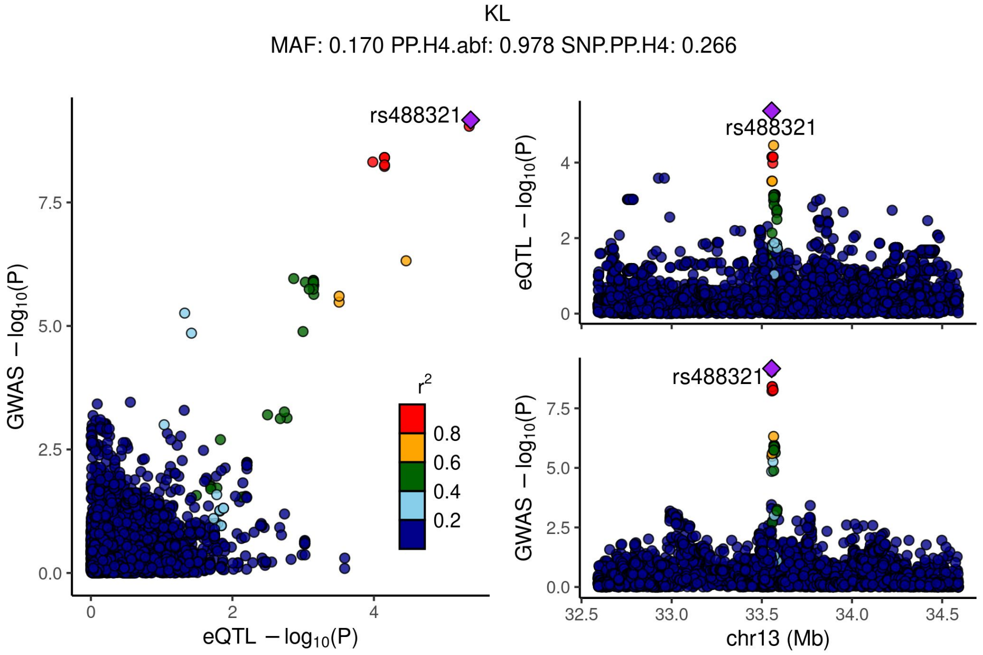


Supplementary Figure 4.5 - Locus compare plots 1Mbp up or down the TSS of KL.

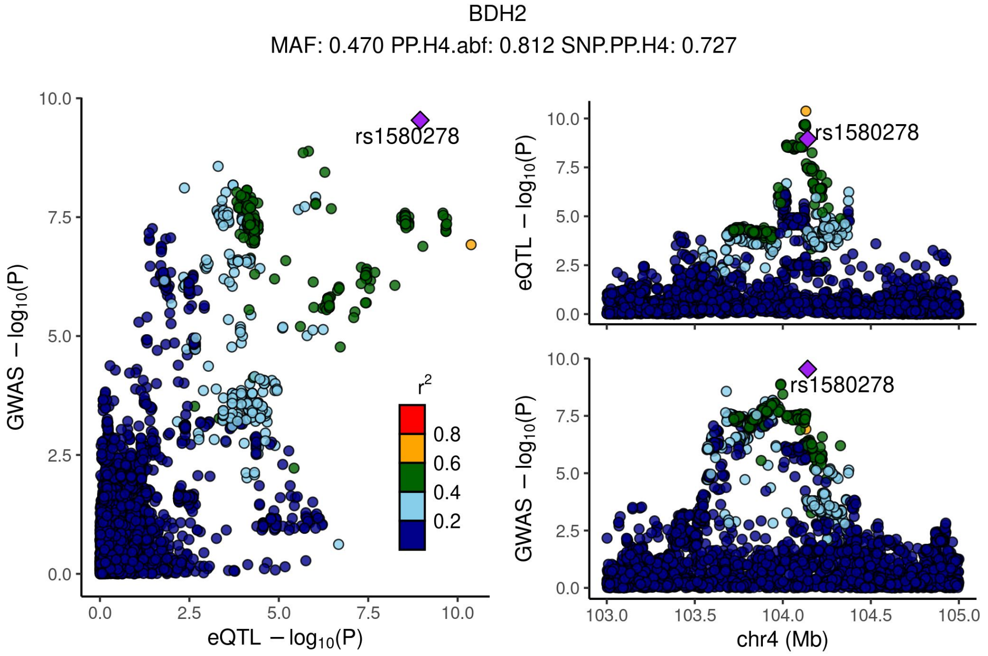


Supplementary Figure 4.6 - Locus compare plots 1Mbp up or down the TSS of BDH2.

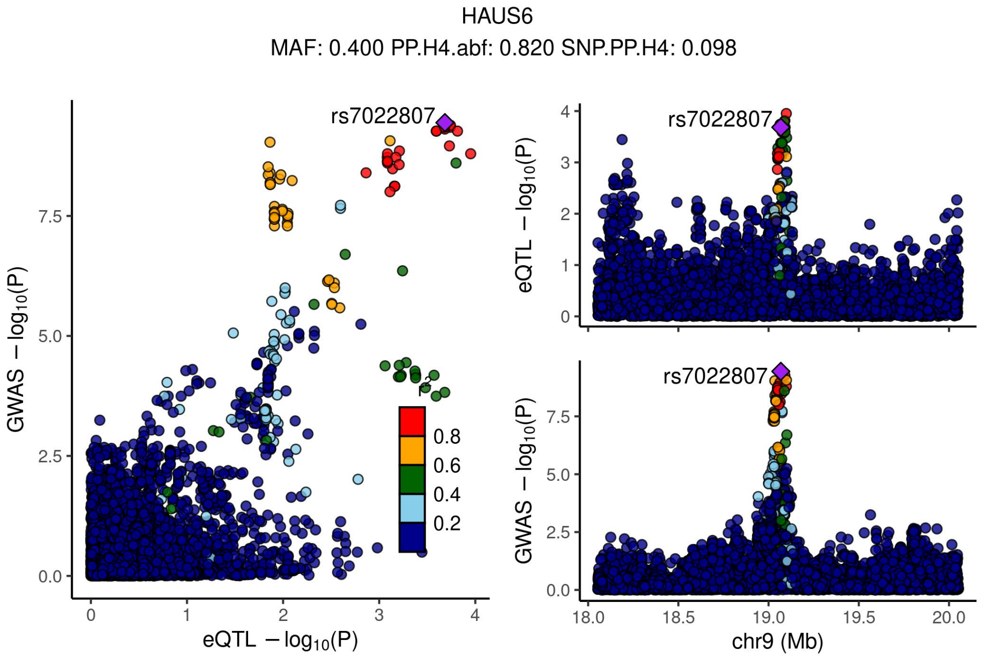


Supplementary Figure 4.7 - Locus compare plots 1Mbp up or down the TSS of HAUS6.

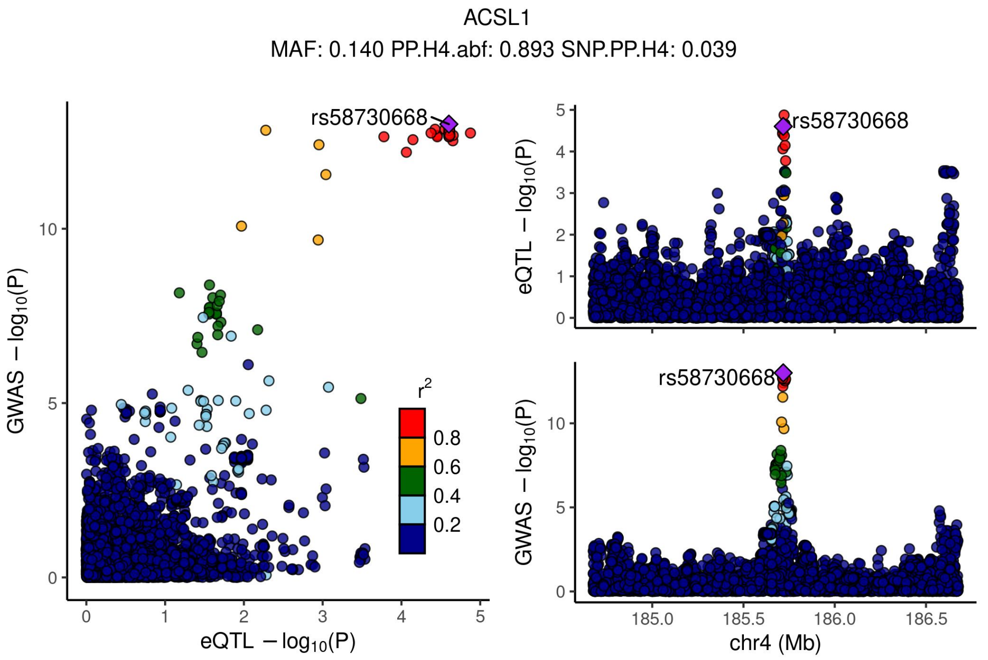


Supplementary Figure 4.8 - Locus compare plots 1Mbp up or down the TSS of ACSL1.

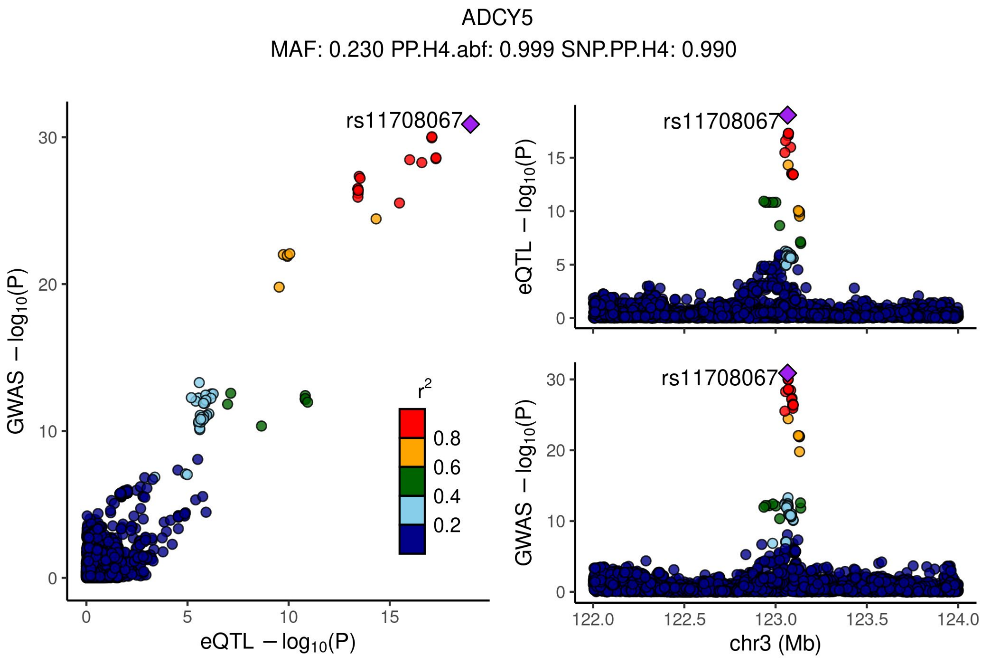


Supplementary Figure 4.9 - Locus compare plots 1Mbp up or down the TSS of ADCY5.

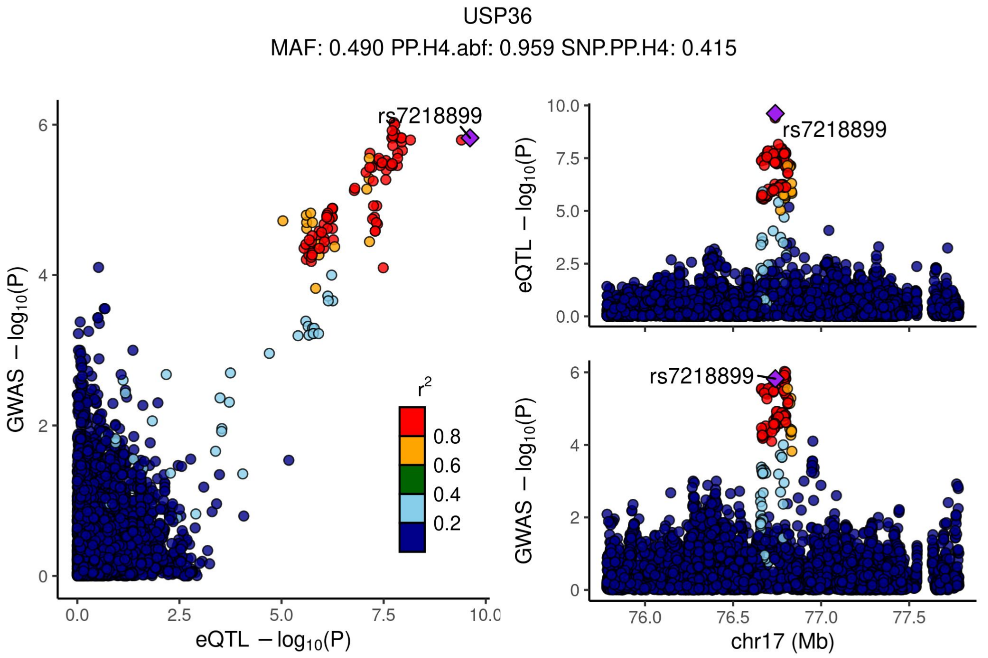


Supplementary Figure 4.10 - Locus compare plots 1Mbp up or down the TSS of USP36.

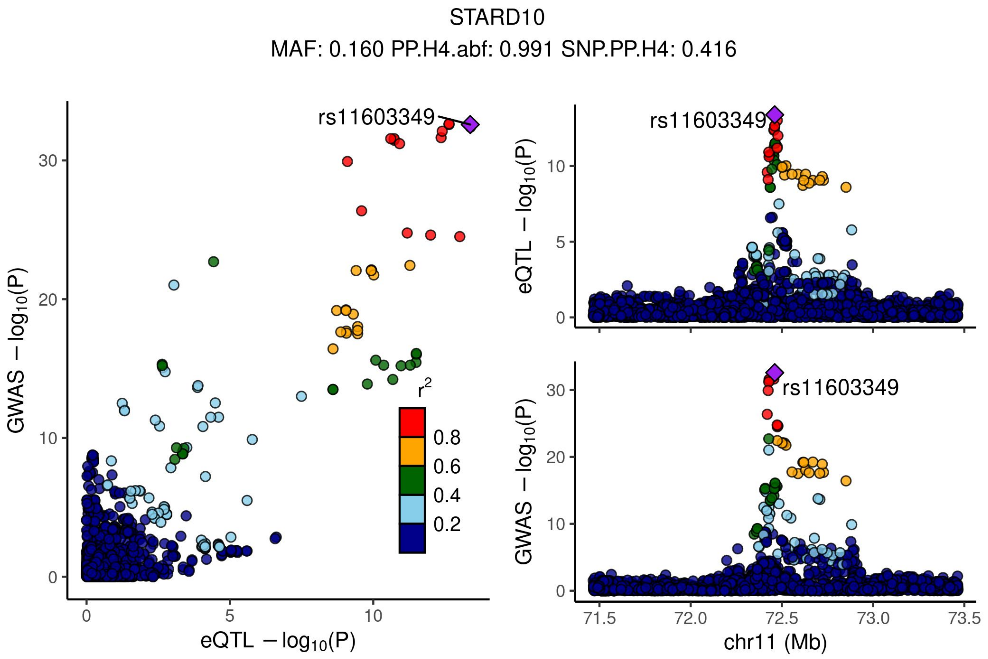


Supplementary Figure 4.11 - Locus compare plots 1Mbp up or down the TSS of STARD10.

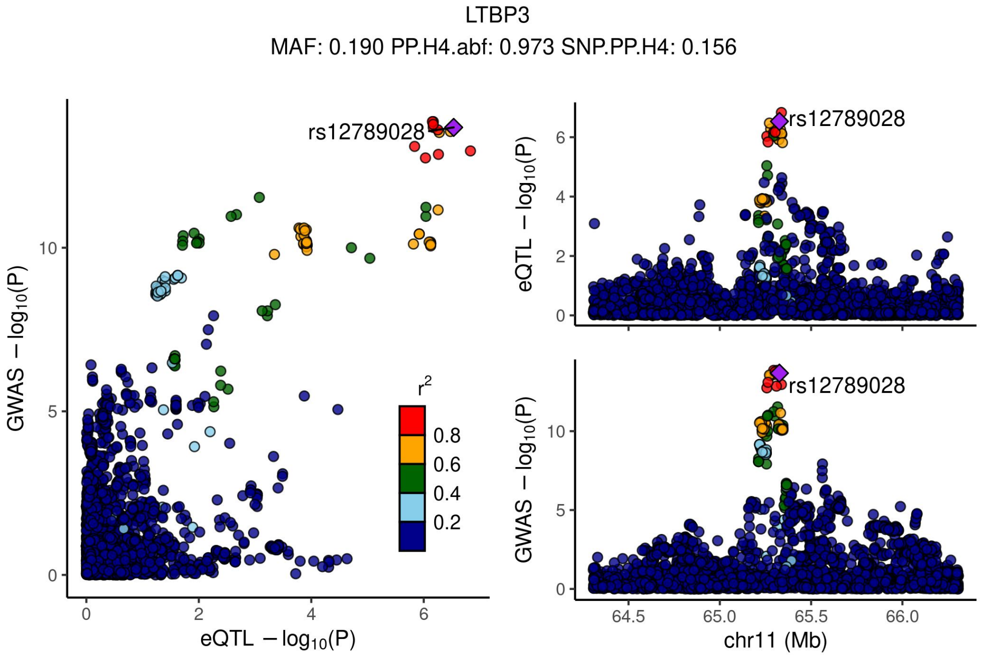


Supplementary Figure 4.12 - Locus compare plots 1Mbp up or down the TSS of LTBP3.

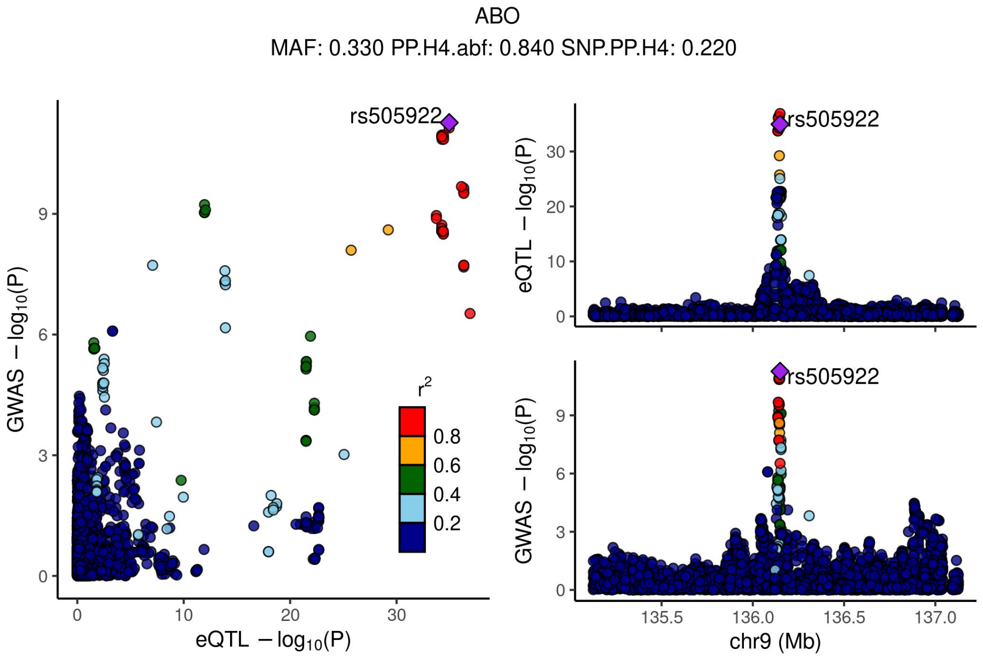


Supplementary Figure 4.13 - Locus compare plots 1Mbp up or down the TSS of ABO.

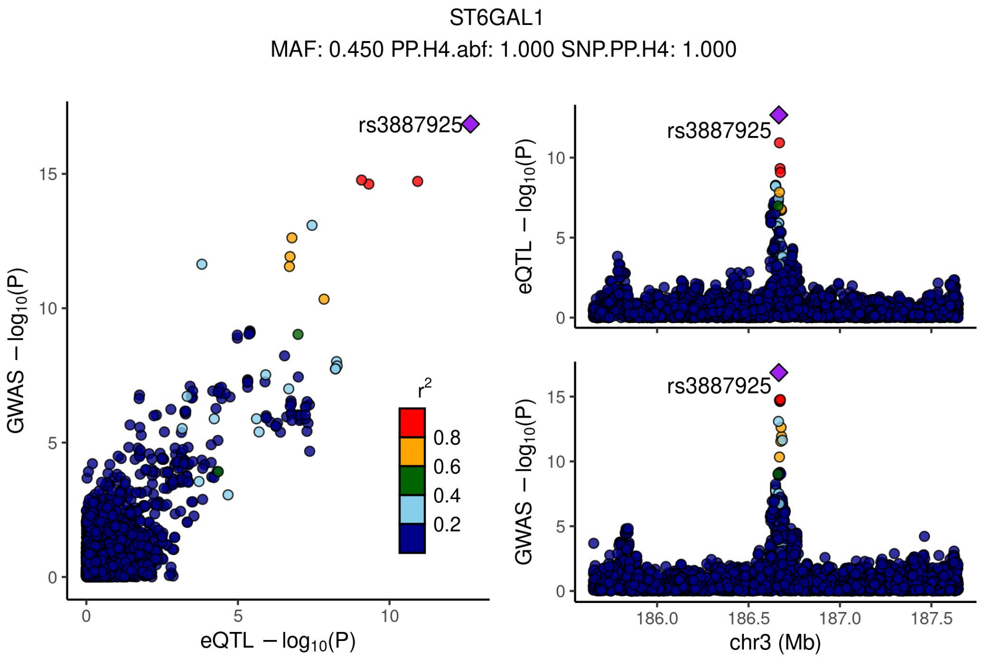


Supplementary Figure 4.14 - Locus compare plots 1Mbp up or down the TSS of ST6GAL1.

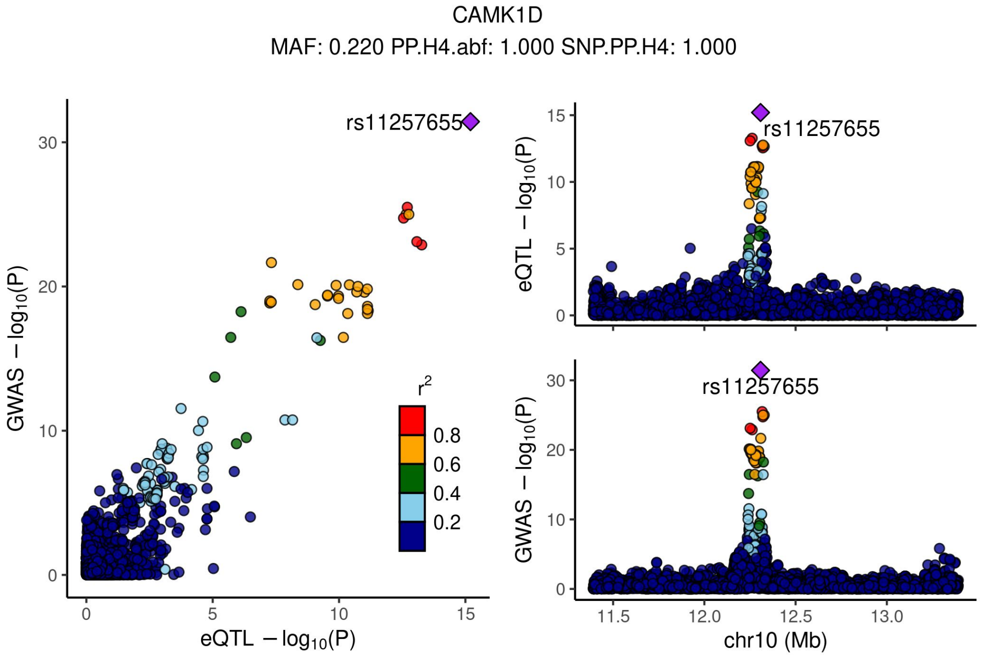


Supplementary Figure 4.15 - Locus compare plots 1Mbp up or down the TSS of CAMK1D.

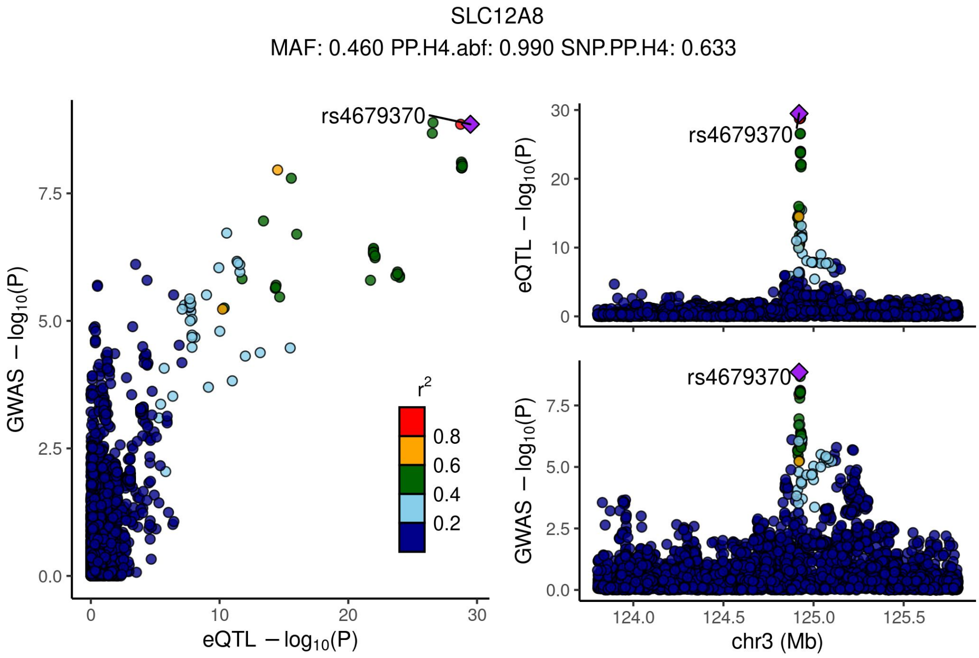


Supplementary Figure 4.16 - Locus compare plots 1Mbp up or down the TSS of SLC12A8.

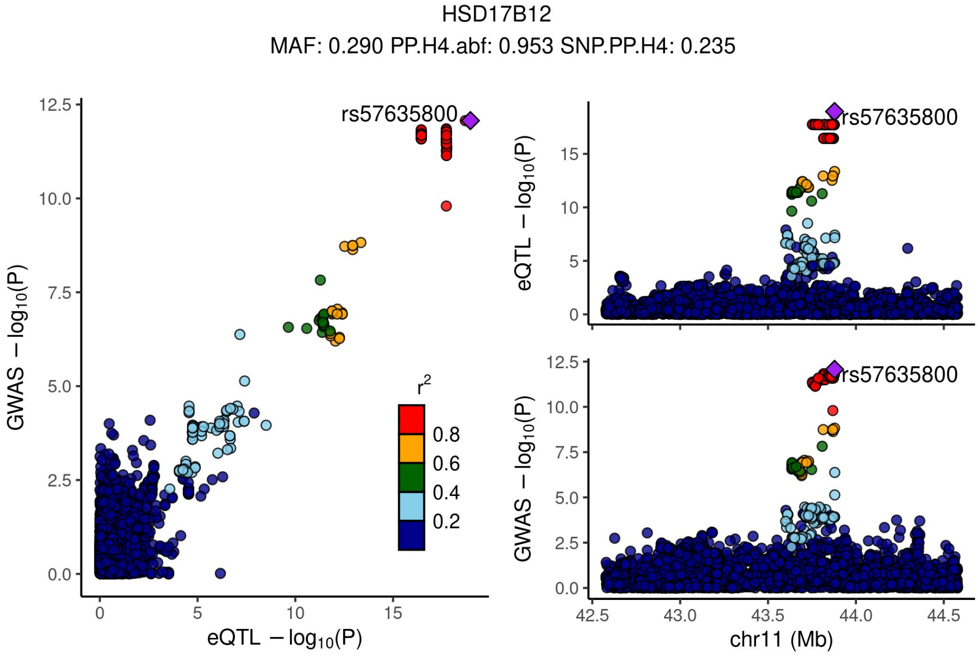


Supplementary Figure 4.17 - Locus compare plots 1Mbp up or down the TSS of HSD17B12.

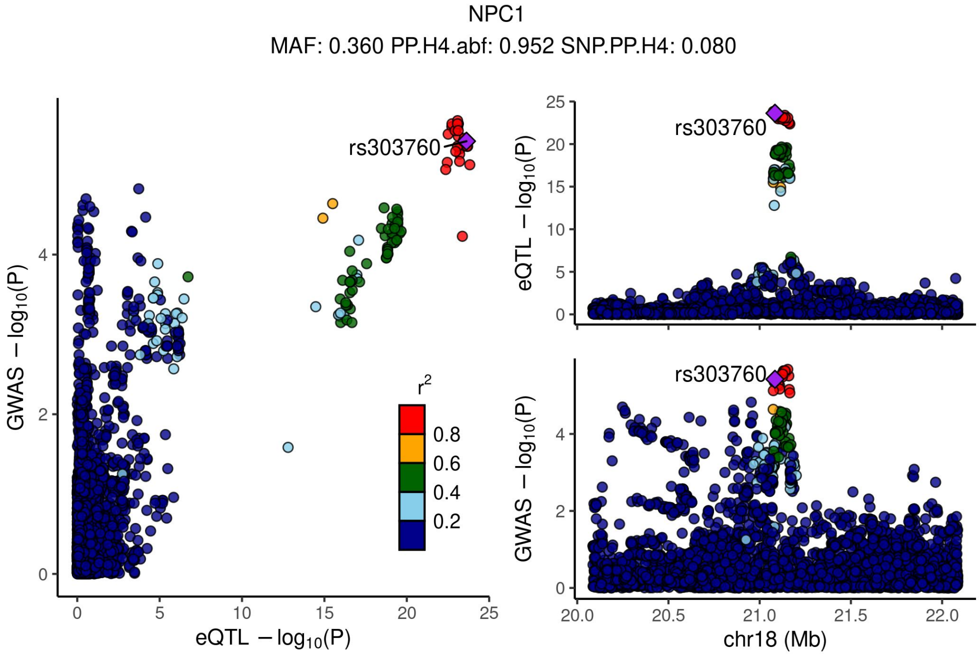


Supplementary Figure 4.18 - Locus compare plots 1Mbp up or down the TSS of NPC1.

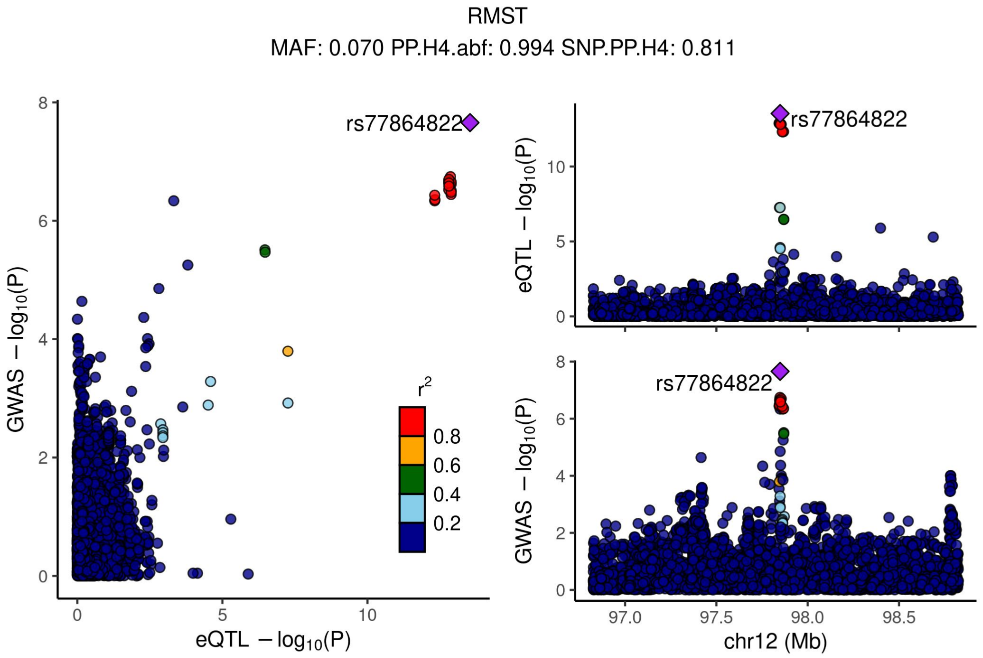


Supplementary Figure 4.19 - Locus compare plots 1Mbp up or down the TSS of RMST.

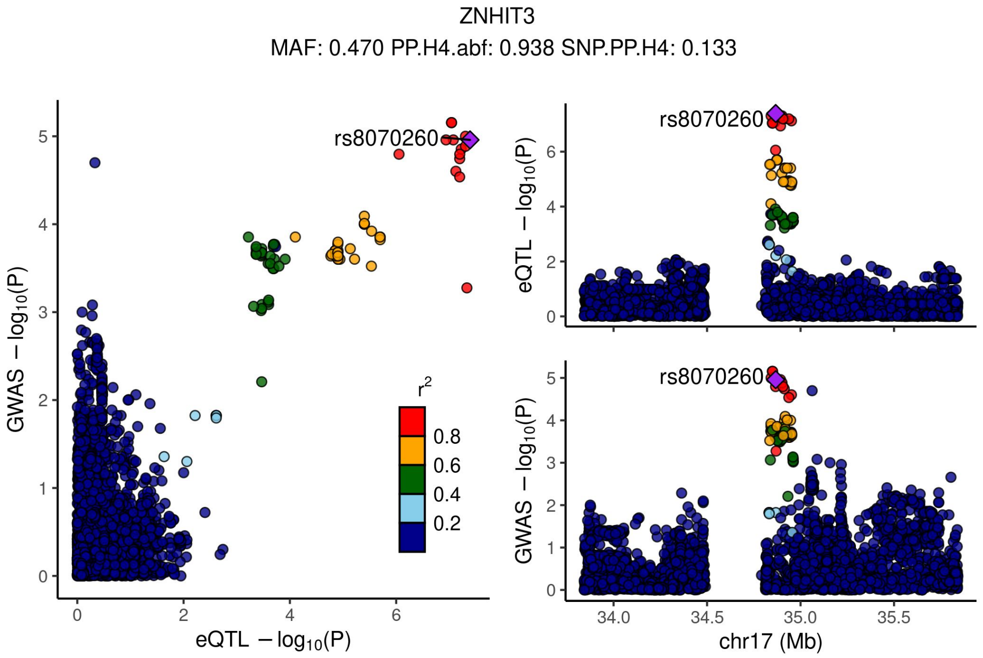


Supplementary Figure 4.20 - Locus compare plots 1Mbp up or down the TSS of ZNHIT3.

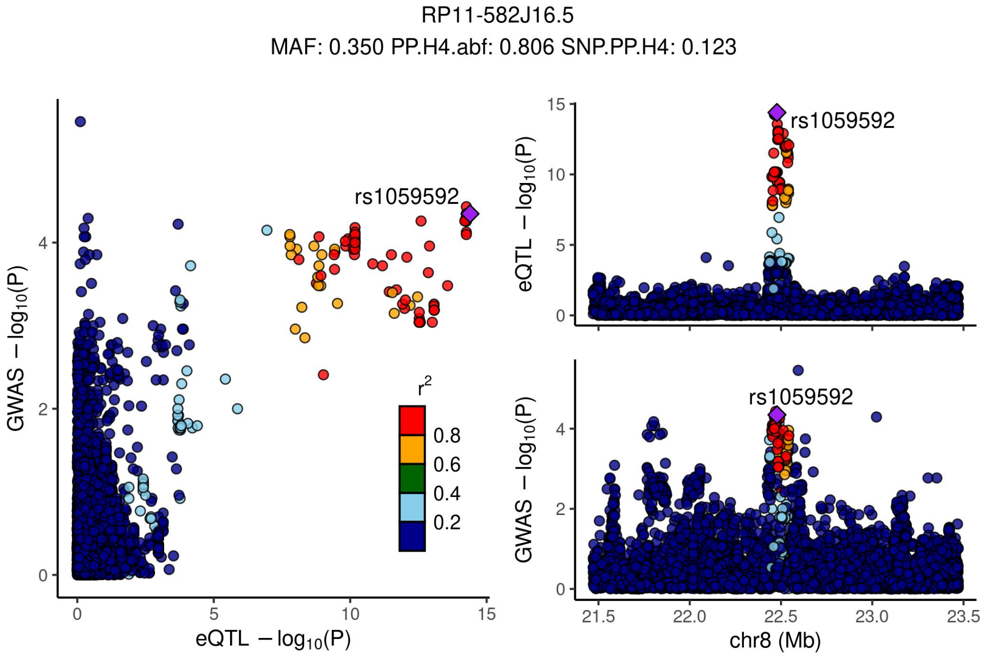


Supplementary Figure 4.21 - Locus compare plots 1Mbp up or down the TSS of RP11-582J16.5.

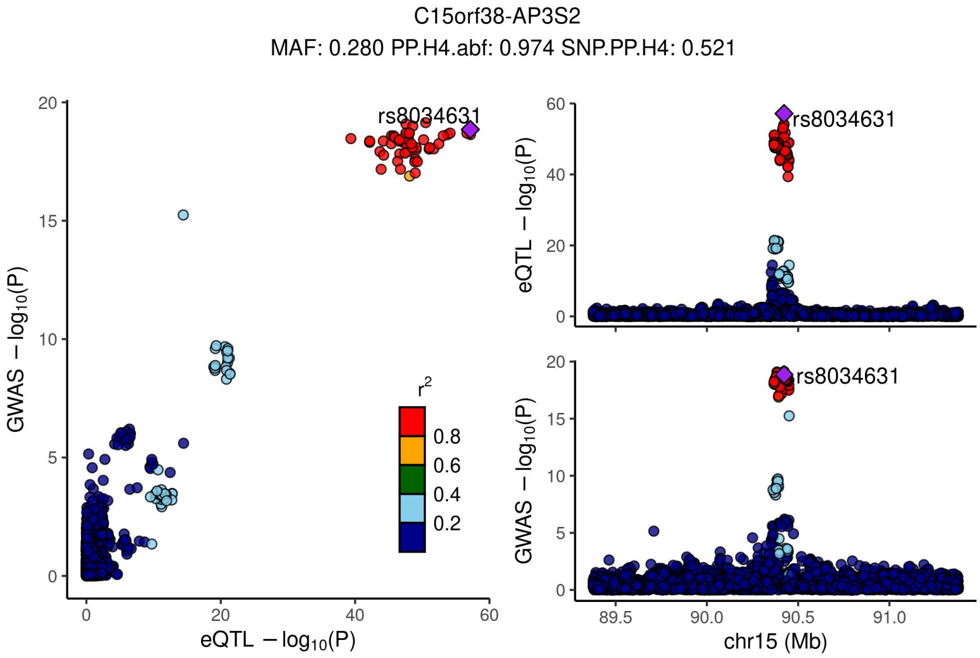


Supplementary Figure 4.22 - Locus compare plots 1Mbp up or down the TSS of C15orf38-AP3S2.

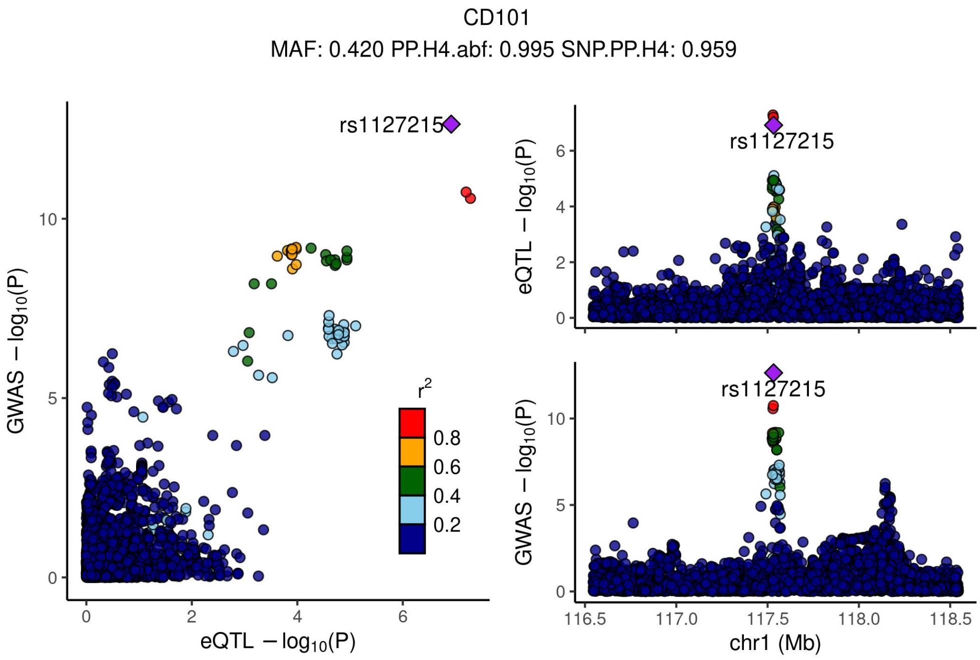


Supplementary Figure 4.23 - Locus compare plots 1Mbp up or down the TSS of CD101.

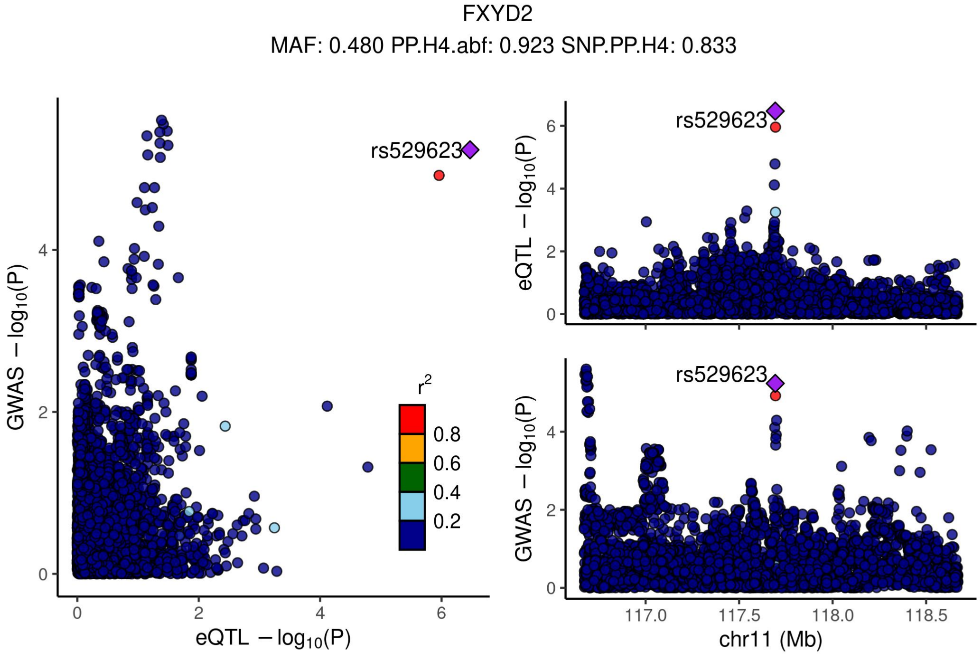


Supplementary Figure 4.24 - Locus compare plots 1Mbp up or down the TSS of FXYD2.

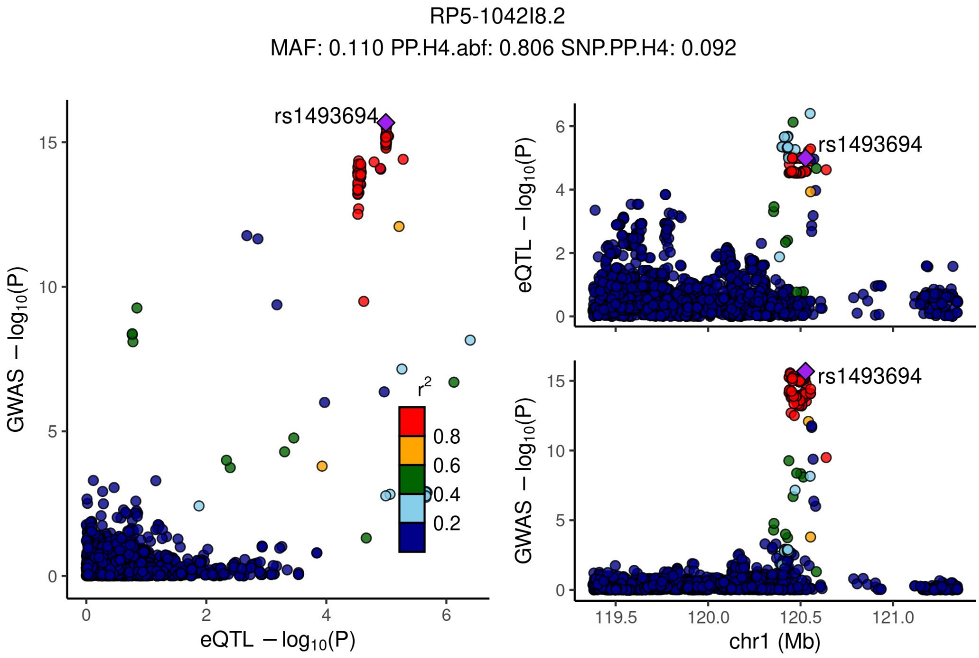


Supplementary Figure 4.25 - Locus compare plots 1Mbp up or down the TSS of RP5-1042I8.2.

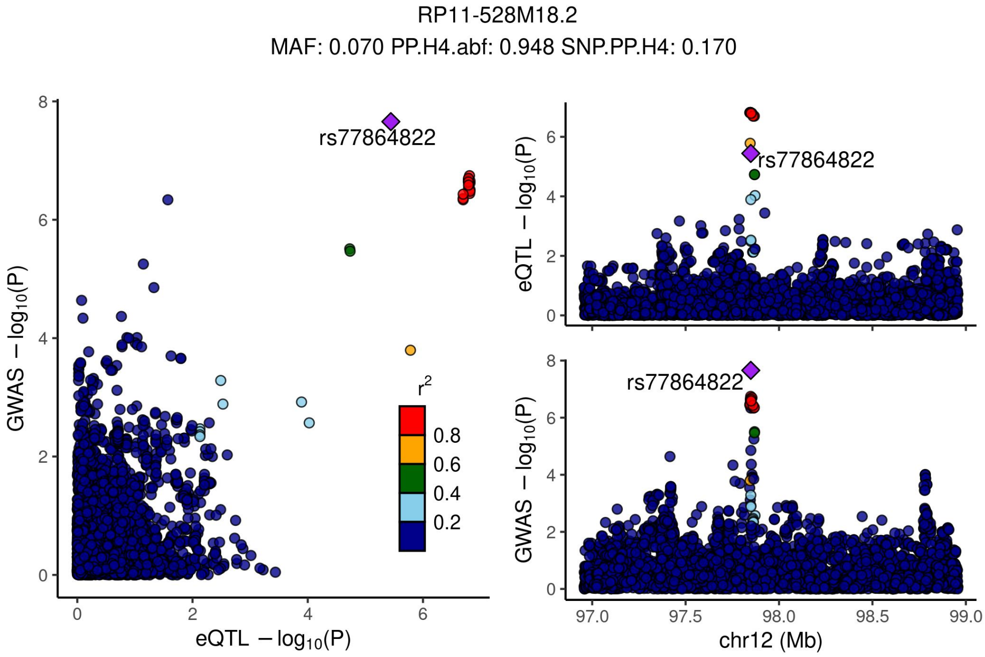


Supplementary Figure 4.26 - Locus compare plots 1Mbp up or down the TSS of RP11-528M18.2.

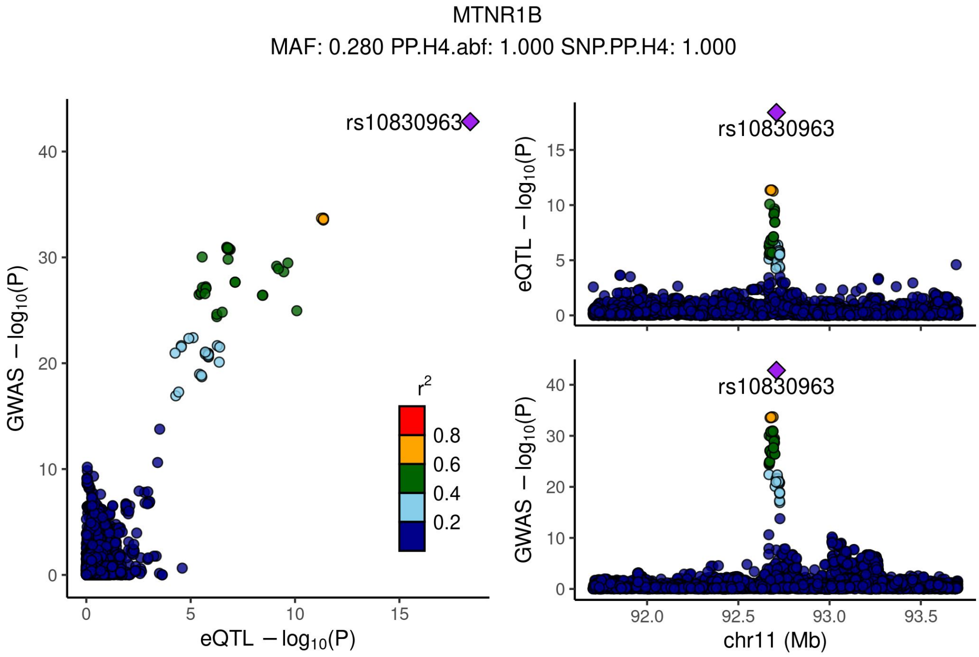


Supplementary Figure 4.27 - Locus compare plots 1Mbp up or down the TSS of MTNR1B.

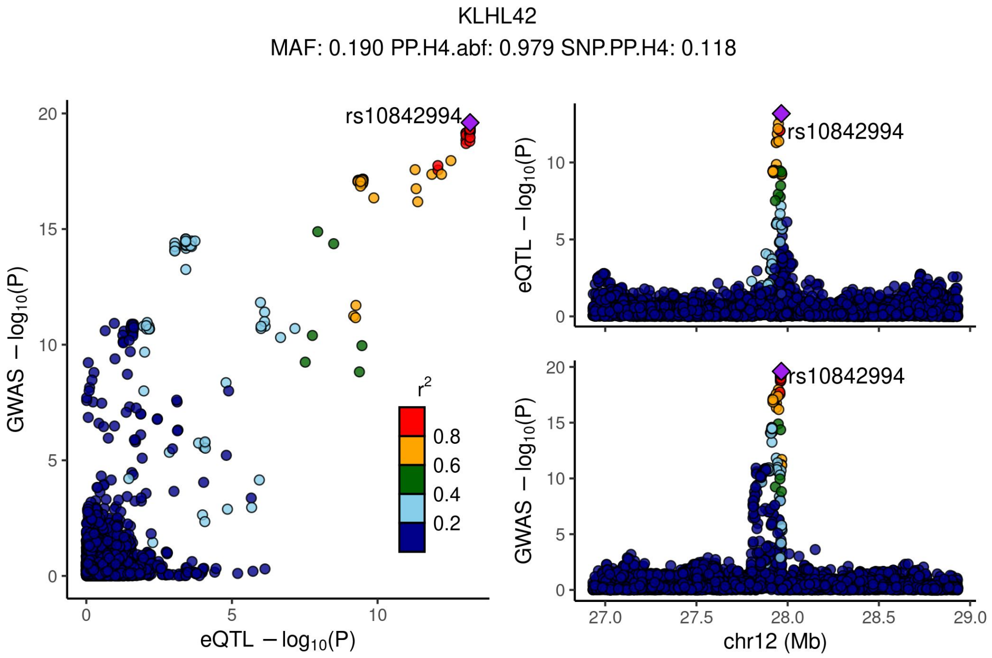


Supplementary Figure 4.28 - Locus compare plots 1Mbp up or down the TSS of KLHL42.

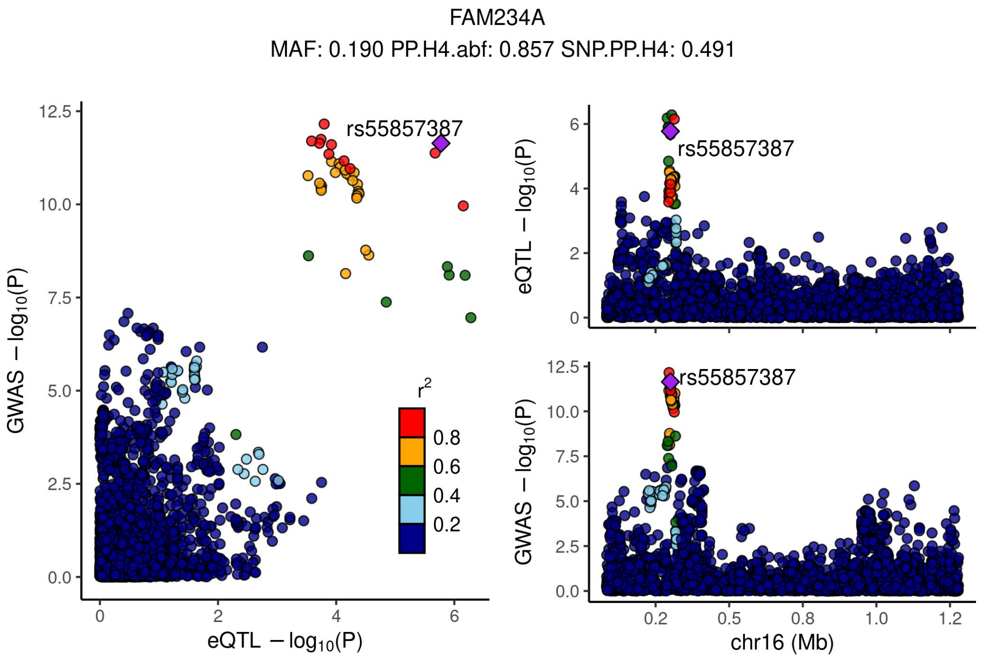


Supplementary Figure 4.29 - Locus compare plots 1Mbp up or down the TSS of FAM234A.

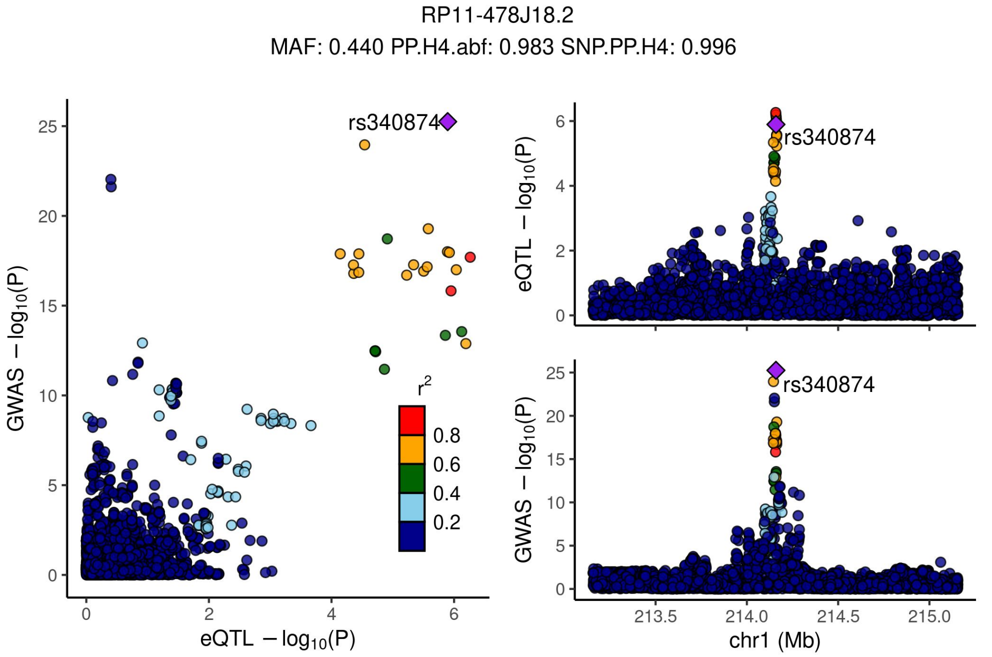


Supplementary Figure 4.30 - Locus compare plots 1Mbp up or down the TSS of RP11-478J18.2.

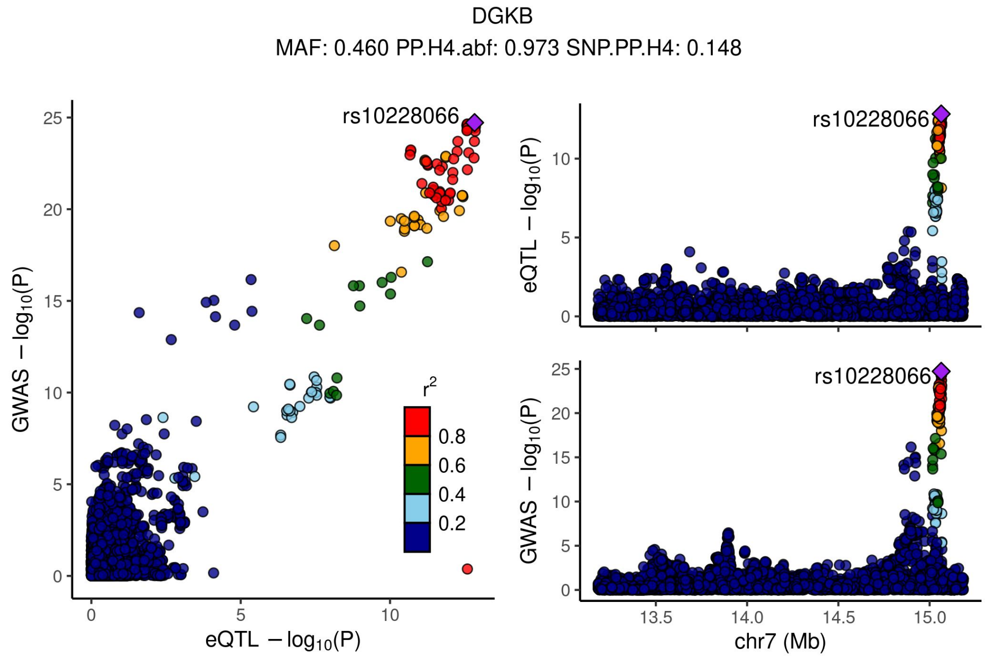


Supplementary Figure 4.31 - Locus compare plots 1Mbp up or down the TSS of DGKB.

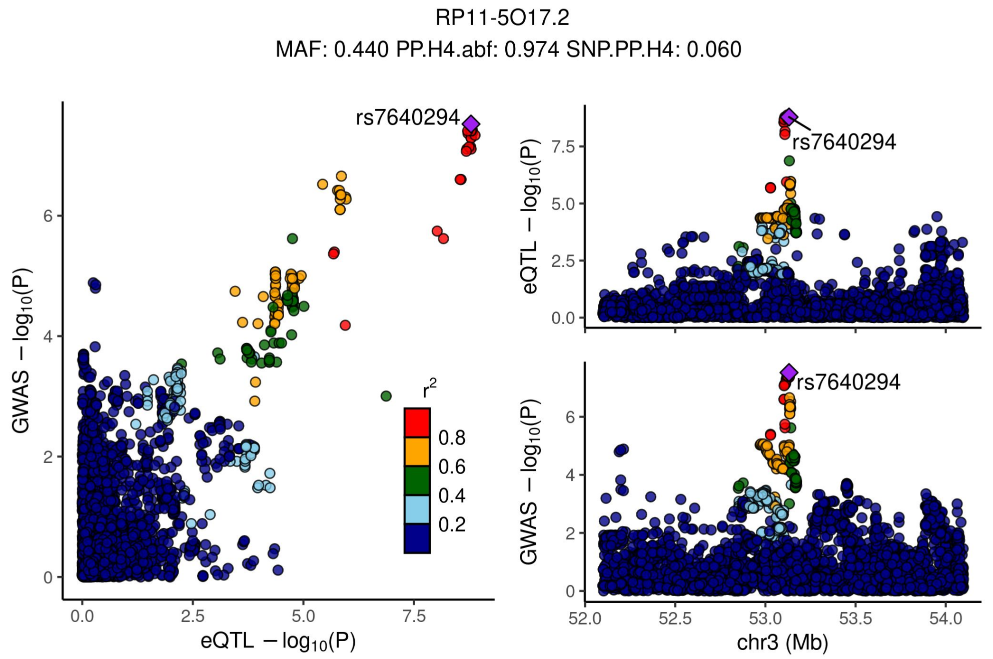


Supplementary Figure 4.32 - Locus compare plots 1Mbp up or down the TSS of RP11-5O17.2.

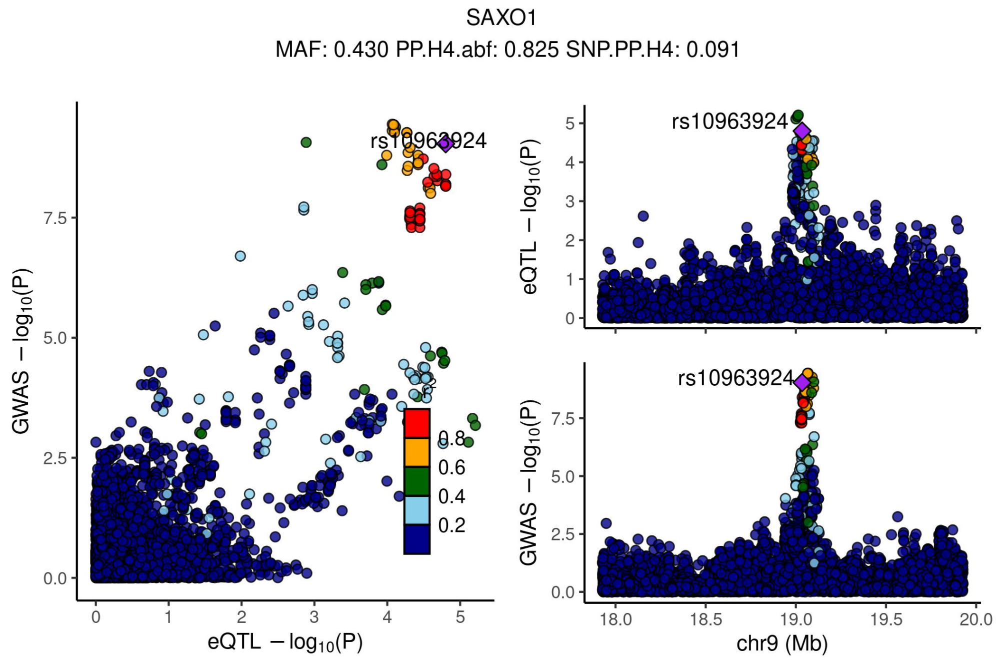


Supplementary Figure 4.33 - Locus compare plots 1Mbp up or down the TSS of SAXO1.

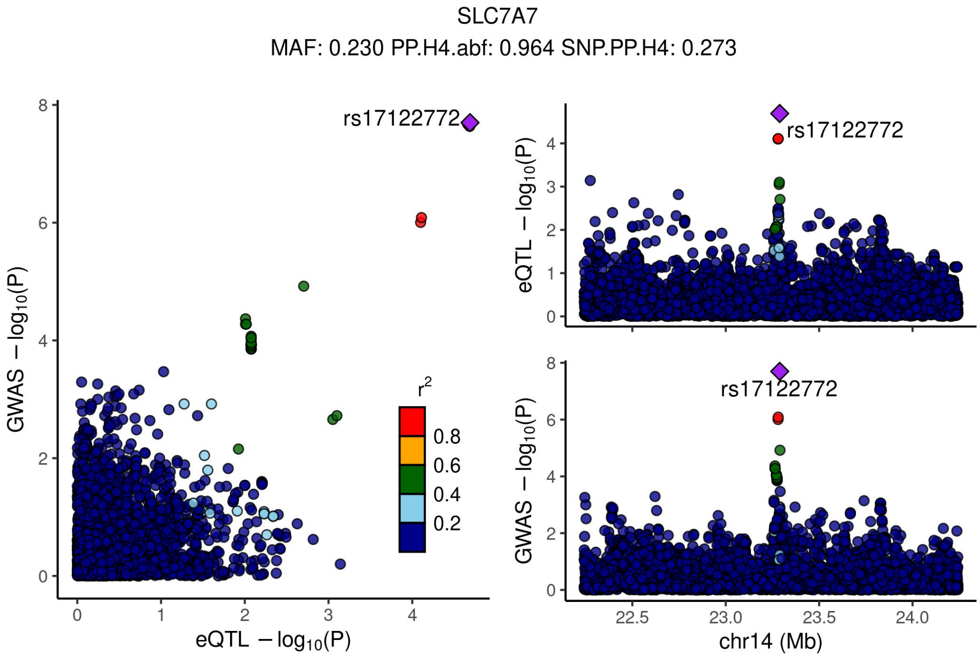


Supplementary Figure 4.34 - Locus compare plots 1Mbp up or down the TSS of SLC7A7.

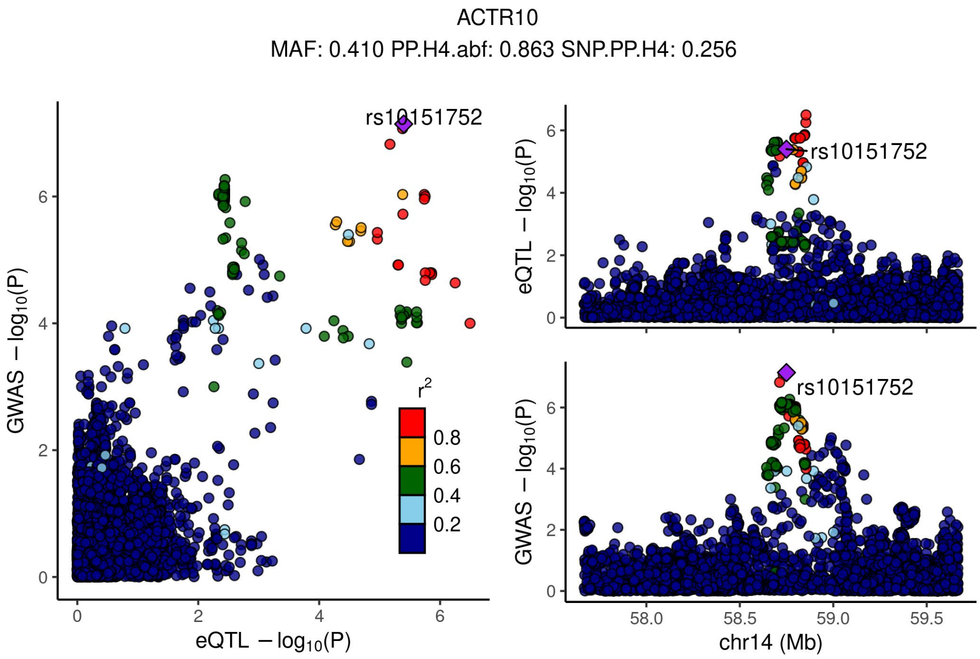


Supplementary Figure 4.35 - Locus compare plots 1Mbp up or down the TSS of ACTR10.

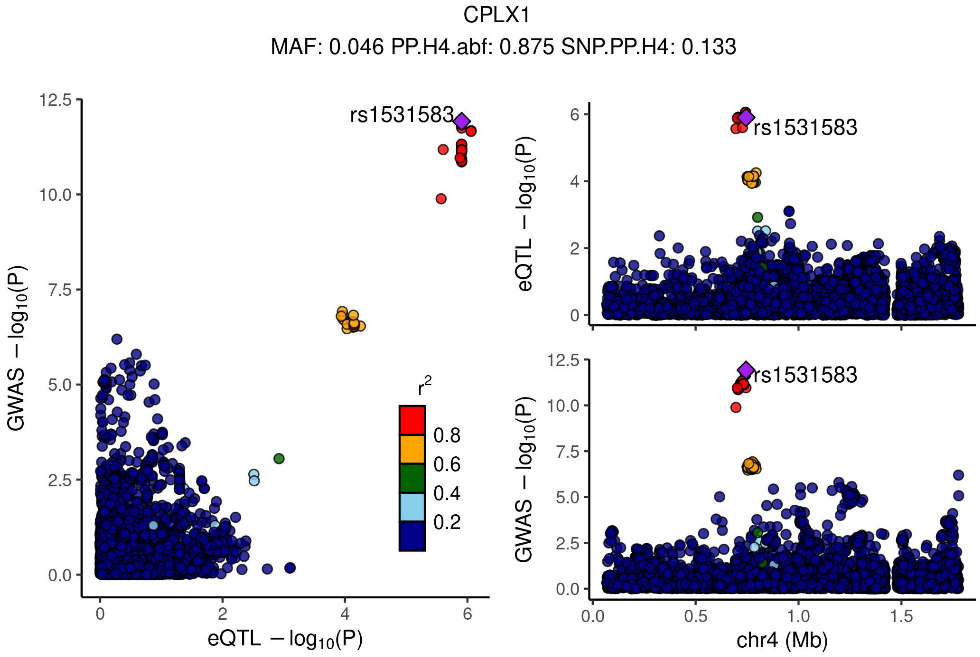


Supplementary Figure 4.36 - Locus compare plots 1Mbp up or down the TSS of CPLX1.

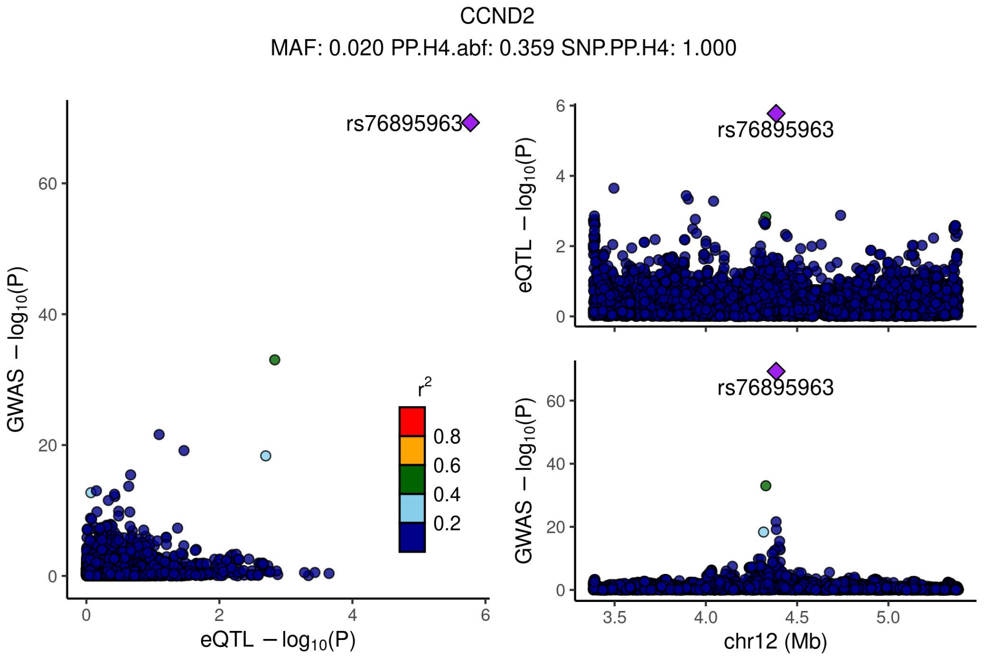


Supplementary Figure 4.37 - Locus compare plots 1Mbp up or down the TSS of CCND2.

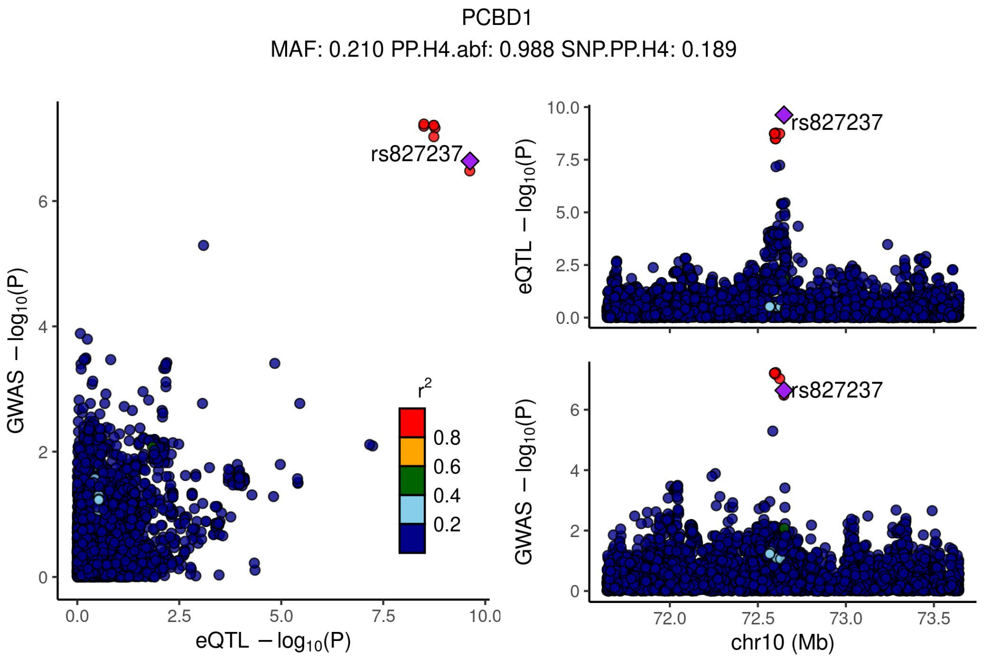


Supplementary Figure 4.38 - Locus compare plots 1Mbp up or down the TSS of PCBD1.

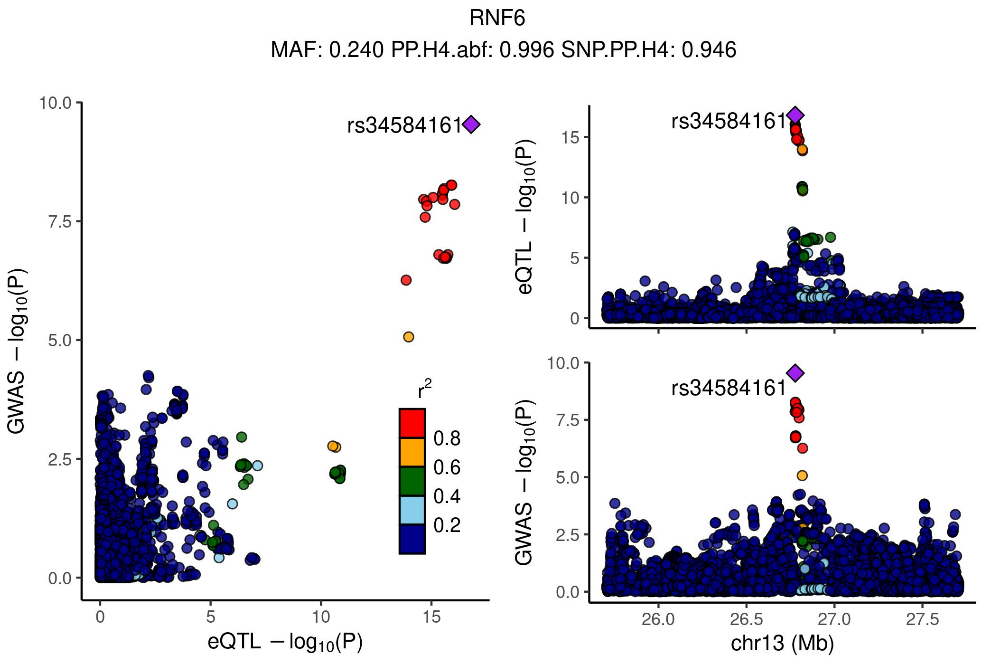


Supplementary Figure 4.39 - Locus compare plots 1Mbp up or down the TSS of RNF6.

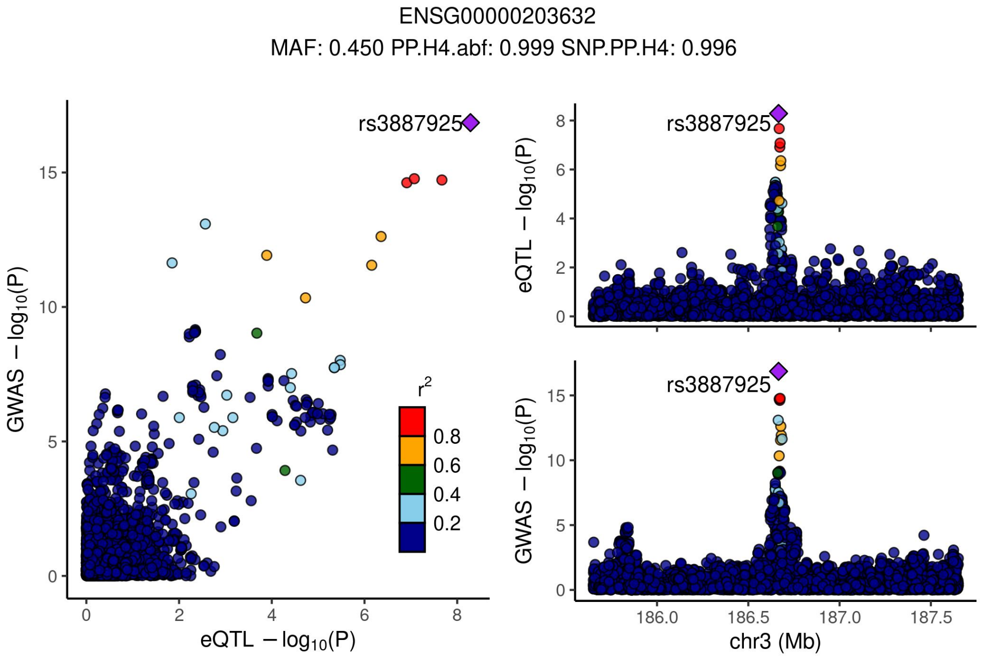


Supplementary Figure 4.40 - Locus compare plots 1Mbp up or down the TSS of ENSG00000203632.

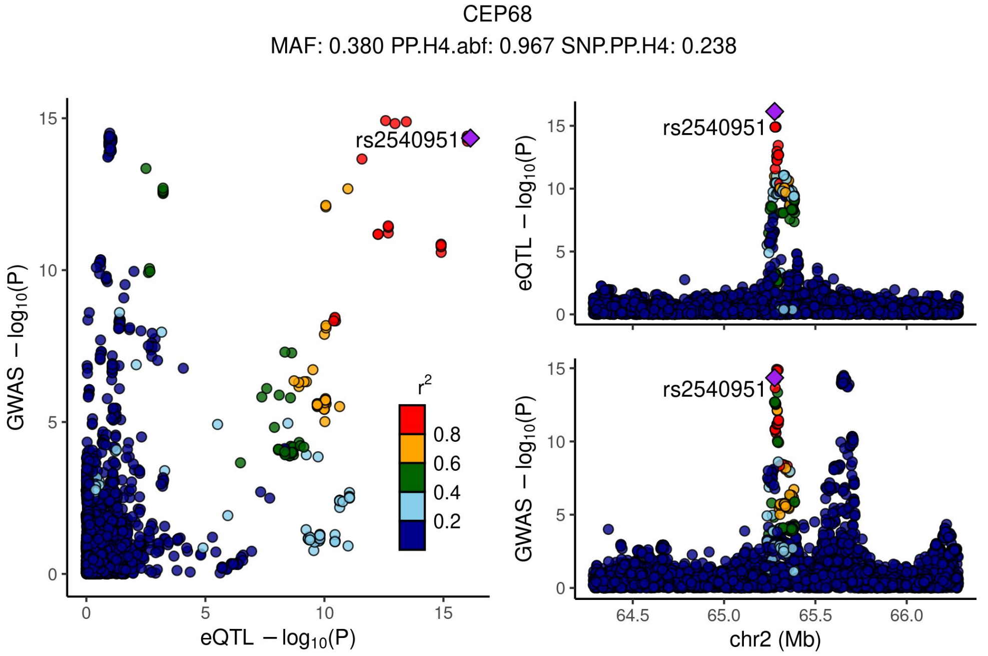


Supplementary Figure 4.41 - Locus compare plots 1Mbp up or down the TSS of CEP68.

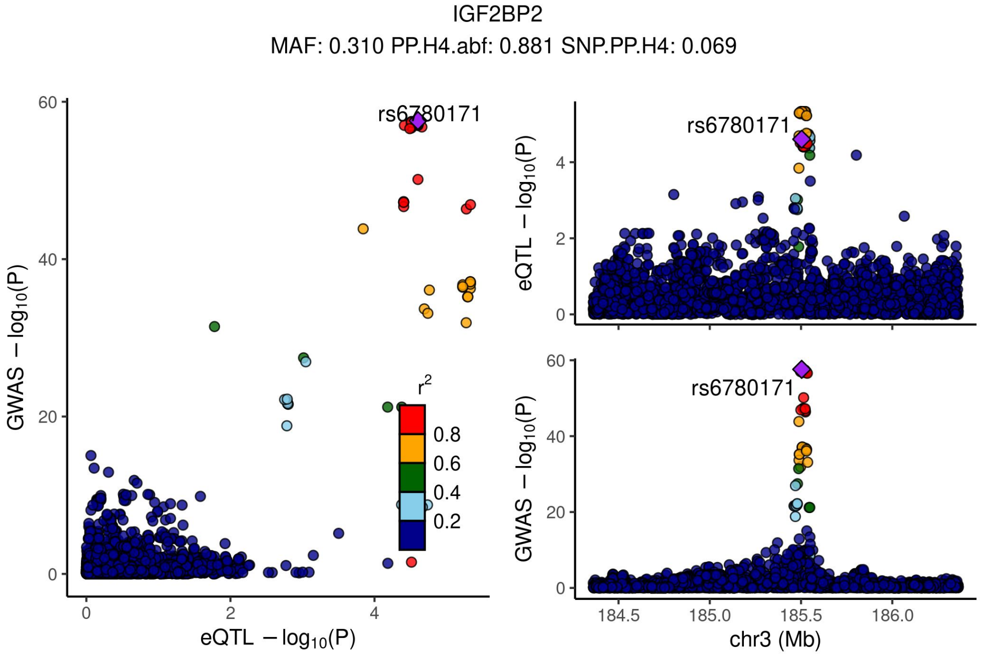


Supplementary Figure 4.42 - Locus compare plots 1Mbp up or down the TSS of IGF2BP2.

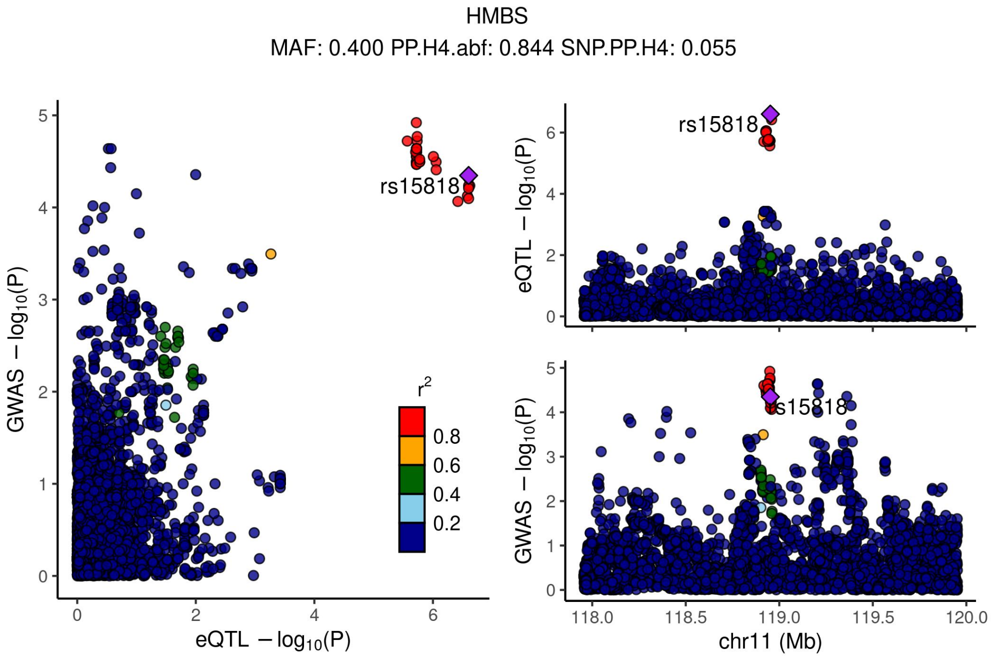


Supplementary Figure 4.43 - Locus compare plots 1Mbp up or down the TSS of HMBS.

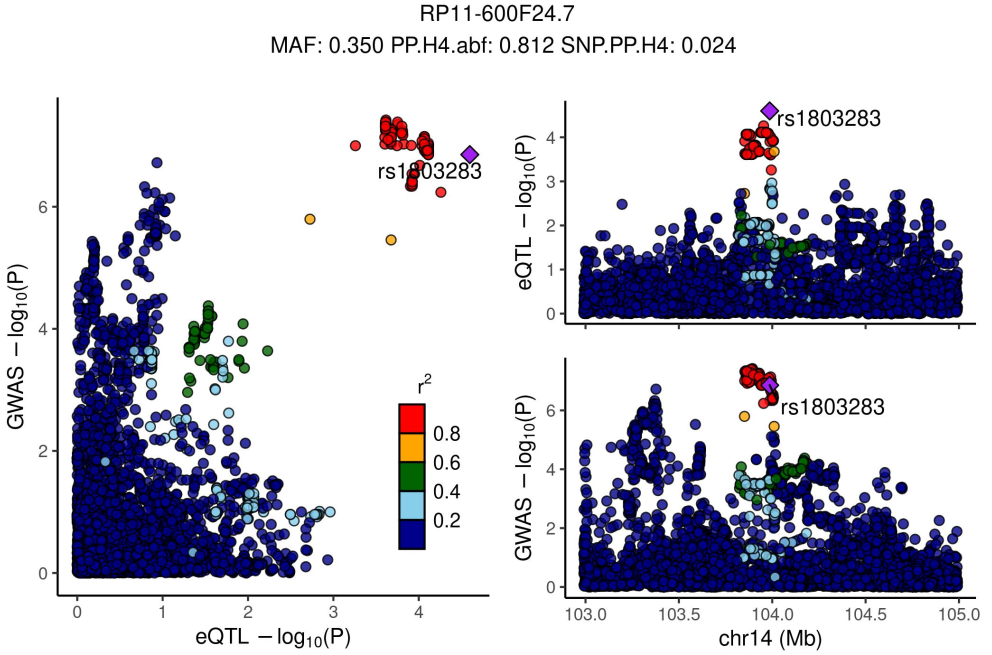


Supplementary Figure 4.44 - Locus compare plots 1Mbp up or down the TSS of RP11-600F24.7.

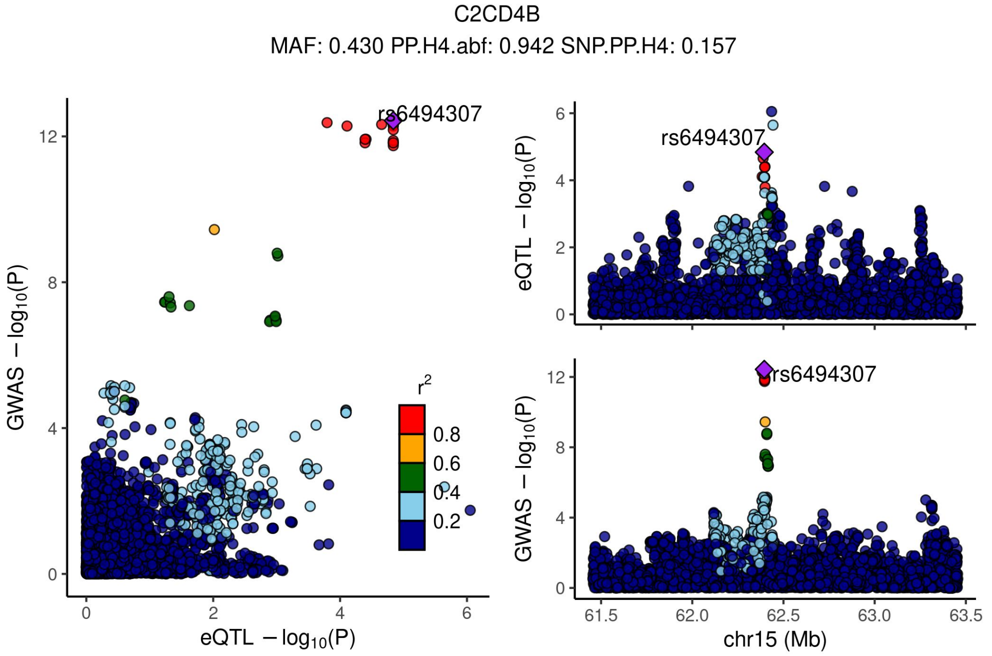


Supplementary Figure 4.45 - Locus compare plots 1Mbp up or down the TSS of C2CD4B.

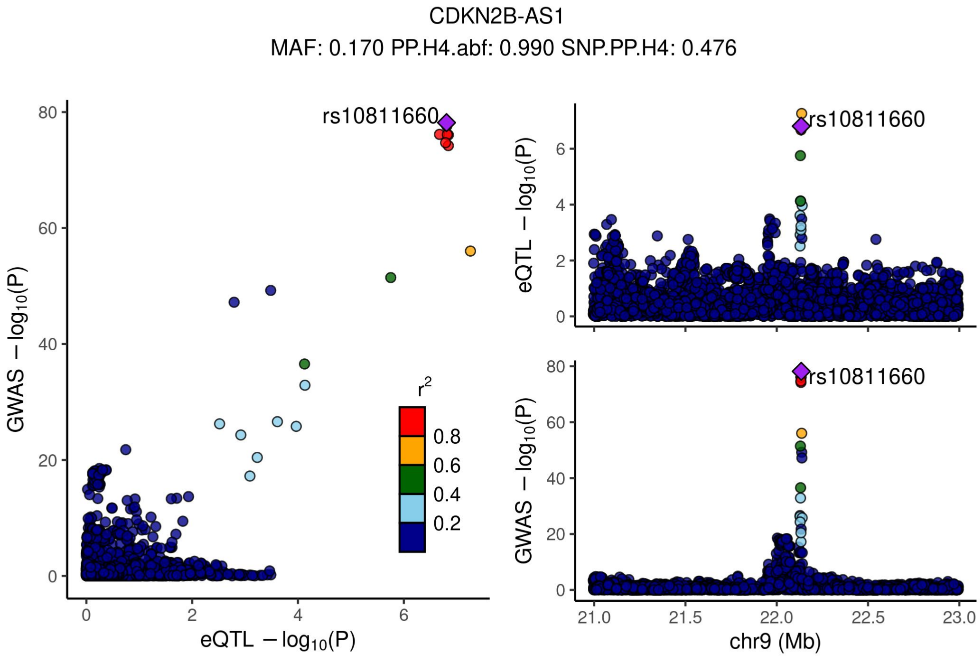


Supplementary Figure 4.46 - Locus compare plots 1Mbp up or down the TSS of CDKN2B-AS1.

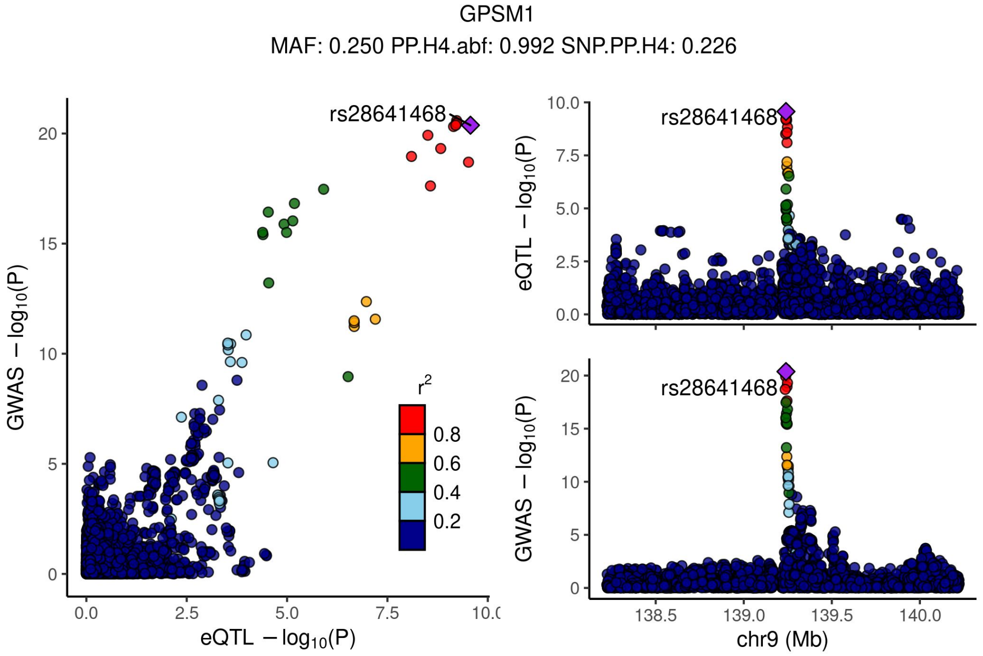


Supplementary Figure 4.47 - Locus compare plots 1Mbp up or down the TSS of GPSM1.

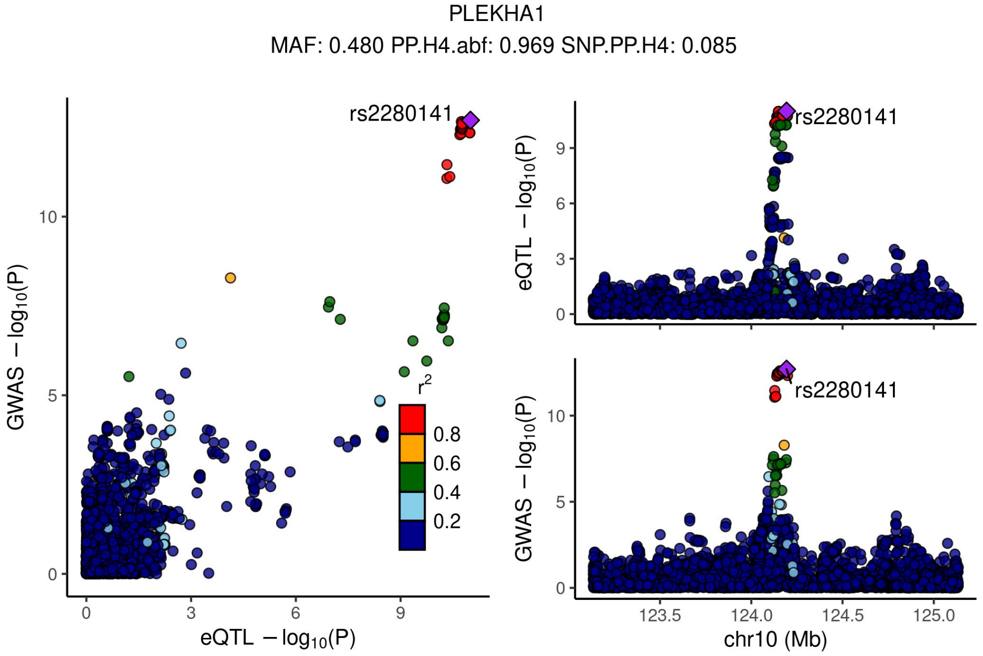


Supplementary Figure 4.48 - Locus compare plots 1Mbp up or down the TSS of PLEKHA1.

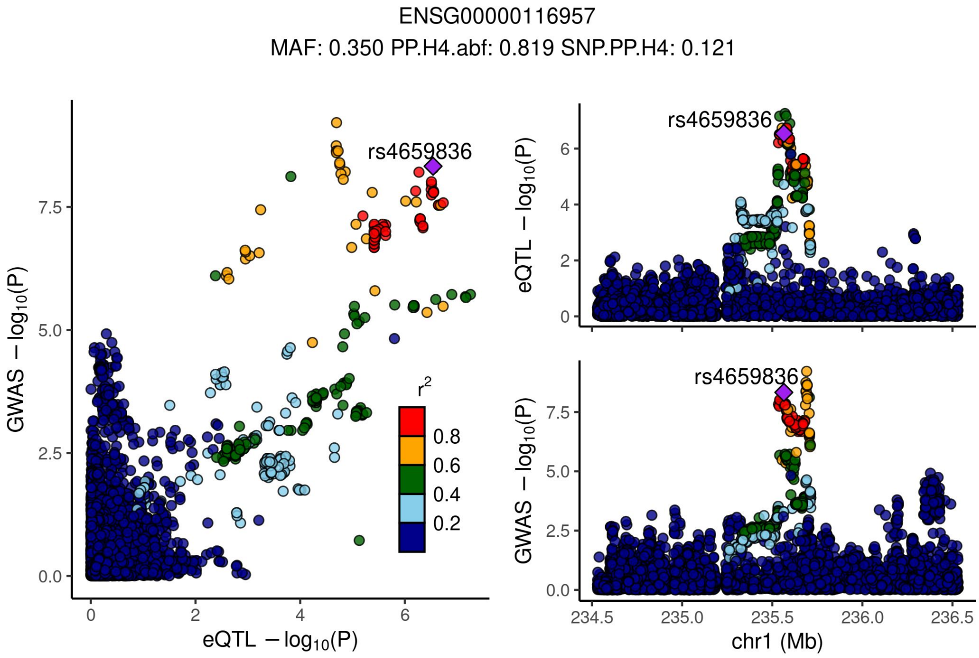


Supplementary Figure 4.49 - Locus compare plots 1Mbp up or down the TSS of ENSG00000116957.

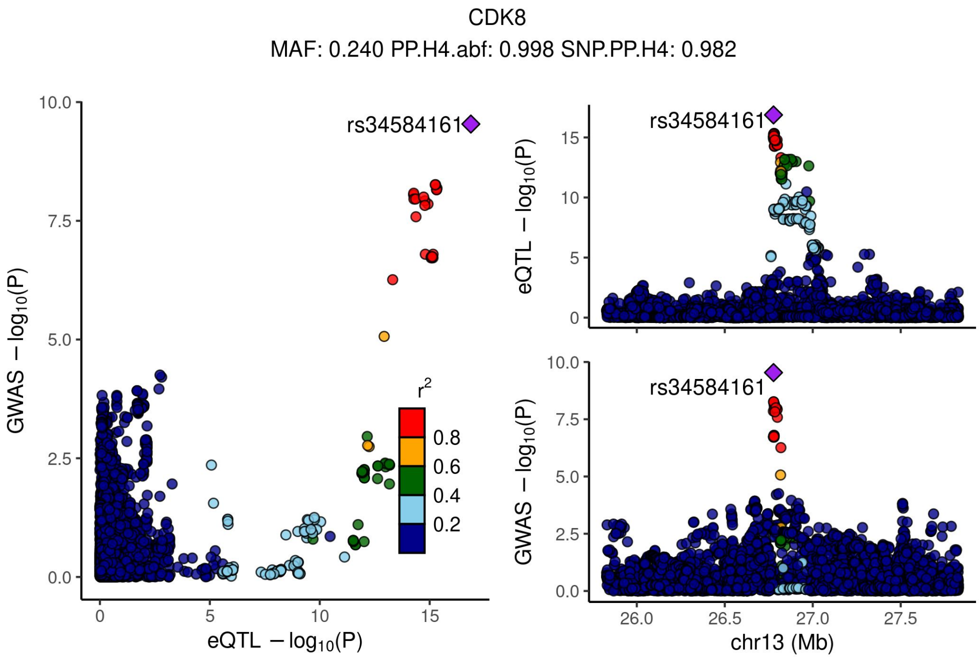


Supplementary Figure 4.50 - Locus compare plots 1Mbp up or down the TSS of CDK8.

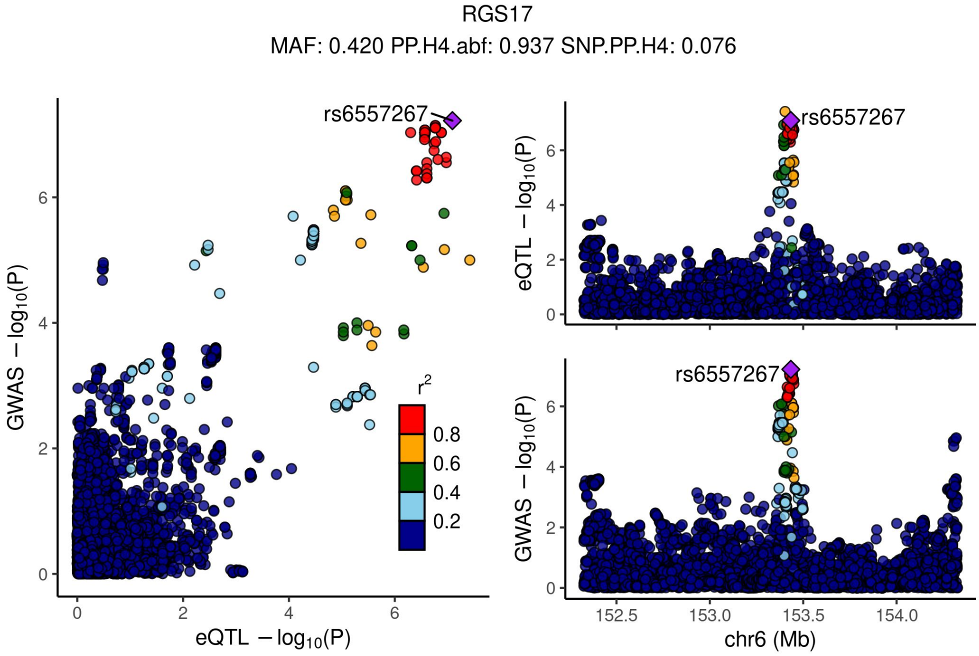


Supplementary Figure 4.51 - Locus compare plots 1Mbp up or down the TSS of RGS17.

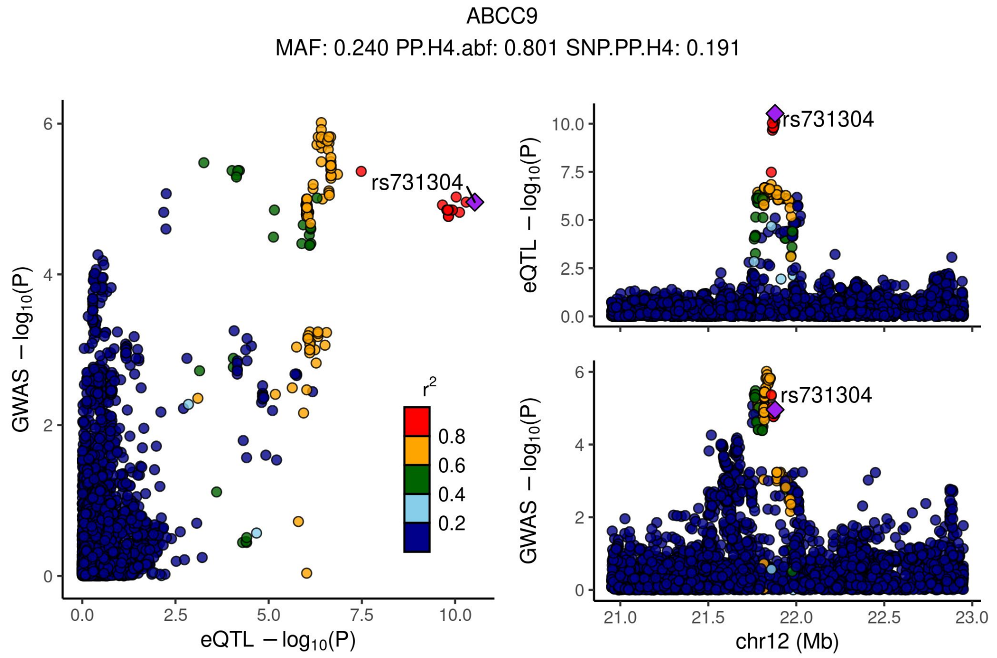


Supplementary Figure 4.52 - Locus compare plots 1Mbp up or down the TSS of ABCC9.

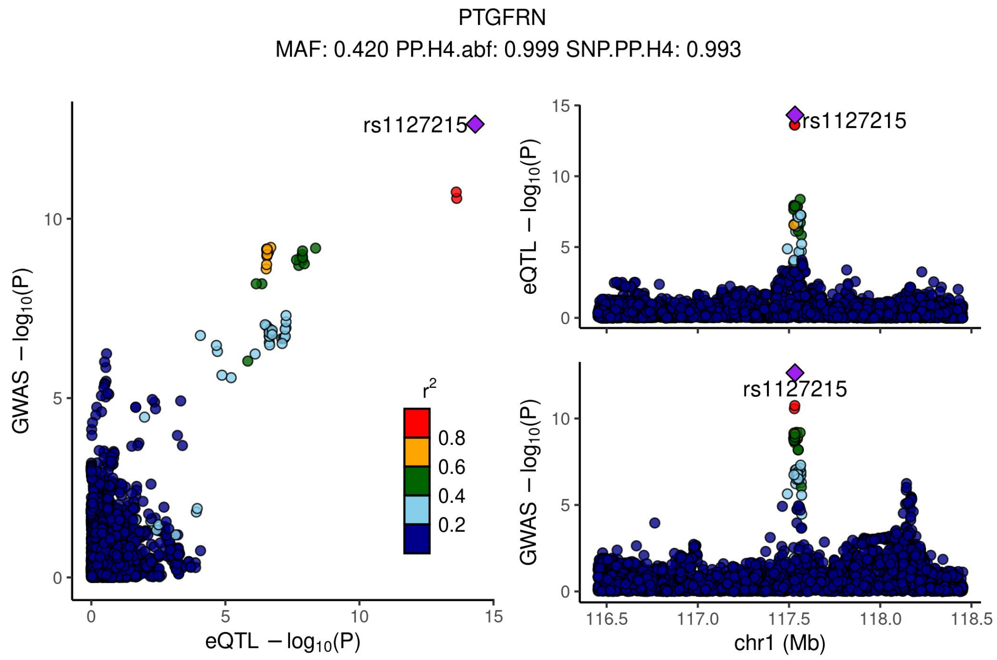


Supplementary Figure 4.53 - Locus compare plots 1Mbp up or down the TSS of PTGFRN.
