## Supplemental Figure 8 for "TIGER: The gene expression regulatory variation landscape of human pancreatic islets"

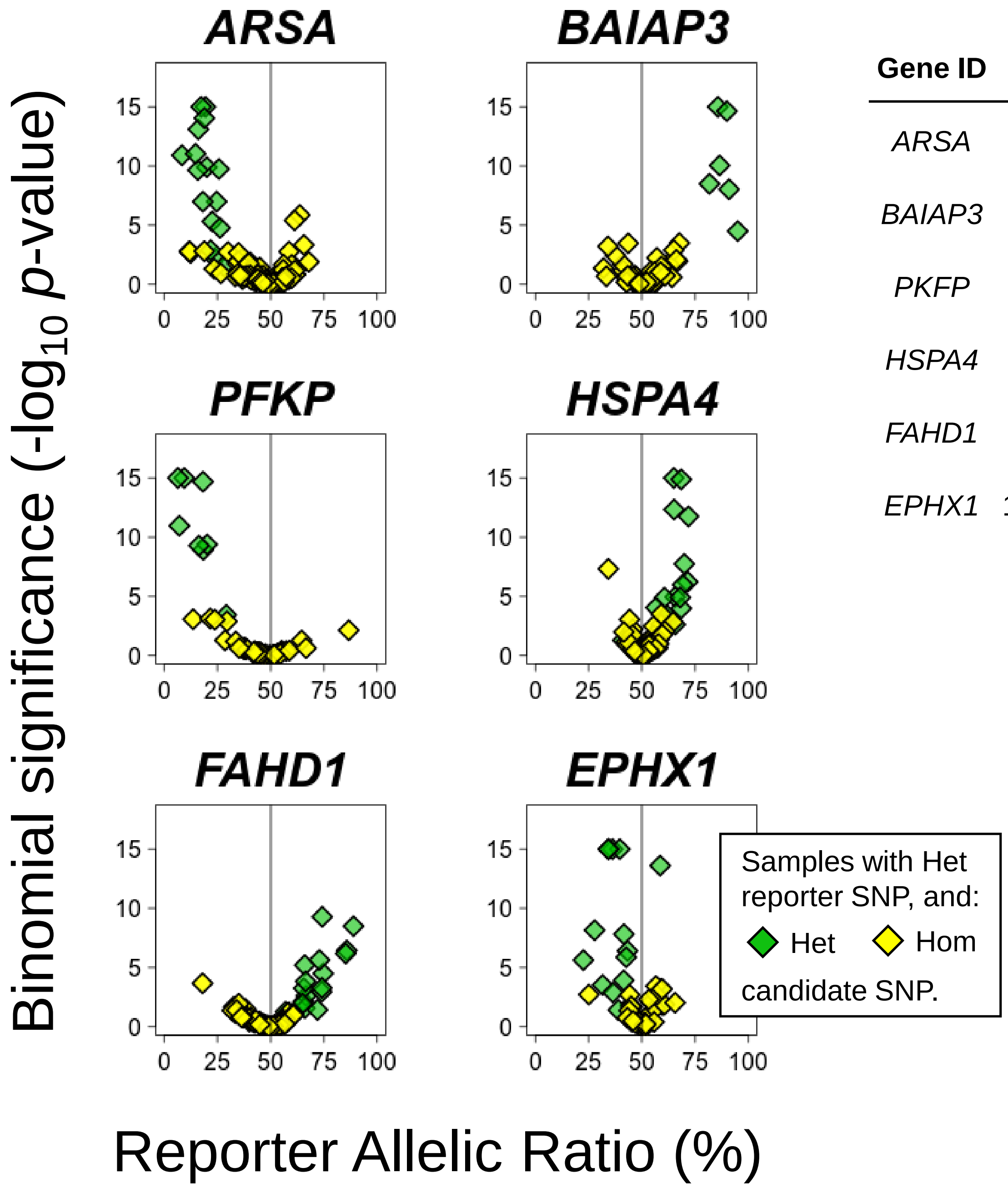


| Gene ID | Candidate variant | Reporter variant | Candidate Z-score | Candidate *p*-value | Reporter Z-score | Reporter *p*-value |
|---|---|---|---|---|---|---|
| *ARSA* | 22:51060049_G_A | 22:51064039_G_C | -23.4568 | 6.99E-20 | -7.6403 | 0.002 |
| *BAIAP3* | 16:1385756_C_T | 16:1398818_G_C | 15.5988 | 8.73E-10 | 5.89466 | 0.002 |
| *PKFP* | 10:4301860_A_G | 10:3163509_A_G | -15.4788 | 1.16E-09 | -4.86632 | 0.006 |
| *HSPA4* | 5:132444128_G_A | 5:132437531_C_T | 15.0939 | 2.85E-09 | 7.33077 | 0.011 |
| *FAHD1* | 16:1919271_G_A | 16:1877913_C_G | 15.0232 | 3.36E-09 | 6.5349 | 0.021 |
| *EPHX1* | 1:225989641_CA_C | 1:226032229_C_T | -13.9759 | 3.45E-08 | -5.25428 | 0.030 |
