## Supplemental Figure 9 for "TIGER: The gene expression regulatory variation landscape of human pancreatic islets"

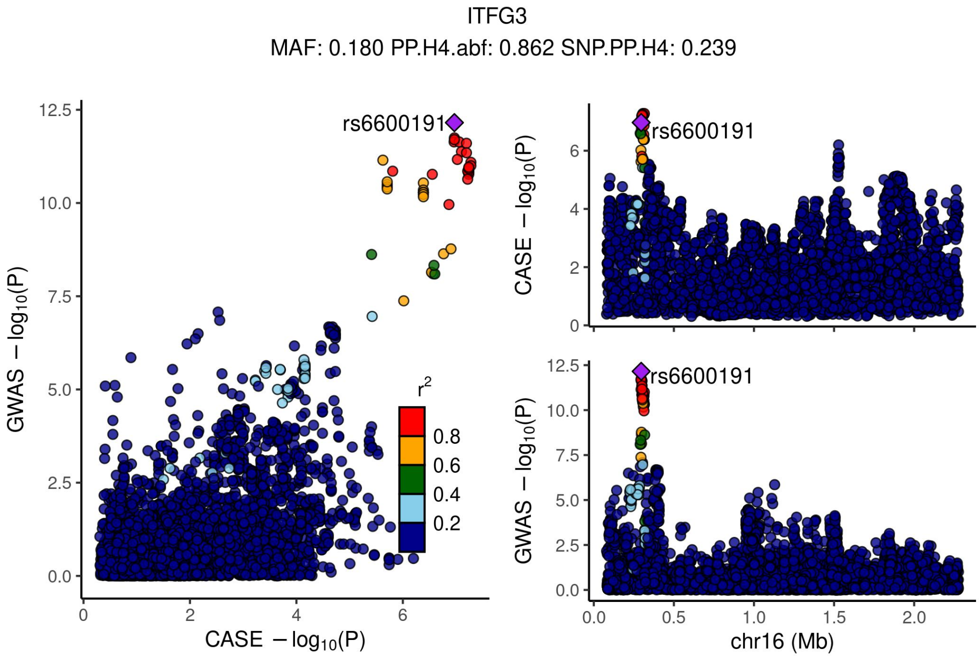


Supplementary Figure 11.1 - Locus compare plots 1Mbp up or down the TSS of ITGF3.

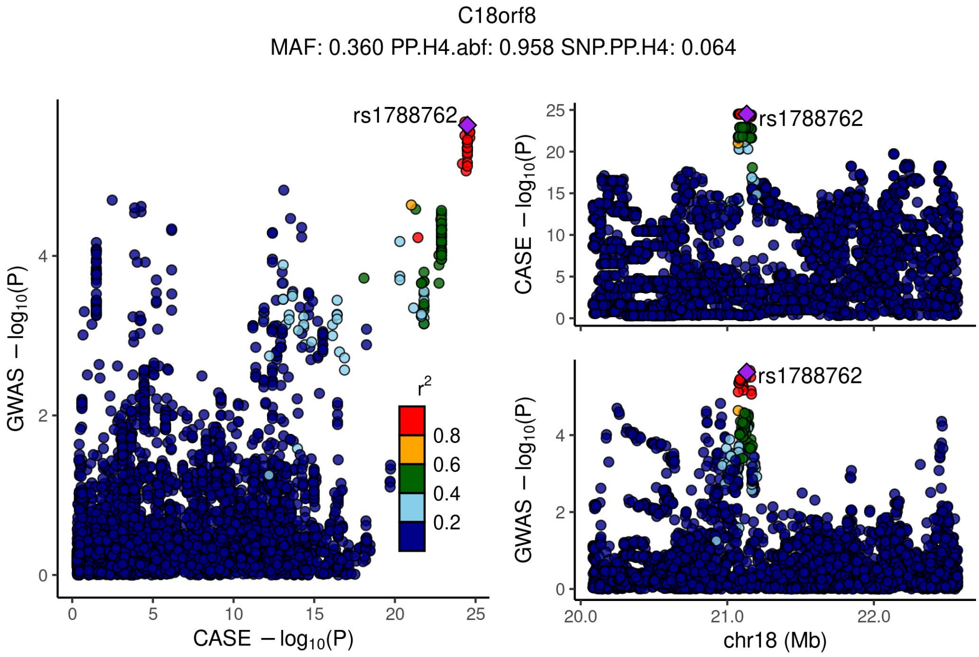


Supplementary Figure 11.2 - Locus compare plots 1Mbp up or down the TSS of C18orf8.

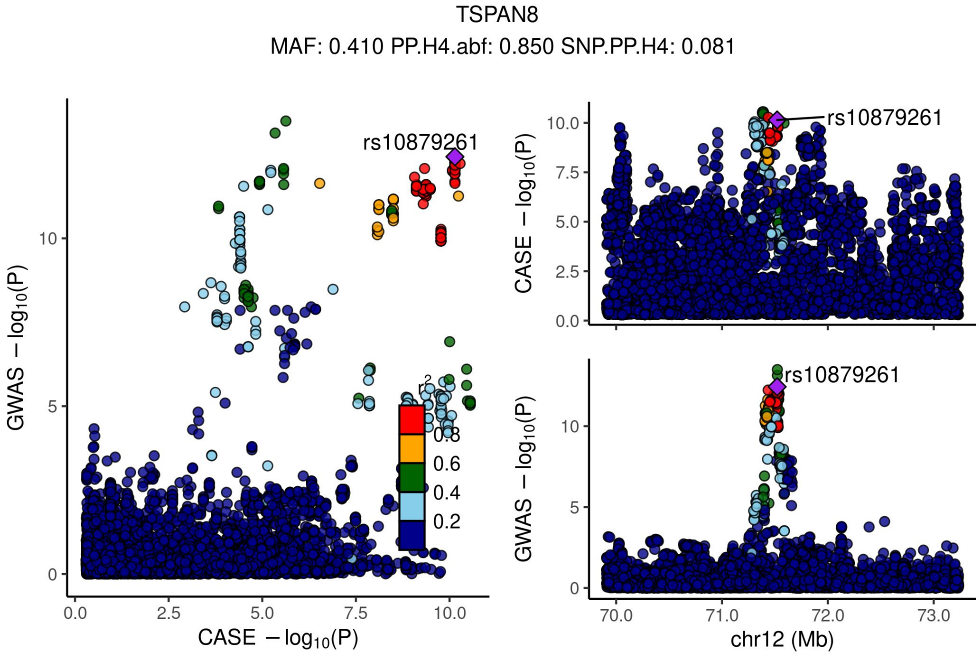


Supplementary Figure 11.3 - Locus compare plots 1Mbp up or down the TSS of TSPAN8.

RP11-613D13.5
MAF: 0.290 PP.H4.abf: 0.930 SNP.PP.H4: 0.074

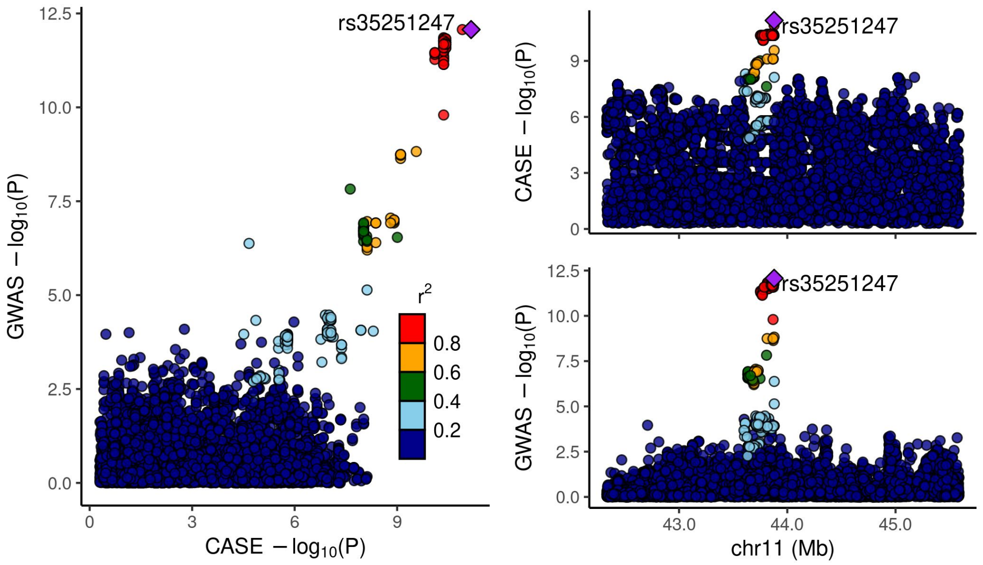


Supplementary Figure 11.4 - Locus compare plots 1Mbp up or down the TSS of RP11-613D13.5.

FXYD2
MAF: 0.480 PP.H4.abf: 0.945 SNP.PP.H4: 1.000

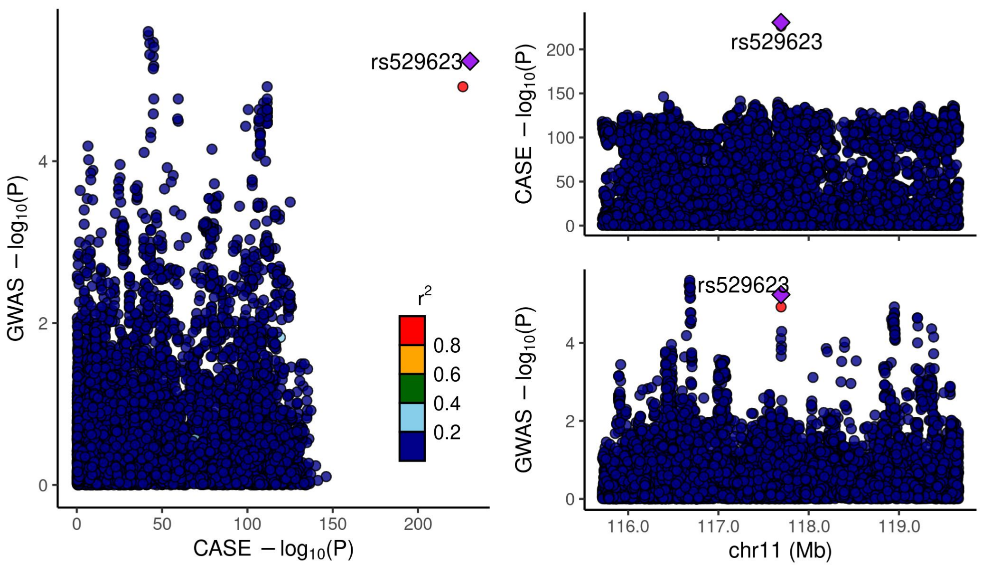


Supplementary Figure 11.5 - Locus compare plots 1Mbp up or down the TSS of FXYD2.

CALR
MAF: 0.410 PP.H4.abf: 0.994 SNP.PP.H4: 0.469

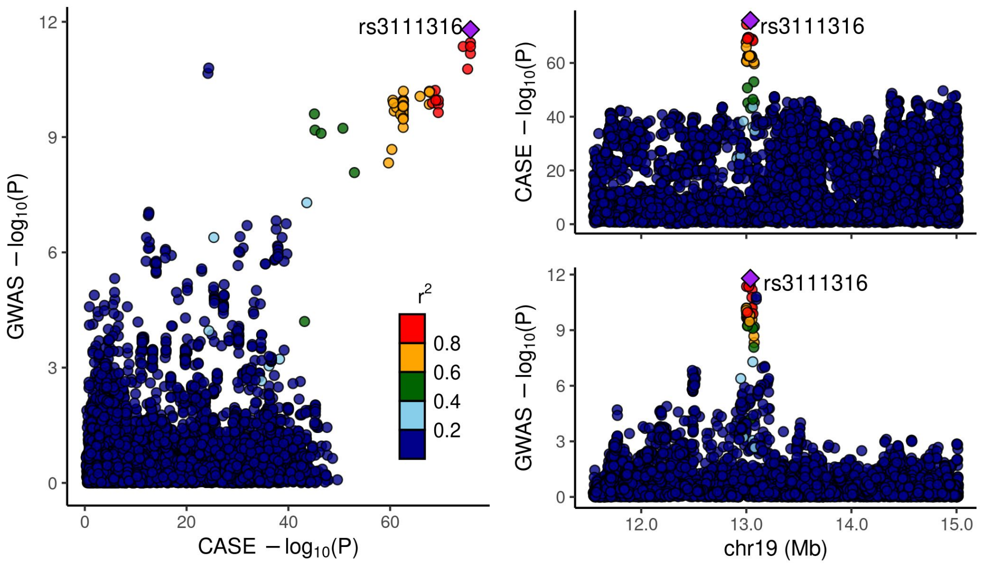


Supplementary Figure 11.6 - Locus compare plots 1Mbp up or down the TSS of CALR.

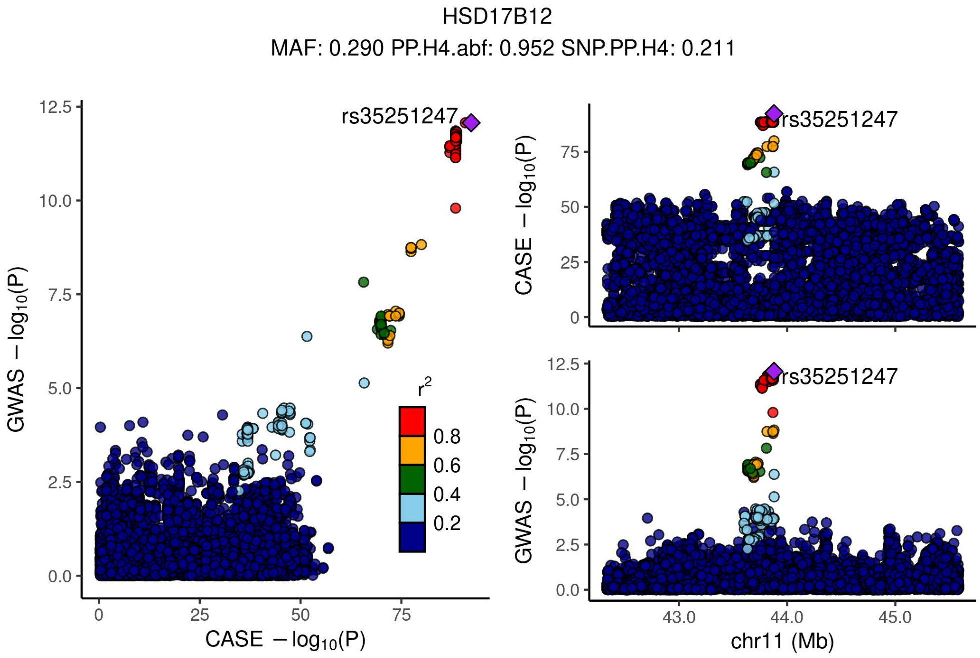


Supplementary Figure 11.7 - Locus compare plots 1Mbp up or down the TSS of HSD17B12.

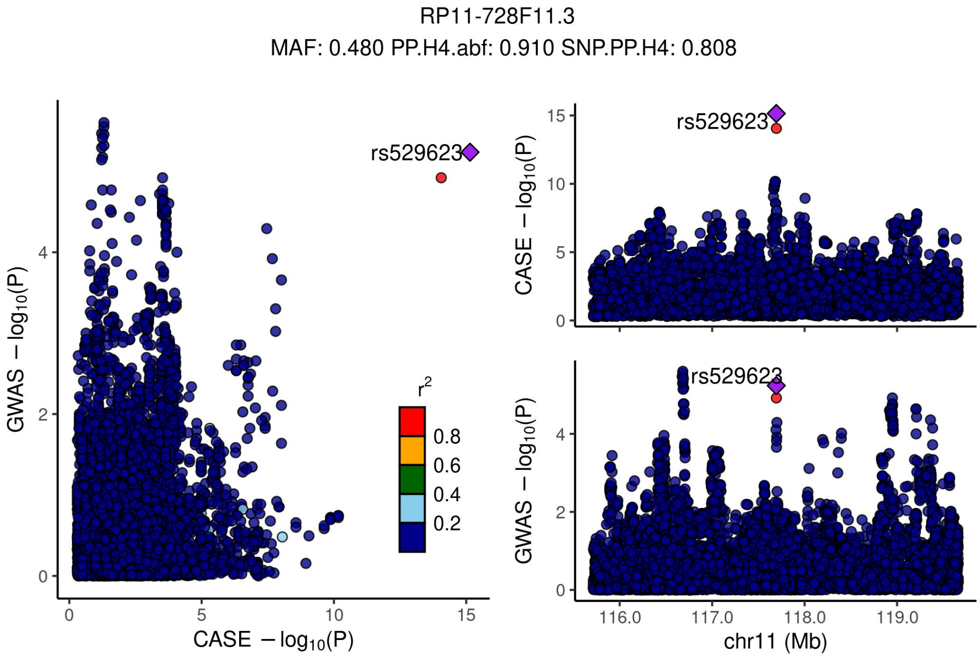


Supplementary Figure 11.8 - Locus compare plots 1Mbp up or down the TSS of RP11-728F11.3.

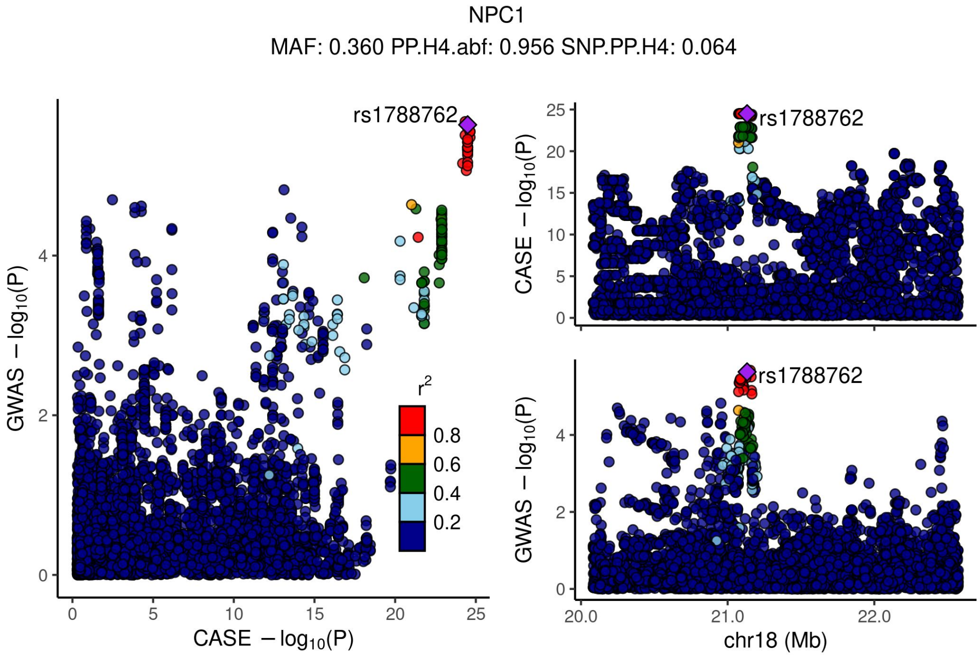


Supplementary Figure 11.9 - Locus compare plots 1Mbp up or down the TSS of NPC1.

KCNJ11
MAF: 0.370 PP.H4.abf: 0.832 SNP.PP.H4: 0.361

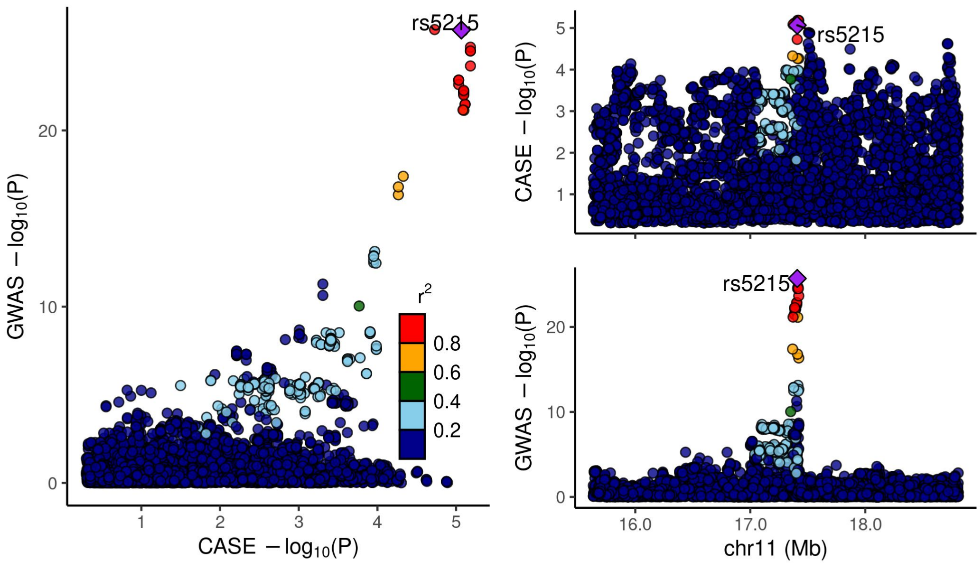


Supplementary Figure 11.10 - Locus compare plots 1Mbp up or down the TSS of KCNJ11.

PLEKHA1
MAF: 0.480 PP.H4.abf: 0.955 SNP.PP.H4: 0.060

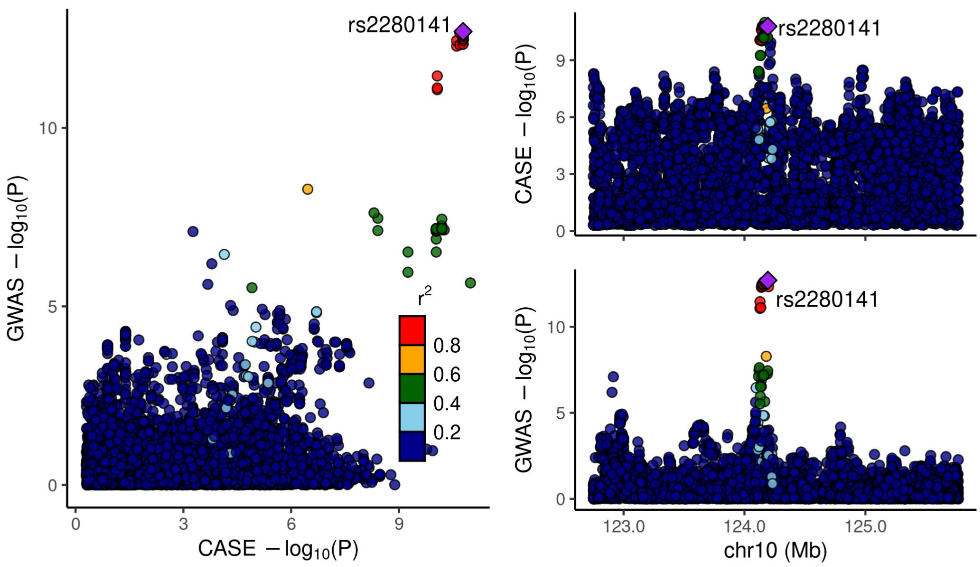


Supplementary Figure 11.11 - Locus compare plots 1Mbp up or down the TSS of PLEKHA1.

SLC30A8
MAF: 0.310 PP.H4.abf: 0.996 SNP.PP.H4: 0.607

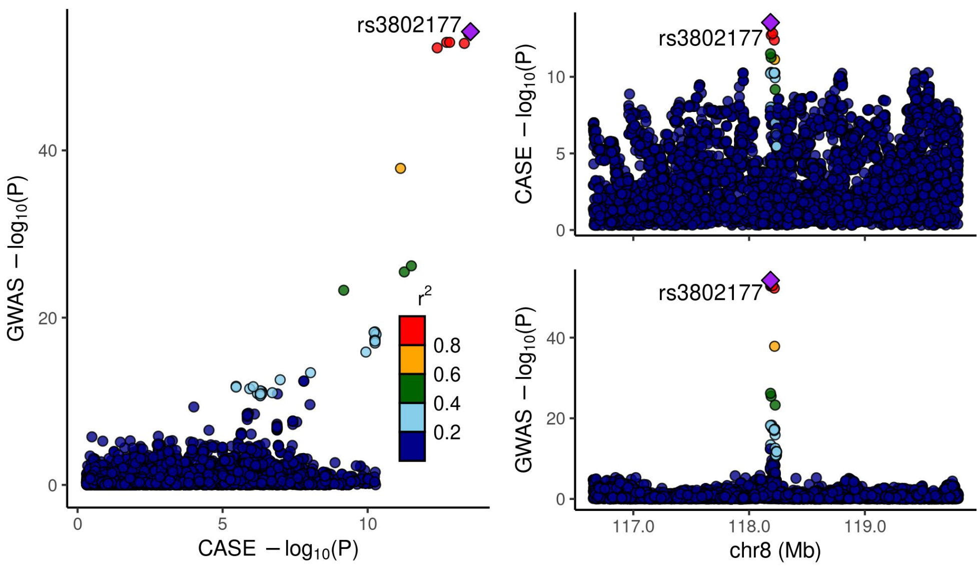


Supplementary Figure 11.12 - Locus compare plots 1Mbp up or down the TSS of SLC30A8.

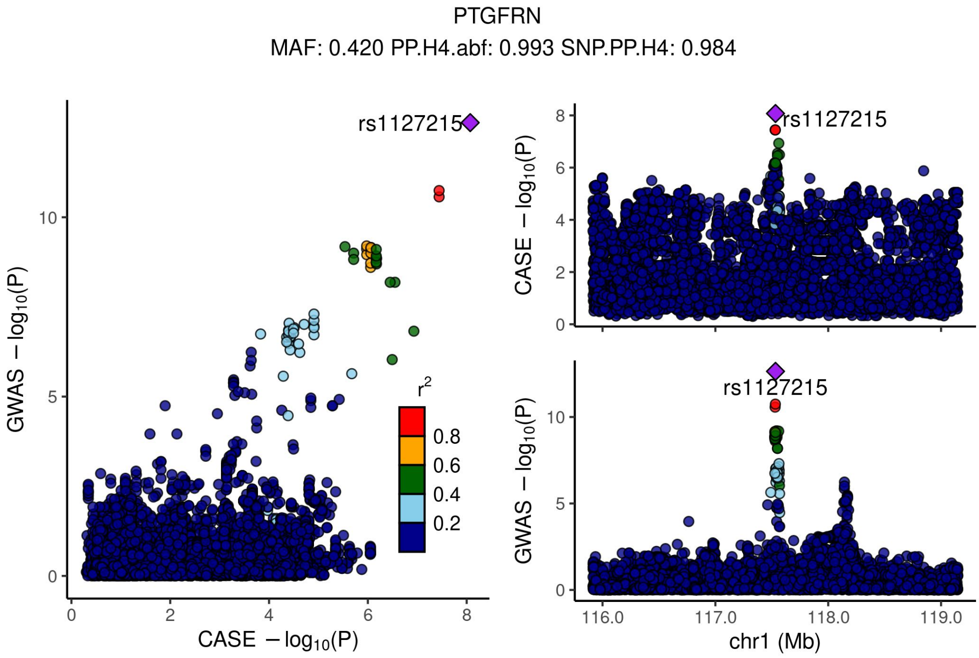


Supplementary Figure 11.13 - Locus compare plots 1Mbp up or down the TSS of PTGFRN.

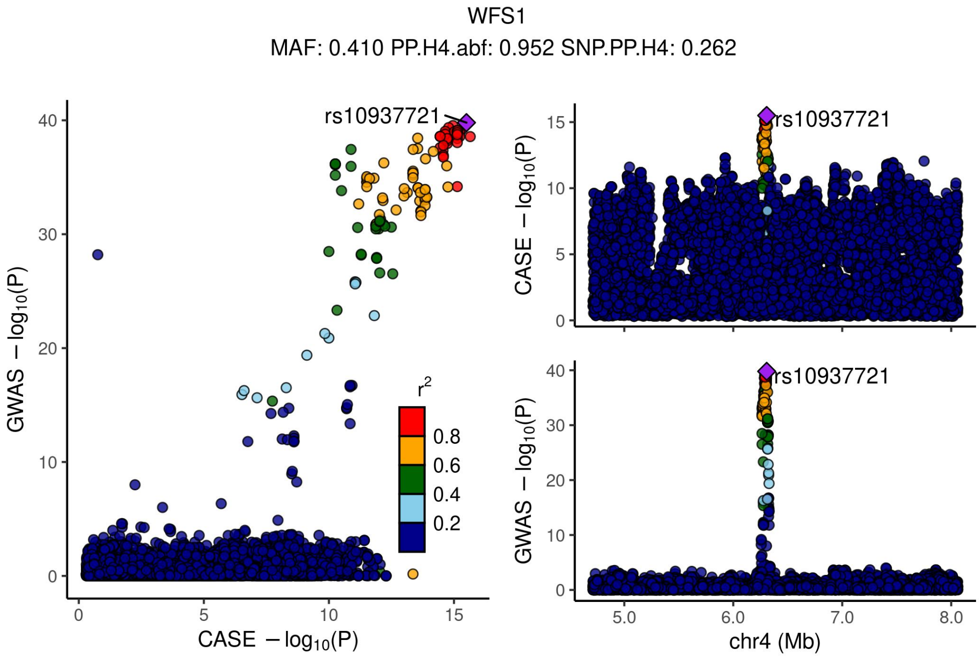


Supplementary Figure 11.14 - Locus compare plots 1Mbp up or down the TSS of WFS1.
