## Supplemental Figure 10 for "TIGER: The gene expression regulatory variation landscape of human pancreatic islets"

### BATCH EFFECTS - limma log QCed (>80% NAs removed)

Sample correlation on gene expression for all genes and all samples. **Before** correcting for **batch effects** in cohorts.

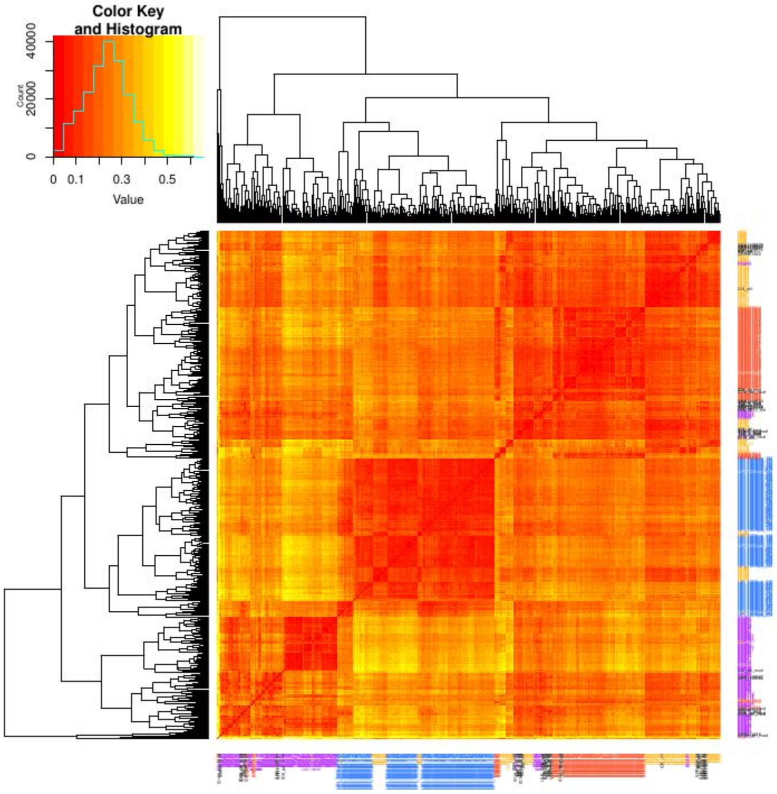


Sample correlation on gene expression for all genes and all samples. **After** correcting for **batch effects** in cohorts **and** considering bmi, age, gender, diabetic type as **covariates**.

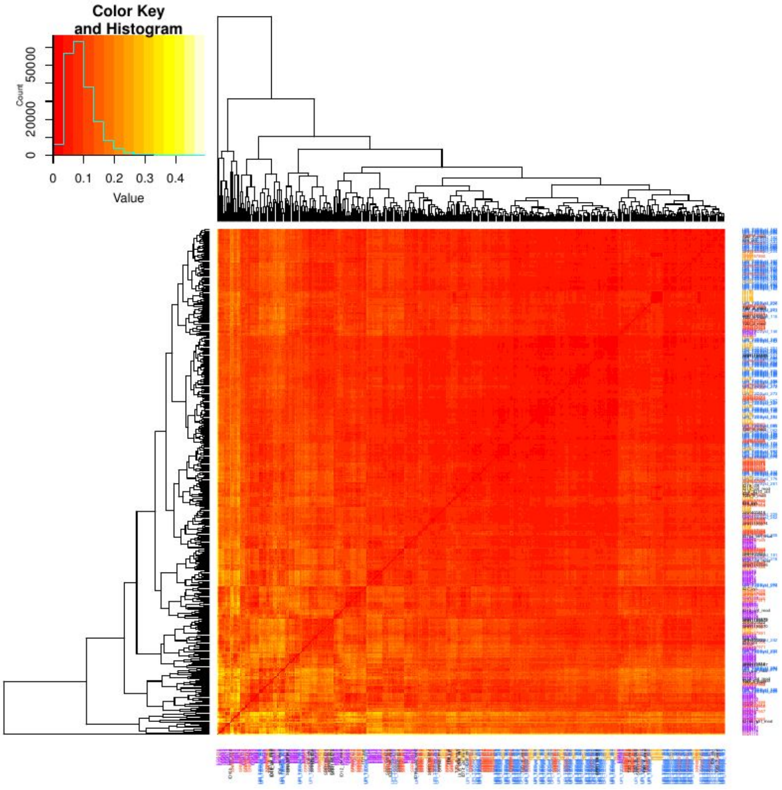


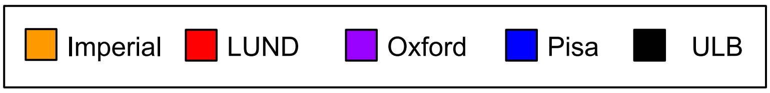
