## Appendix MAGIC Authors for "TIGER: The gene expression regulatory variation landscape of human pancreatic islets"

**Meta-Analysis of Glucose and Insulin-related Traits Consortium (MAGIC)**

Ji Chen^1,2#^, Cassandra N. Spracklen^3,4#^, Gaëlle Marenne^2,5#^, Arushi Varshney^6#^, Laura J Corbin^7,8#^, Jian’an Luan^9^, Sara M Willems^9^, Ying Wu^3^, Xiaoshuai Zhang^9,10^, Momoko Horikoshi^11,12,13^, Thibaud S Boutin^14^, Reedik Mägi^15^, Johannes Waage^16^, Ruifang Li-Gao^17^, Kei Hang Katie Chan^18,19,20^, Jie Yao^21^, Mila D Anasanti^22^, Audrey Y Chu^23^, Annique Claringbould^24^, Jani Heikkinen^22^, Jaeyoung Hong^25^, Jouke-Jan Hottenga^26,27^, Shaofeng Huo^28^, Marika A. Kaakinen^29,22^, Tin Louie^30^, Winfried März^31,32,33^, Hortensia Moreno-Macias^34^, Anne Ndungu^12^, Sarah C. Nelson^30^, Ilja M. Nolte^35^, Kari E North^36^, Chelsea K. Raulerson^3^, Debashree Ray^37^, Rebecca Rohde^36^, Denis Rybin^25^, Claudia Schurmann^38,39^, Xueling Sim^40,41,42^, Loz Southam^2^, Isobel D Stewart^9^, Carol A. Wang^43^, Yujie Wang^36^, Peitao Wu^25^, Weihua Zhang^44,45^, Tarunveer S. Ahluwalia^16,46,47^, Emil VR Appel^48^, Lawrence F. Bielak^49^, Jennifer A. Brody^50^, Noël P Burtt^51^, Claudia P Cabrera^52,53^, Brian E Cade^54,55^, Jin Fang Chai^40^, Xiaoran Chai^56,57^, Li-Ching Chang^58^, Chien-Hsiun Chen^58^, Brian H Chen^59^, Kumaraswamy Naidu Chitrala^60^, Yen-Feng Chiu^61^, Hugoline G. de Haan^17^, Graciela E Delgado^33^, Ayse Demirkan^62,29^, Qing Duan^3,63^, Jorgen Engmann^64^, Segun A Fatumo^65,66,67^, Javier Gayán^68^, Franco Giulianini^69^, Jung Ho Gong^18^, Stefan Gustafsson^70^, Yang Hai^71^, Fernando P Hartwig^72,7^, Jing He^73^, Yoriko Heianza^74^, Tao Huang^75^, Alicia Huerta-Chagoya^76,77^, Mi Yeong Hwang^78^, Richard A. Jensen^50^, Takahisa Kawaguchi^79^, Katherine A Kentistou^80,81^, Young Jin Kim^78^, Marcus E Kleber^33^, Ishminder K Kooner^45^, Shuiqing Lai^18^, Leslie A Lange^82^, Carl D Langefeld^83^, Marie Lauzon^21^, Man Li^84^, Symen Ligthart^62^, Jun Liu^62,85^, Marie Loh^86,44^, Jirong Long^87^, Valeriya Lyssenko^88,89^, Massimo Mangino^90,91^, Carola Marzi^92,93^, May E Montasser^94^, Abhishek Nag^12^, Masahiro Nakatochi^95^, Damia Noce^96^, Raymond Noordam^97^, Giorgio Pistis^98^, Michael Preuss^38,99^, Laura Raffield^3^, Laura J. Rasmussen-Torvik^100^, Stephen S Rich^101,102^, Neil R Robertson^11,12^, Rico Rueedi^103,104^, Kathleen Ryan^94^, Serena Sanna^98,24^, Richa Saxena^105,106,107^, Katharina E Schraut^80,81^, Bengt Sennblad^108^, Kazuya Setoh^79^, Albert V Smith^109,110^, Lorraine Southam^111,112^, Thomas Sparsø^48^, Rona J Strawbridge^113,114^, Fumihiko Takeuchi^115^, Jingyi Tan^21^, Stella Trompet^97,116^, Erik van den Akker^117,118,119^, Peter J van der Most^35^, Niek Verweij^120,121^, Mandy Vogel^122^, Heming Wang^54,55^, Chaolong Wang^123,124^, Nan Wang^125,126^, Helen R Warren^52,53^, Wanqing Wen^87^, Tom Wilsgaard^127^, Andrew Wong^128^, Andrew R Wood^1^, Tian Xie^35^, Mohammad Hadi Zafarmand^129,130^, Jing-Hua Zhao^131^, Wei Zhao^49^, Najaf Amin^62,85^, Zorayr Arzumanyan^21^, Arne Astrup^132^, Stephan JL Bakker^133^, Damiano Baldassarre^134,135^, Marian Beekman^117^, Richard N Bergman^136^, Alain Bertoni^137^, Matthias Blüher^138^, Lori L. Bonnycastle^139^, Stefan R Bornstein^140^, Donald W Bowden^141^, Qiuyin Cai^73^, Archie Campbell^142,143^, Harry Campbell^80^, Yi Cheng Chang^144,145,146^, Eco J.C. de Geus^26,27^, Abbas Dehghan^62^, Shufa Du^147^, Gudny Eiriksdottir^110^, Aliki Eleni Farmaki^148,149^, Mattias Frånberg^150^, Christian Fuchsberger^96^, Yutang Gao^151^, Anette P Gjesing^48^, Anuj Goel^152,12^, Sohee Han^78^, Catharina A Hartman^153^, Christian Herder^154,155,156^, Andrew A. Hicks^96^, Chang-Hsun Hsieh^157,158^, Willa A. Hsueh^159^, Sahoko Ichihara^160^, Michiya Igase^161^, M. Arfan Ikram^62^, W. Craig Johnson^30^, Marit E Jørgensen^46,162^, Peter K Joshi^80^, Rita R Kalyani^163^, Fouad R. Kandeel^164^, Tomohiro Katsuya^165,166^, Chiea Chuen Khor^124^, Wieland Kiess^122^, Ivana Kolcic^167^, Teemu Kuulasmaa^168^, Johanna Kuusisto^169^, Kristi Läll^15^, Kelvin Lam^21^, Deborah A Lawlor^170,8^, Nanette R. Lee^171,172^, Rozenn N. Lemaitre^50^, Honglan Li^173^, Shih-Yi Lin^174,175,176^, Jaana Lindström^177^, Allan Linneberg^178,179^, Jianjun Liu^124,180^, Carlos Lorenzo^181^, Tatsuaki Matsubara^182^, Fumihiko Matsuda^79^, Geltrude Mingrone^183^, Simon Mooijaart^97^, Sanghoon Moon^78^, Toru Nabika^184^, Girish N. Nadkarni^38^, Jerry L. Nadler^185^, Mari Nelis^15^, Matt J Neville^11,186^, Jill M Norris^187^, Yasumasa Ohyagi^188^, Annette Peters^189,93,190^, Patricia A. Peyser^49^, Ozren Polasek^167,191^, Qibin Qi^192^, Dennis Raven^153^, Dermot F Reilly^193^, Alex Reiner^194^, Fernando Rivideneira^195^, Kathryn Roll^21^, Igor Rudan^196^, Charumathi Sabanayagam^56,197^, Kevin Sandow^21^, Naveed Sattar^198^, Annette Schürmann^199,200^, Jinxiu Shi^201^, Heather M Stringham^42,41^, Kent D. Taylor^21^, Tanya M. Teslovich^202^, Betina Thuesen^178^, Paul RHJ Timmers^80,203^, Elena Tremoli^135^, Michael Y Tsai^204^, Andre Uitterlinden^195^, Rob M van Dam^40,180,205^, Diana van Heemst^97^, Astrid van Hylckama Vlieg^17^, Jana V Van Vliet-Ostaptchouk^35^, Jagadish Vangipurapu^206^, Henrik Vestergaard^48,207^, Tao Wang^192^, Ko Willems van Dijk^208,209,210^, Tatijana Zemunik^211^, Goncalo R Abecasis^42^, Linda S. Adair^147,212^, Carlos Alberto Aguilar-Salinas^213,214,215^, Marta E Alarcón-Riquelme^216,217^, Ping An^218^, Larissa Aviles-Santa^219^, Diane M Becker^220^, Lawrence J Beilin^221^, Sven Bergmann^103,104,222^, Hans Bisgaard^16^, Corri Black^223^, Michael Boehnke^42,41^, Eric Boerwinkle^224,225^, Bernhard O Böhm^226,227^, Klaus Bønnelykke^16^, D I. Boomsma^26,27^, Erwin P. Bottinger^38,228,229^, Thomas A Buchanan^230,231,126^, Mickaël Canouil^232,233^, Mark J Caulfield^52,53^, John C. Chambers^86,44,45,234,235^, Daniel I. Chasman^69,236^, Yii-Der Ida Chen^21^, Ching-Yu Cheng^56,197^, Francis S. Collins^139^, Adolfo Correa^237^, Francesco Cucca^98^, H. Janaka de Silva^238^, George Dedoussis^239^, Sölve Elmståhl^240^, Michele K. Evans^241^, Ele Ferrannini^242^, Luigi Ferrucci^243^, Jose C Florez^244,245,107^, Paul W Franks^89,246^, Timothy M Frayling^1^, Philippe Froguel^232,233,247^, Bruna Gigante^248^, Mark O. Goodarzi^249^, Penny Gordon-Larsen^147,212^, Harald Grallert^92,93^, Niels Grarup^48^, Sameline Grimsgaard^127^, Leif Groop^250,251^, Vilmundur Gudnason^110,252^, Xiuqing Guo^21^, Anders Hamsten^114^, Torben Hansen^48^, Caroline Hayward^203^, Susan R. Heckbert^253^, Bernardo L Horta^72^, Wei Huang^201^, Erik Ingelsson^254^, Pankow S James^255^, Marjo-Ritta Jarvelin^256,257,258,259^, Jost B Jonas^260,261,262^, J. Wouter Jukema^116,263^, Pontiano Kaleebu^264^, Robert Kaplan^192,194^, Sharon L.R. Kardia^49^, Norihiro Kato^115^, Sirkka M. Keinanen-Kiukaanniemi^265,266^, Bong-Jo Kim^78^, Mika Kivimaki^267^, Heikki A. Koistinen^268,269,270^, Jaspal S. Kooner^45,234,235,271^, Antje Körner^122^, Peter Kovacs^138,272^, Diana Kuh^128^, Meena Kumari^273^, Zoltan Kutalik^274,104^, Markku Laakso^169^, Timo A. Lakka^275,276,277^, Lenore J Launer^60^, Karin Leander^278^, Huaixing Li^28^, Xu Lin^28^, Lars Lind^279^, Cecilia Lindgren^12,280,281^, Simin Liu^18^, Ruth J.F. Loos^38,99^, Patrik KE Magnusson^282^, Anubha Mahajan^12^, Andres Metspalu^15^, Dennis O Mook-Kanamori^17,283^, Trevor A Mori^221^, Patricia B Munroe^52,53^, Inger Njølstad^127^, Jeffrey R O'Connell^94^, Albertine J Oldehinkel^153^, Ken K Ong^9^, Sandosh Padmanabhan^284^, Colin N.A. Palmer^285^, Nicholette D Palmer^141^, Oluf Pedersen^48^, Craig E Pennell^43^, David J Porteous^142,286^, Peter P. Pramstaller^96^, Michael A. Province^218^, Bruce M. Psaty^50,253,287^, Lu Qi^288^, Leslie J. Raffel^289^, Rainer Rauramaa^277^, Susan Redline^54,55^, Paul M Ridker^69,290^, Frits R. Rosendaal^17^, Timo E. Saaristo^291,292^, Manjinder Sandhu^293^, Jouko Saramies^294^, Neil Schneiderman^295^, Peter Schwarz^140,296,200^, Laura J. Scott^42,41^, Elizabeth Selvin^37^, Peter Sever^271^, Xiao-ou Shu^87^, P Eline Slagboom^117^, Kerrin S Small^90^, Blair H Smith^297^, Harold Snieder^35^, Tamar Sofer^298,245^, Thorkild I.A. Sørensen^48,299,7,8^, Tim D Spector^90^, Alice Stanton^300^, Claire J Steves^90,301^, Michael Stumvoll^138^, Liang Sun^28^, Yasuharu Tabara^79^, E Shyong Tai^180,40,302^, Nicholas J Timpson^7,8^, Anke Tönjes^138^, Jaakko Tuomilehto^303,304,305^, Teresa Tusie^77,306^, Matti Uusitupa^307^, Pim van der Harst^120,24^, Cornelia van Duijn^85,62^, Veronique Vitart^203^, Peter Vollenweider^308^, Tanja GM Vrijkotte^129^, Lynne E Wagenknecht^309^, Mark Walker^310^, Ya X Wang^261^, Nick J Wareham^9^, Richard M Watanabe^125,231,126^, Hugh Watkins^152,12^, Wen B Wei^311^, Ananda R Wickremasinghe^312^, Gonneke Willemsen^26,27^, James F Wilson^80,203^, Tien-Yin Wong^56,197^, Jer-Yuarn Wu^58^, Anny H Xiang^313^, Lisa R Yanek^220^, Loïc Yengo^314^, Mitsuhiro Yokota^315^, Eleftheria Zeggini^111,316,317^, Wei Zheng^87^, Alan B Zonderman^60^, Jerome I Rotter^21^, Anna L Gloyn^11,12,186,318^, Mark I. McCarthy^11,319,186,12^, Josée Dupuis^25^, James B Meigs^320,245,107^, Robert A Scott^9^, Inga Prokopenko^29,22^, Aaron Leong^321,322,236^, Ching-Ti Liu^25^, Stephen CJ Parker^6,323#^, Karen L. Mohlke^3#^, Claudia Langenberg^9#^, Eleanor Wheeler^2,9#^, Andrew P. Morris^324,325,326,12#^, Inês Barroso ^1,2,9,327#$^ and the Meta-Analysis of Glucose and Insulin-related Traits Consortium (MAGIC)^*^

^1^Exeter Centre of Excellence for Diabetes Research (ExCEED), Genetics of Complex Traits, University of Exeter Medical School, University of Exeter, Exeter, UK, ^2^Department of Human Genetics, Wellcome Sanger Institute, Hinxton, Cambridge, UK, ^3^Department of Genetics, University of North Carolina, Chapel Hill, NC, USA, ^4^Department of Biostatistics and Epidemiology, University of Massachusetts, Amherst, MA, USA, ^5^Inserm, Univ Brest, EFS, UMR 1078, GGB, Brest, France, ^6^Department of Computational Medicine and Bioinformatics, University of Michigan, Ann Arbor, MI, USA, ^7^MRC Integrative Epidemiology Unit, University of Bristol, Bristol, UK, ^8^Department of Population Health Sciences, Bristol Medical School, University of Bristol, Bristol, Bristol, UK, ^9^MRC Epidemiology Unit, Institute of Metabolic Science, University of Cambridge, Cambridge, UK, ^10^Department of Biostatistics, School of Public Health, Shandong University, Jinan, Shandong, China, ^11^Oxford Centre for Diabetes, Endocrinology and Metabolism, Radcliffe Department of Medicine, University of Oxford, Oxford, UK, ^12^Wellcome Centre for Human Genetics, University of Oxford, Oxford, UK, ^13^Laboratory for Genomics of Diabetes and Metabolism, RIKEN Centre for Integrative Medical Sciences, Yokohama, Japan, ^14^Medical Research Council Human Genetics Unit, Institute for Genetics and Molecular Medicine, Edinburgh, UK, ^15^Estonian Genome Center, Institute of Genomics, University of Tartu, Tartu, Estonia, ^16^COPSAC, Copenhagen Prospective Studies on Asthma in Childhood, Herlev and Gentofte Hospital, University of Copenhagen, Copenhagen, Denmark, ^17^Department of Clinical Epidemiology, Leiden University Medical Center, Leiden, The Netherlands, ^18^Department of Epidemiology, Brown University School of Public Health, Brown University, Providence, RI, USA, ^19^Department of Biomedical Sciences, City University of Hong Kong, Hong Kong SAR, China, ^20^Department of Electrical Engineering, City University of Hong Kong, Hong Kong SAR, China, ^21^The Institute for Translational Genomics and Population Sciences, Department of Pediatrics, The Lundquist Institute for Biomedical Innovation at Harbor-UCLA Medical Center, Torrance, CA, USA, ^22^Department of Metabolism, Digestion, and Reproduction, Imperial College London, London, UK, ^23^Division of Preventive Medicine, Brigham and Women's Hospital, Boston, MA, USA, ^24^Department of Genetics, University of Groningen, University Medical Center Groningen, Groningen, The Netherlands, ^25^Department of Biostatistics, Boston University School of Public Health, Boston, MA, USA, ^26^Department of Biological Psychology, Faculty of Behaviour and Movement Sciences, Vrije Universiteit Amsterdam, Amsterdam, The Netherlands, ^27^Amsterdam Public Health Research Institute, Amsterdam Universities Medical Center, Amsterdam, The Netherlands, ^28^CAS Key Laboratory of Nutrition, Metabolism and Food Safety, Shanghai Institute of Nutrition and Health, University of Chinese Academy of Sciences, Chinese Academy of Sciences, Shanghai, China, ^29^Section of Statistical Multi-omics, Department of Clinical and Experimental Research, University of Surrey, Guildford, Surrey, UK, ^30^Department of Biostatistics, University of Washington, Seattle, WA, USA, ^31^SYNLAB Academy, SYNLAB Holding Deutschland GmbH, Mannheim, Germany, ^32^Clinical Institute of Medical and Chemical Laboratory Diagnostics, Medical University Graz, Graz, Austria, ^33^Vth Department of Medicine (Nephrology, Hypertensiology, Rheumatology, Endocrinology, Diabetology), Medical Faculty Mannheim, Heidelberg University, Mannheim, Baden-Württemberg, Germany, ^34^Department of Economics, Metropolitan Autonomous University, Mexico City, Mexico, ^35^Department of Epidemiology, University of Groningen, University Medical Center Groningen, Groningen, The Netherlands, ^36^CVD Genetic Epidemiology Computational Laboratory, Gillings School of Global Public Health, University of North Carolina, Chapel Hill, NC, USA, ^37^Department of Epidemiology, Johns Hopkins Bloomberg School of Public Health, Baltimore, MD, USA, ^38^The Charles Bronfman Institute for Personalized Medicine, Icahn School of Medicine at Mount Sinai, New York, NY, USA, ^39^HPI Digital Health Center, Digital Health and Personalized Medicine, Hasso Plattner Institute, Potsdam, Germany, ^40^Saw Swee Hock School of Public Health, National Univeristy of Singapore and National University Health System, Singapore, Singapore, ^41^Center for Statistical Genetics, University of Michigan, Ann Arbor, MI, USA, ^42^Department of Biostatistics, School of Public Health, University of Michigan, Ann Arbor, MI, USA, ^43^School of Medicine and Public Health, College of Health, Medicine and Wellbeing, The University of Newcastle, Newcastle, NSW, Australia, ^44^Department of Epidemiology and Biostatistics, Imperial College London, London, UK, ^45^Department of Cardiology, Ealing Hospital, London North West Healthcare NHS Trust, Middlesex, UK, ^46^Steno Diabetes Center Copenhagen, Gentofte, Denmark, ^47^The Bioinformatics Centre, Department of Biology, University of Copenhagen, Copenhagen, Denmark, ^48^Novo Nordisk Foundation Center for Basic Metabolic Research, Faculty of Health and Medical Sciences, University of Copenhagen, Copenhagen, Denmark, ^49^Department of Epidemiology, School of Public Health, University of Michigan, Ann Arbor, MI, USA, ^50^Department of Medicine, Cardiovascular Health Research Unit, University of Washington, Seattle, WA, USA, ^51^Metabolism Program, Program in Medical and Population Genetics, Broad Institute, Cambridge, MA, USA, ^52^Department of Clinical Pharmacology, William Harvey Research Institute, Barts and The London School of Medicine and Dentistry, Queen Mary University of London, London, UK, ^53^NIHR Barts Cardiovascular Biomedical Research Centre, Queen Mary University of London, London, UK, ^54^Department of Medicine, Sleep and Circadian Disorders, Brigham and Women's Hospital, Boston, MA, USA, ^55^Department of Medicine, Sleep Medicine, Harvard Medical School, Boston, MA, USA, ^56^Ocular Epidemiology, Singapore Eye Research Institute, Singapore National Eye Centre, Singapore, Singapore, ^57^Department of Ophthalmology, National University of Singapore and National University Health System, Singapore, Singapore, ^58^Institute of Biomedical Sciences, Academia Sinica, Taipei, Taiwan, Taiwan, ^59^Department of Epidemiology, The Herbert Wertheim School of Public Health and Human Longevity Science, UC San Diego, La Jolla, CA, USA, ^60^Laboratory of Epidemiology and Population Sciences, National Institute on Aging, National Institutes of Health, Baltimore, MD, USA, ^61^Institute of Population Health Sciences, National Health Research Institutes, Miaoli, Taiwan, ^62^Department of Epidemiology, Erasmus Medical Center, Rotterdam, The Netherlands, ^63^Department of Statistics, University of North Carolina at Chapel Hill, Chapel Hill, NC, USA, ^64^Institute of Cardiovascular Science, UCL, London, UK, ^65^Uganda Medical Informatics Centre (UMIC), MRC/UVRI and London School of Hygiene & Tropical Medicine (Uganda Research Unit), Entebbe, Uganda, ^66^London School of Hygiene & Tropical Medicine, London, UK, ^67^H3Africa Bioinformatics Network (H3ABioNet) Node, Centre for Genomics Research and Innovation, NABDA/FMST, Abuja, Nigeria, ^68^Bioinfosol, Sevilla, Spain, ^69^Division of Preventive Medicine, Brigham and Women's Hospital, Boston, MA, USA, ^70^Department of Medical Sciences, Molecular Epidemiology and Science for Life Laboratory, Uppsala University, Uppsala, Sweden, ^71^Department of Statistics, The University of Auckland, Science Center, Auckland, New Zealand, ^72^Postgraduate Program in Epidemiology, Federal University of Pelotas, Pelotas, RS, Brazil, ^73^Department of Medicine, Epidemiology, Vanderbilt University Medical Center, Nashville, TN, USA, ^74^Department of Epidemiology, Tulane University Obesity Research Center,, Tulane University, New Orleans, USA, ^75^Department of Epidemiology and Biostatistics, School of Public Health, Peking University, Beijing, China, ^76^Molecular Biology and Genomic Medicine Unit, National Council for Science and Technology, Mexico City, Mexico, ^77^Molecular Biology and Genomic Medicine Unit, National Institute of Medical Sciences and Nutrition, Mexico City, Mexico, ^78^Division of Genome Science, Department of Precision Medicine, National Institute of Health, Cheongju-si, Chungcheongbuk-do, South Korea, ^79^Center for Genomic Medicine, Kyoto University Graduate School of Medicine, Kyoto, Japan, ^80^Centre for Global Health Research, Usher Institute, University of Edinburgh, Edinburgh, Scotland, ^81^Centre for Cardiovascular Sciences, Queen's Medical Research Institute, University of Edinburgh, Edinburgh, Scotland, ^82^Department of Medicine, Divison of Biomedical Informatics and Personalized Medicine, University of Colorado Anschutz Medical Campus, Denver, CO, USA, ^83^Department of Biostatistics and Data Science, Wake Forest School of Medicine, Winston-Salem, NC, USA, ^84^Department of Medicine, Division of Nephrology and Hypertension, University of Utah, Salt Lake City, UT, USA, ^85^Nuffield Department of Population Health, University of Oxford, Oxford, UK, ^86^Lee Kong Chian School of Medicine, Nanyang Technological University, Singapore, Singapore, ^87^Division of Epidemiology, Department of Medicine, Vanderbilt Epidemiology Center, Vanderbilt University Medical Center, Nashville, TN, USA, ^88^Department of Clinical Science, Center for Diabetes Research, University of Bergen, Bergen, Norway, ^89^Department of Clinical Sciences, Lund University Diabetes Centre, Lund University, Malmo, Sweden, ^90^Department of Twin Research and Genetic Epidemiology, School of Life Course Sciences, King's College London, London, UK, ^91^NIHR Biomedical Research Centre, Guy’s and St Thomas’ Foundation Trust, London, UK, ^92^Institute of Epidemiology, Research Unit of Molecular Epidemiology, Helmholtz Zentrum München Research Center for Environmental Health, Neuherberg, Bavaria, Germany, ^93^German Center for Diabetes Research (DZD), Neuherberg, Bavaria, Germany, ^94^Department of Medicine, Division of Endocrinology, Diabetes, and Nutrition, University of Maryland School of Medicine, Baltimore, MD, USA, ^95^Public Health Informatics Unit, Department of Integrated Sciences, Nagoya University Graduate School of Medicine, Nagoya, Japan, ^96^Institute for Biomedicine, Eurac Research, Bolzano, BZ, Italy, ^97^Department of Internal Medicine, Section of Gerontology and Geriatrics, Leiden University Medical Center, Leiden, The Netherlands, ^98^Istituto di Ricerca Genetica e Biomedica (IRGB), Consiglio Nazionale delle Ricerche (CNR), Monserrato, Italy, ^99^The Mindich Child Health and Development Institute for Personalized Medicine, Icahn School of Medicine at Mount Sinai, New York, NY, USA, ^100^Department of Preventive Medicine, Northwestern University Feinberg School of Medicine, Chicago, IL, USA, ^101^Center for Public Health Genomics, University of Virginia, Charlottesville, VA, USA, ^102^Department of Public Health Sciences, University of Virginia, Charlottesville, VA, USA, ^103^Department of Computational Biology, University of Lausanne, Lausanne, Switzerland, ^104^Swiss Institute of Bioinformatics, Lausanne, Switzerland, ^105^Center for Genomic Medicine, Massachusetts General Hospital, Harvard Medical School, Boston, MA, USA, ^106^Department of Anesthesia, Critical Care and Pain Medicine, Massachusetts General Hospital, Boston, MA, USA, ^107^Program in Medical and Population Genetics,, Broad Institute, Cambridge, MA, USA, ^108^Department of Cell and Molecular Biology., National Bioinformatics Infrastructure Sweden,, Science for Life Laboratory, Uppsala University, Uppsala, Sweden, ^109^Department of Biostatistics, University of Michigan, Ann Arbor, MI, USA, ^110^Icelandic Heart Association, Kopavogur, Iceland, ^111^Institute of Translational Genomics, Helmholtz Zentrum München – German Research Center for Environmental Health, Neuherberg, Germany, ^112^Wellcome Sanger Institute, Hinxton, Cambridge, UK, ^113^Institute of Health and Wellbeing, University of Glasgow, Glasgow, Glasgow, UK, ^114^Department of Medicine Solna, Cardiovascular medicine, Karolinska Institutet, Stockholm, Sweden, ^115^National Center for Global Health and Medicine, Tokyo, Japan, ^116^Department of Cardiology, Leiden University Medical Center, Leiden, The Netherlands, ^117^Department of Biomedical Data Sciences, Molecular Epidemiology, Leiden University Medical Center, Leiden, The Netherlands, ^118^Department of Pattern Recognition & Bioinformatics, Delft University of Technology, Delft, The Netherlands, ^119^Department of Biomedical Data Sciences, Leiden Computational Biology Center, Leiden University Medical Center, Leiden, The Netherlands, ^120^Department of Cardiology, University of Groningen, University Medical Center Groningen, Groningen, The Netherlands, ^121^Genomics plc, Oxford, UK, ^122^Center of Pediatric Research, University Children´s Hospital Leipzig, University of Leipzig Medical Center, Leipzig, Germany, ^123^Department of Epidemiology and Biostatistics, School of Public Health, Tongji Medical College, Huazhong University of Science and Technology, Wuhan, China, ^124^Genome Institute of Singapore, Agency for Science, Technology and Research, Singapore, Singapore, ^125^Department of Preventive Medicine, Keck School of Medicine of USC, Los Angeles, CA, USA, ^126^USC Diabetes and Obesity Research Institute, Keck School of Medicine of USC, Los Angeles, CA, USA, ^127^Department of Community Medicine, Faculty of Health Sciences, UIT the Arctic University of Norway, Tromsø, Norway, ^128^MRC Unit for Lifelong Health & Ageing at UCL, London, UK, ^129^Department of Public Health, Amsterdam Public Health Research Institute, Amsterdam Universities Medical Center, Amsterdam, The Netherlands, ^130^Department of Clinical Epidemiology, Biostatistics, and Bioinformatics, Amsterdam Public Health Research Institute, Amsterdam Universities Medical Center, Amsterdam, The Netherlands, ^131^Department of Public Health and Primary Care, School of Clinical Medicine, University of Cambridge, Cambridge, UK, ^132^Department of Nutrition, Exercise, and Sports, Faculty of Science, University of Copenhagen, Copenhagen, Denmark, ^133^Department of Internal Medicine, University of Groningen, University Medical Center Groningen, Groningen, The Netherlands, ^134^Department of Medical Biotechnology and Translational Medicine, University of Milan, Milan, Italy, ^135^Centro Cardiologico Monzino, IRCCS, Milan, Italy, ^136^Diabetes and Obesity Research Institute, Cedars-Sinai Medical Center, Los Angeles, CA, USA, ^137^Department of Epidemiology and Prevention, Division of Public Health Sciences, Wake Forest School of Medicine, Winston-Salem, NC, USA, ^138^Medical Department III – Endocrinology, Nephrology, Rheumatology, University of Leipzig Medical Center, Leipzig, Germany, ^139^Medical Genomics and Metabolic Genetics Branch, National Human Genome Research Institute, National Institues of Health, Bethesda, MD, USA, ^140^Department for Prevention and Care of Diabetes, Faculty of Medicine Carl Gustav Carus, Technische Universität Dresden, Dresden, Germany, ^141^Department of Biochemistry, Wake Forest School of Medicine, Winston-Salem, NC, USA, ^142^Centre for Genomic and Experimental Medicine, Institute of Genetics & Molecular Medicine, University of Edinburgh, Western General Hospital, Edinburgh, UK, ^143^Usher Institute, University of Edinburgh, Edinburgh, UK, ^144^Department of Internal Medicine, National Taiwan University Hospital, Taipei, Taiwan, ^145^Graduate Institute of Medical Genomics and Proteomics, National Taiwan University, Taipei, Taiwan, ^146^Institute of Biomedical Sciences, Academia Sinica, Taipei, Taiwan, ^147^Department of Nutrition, Gillings School of Global Public Health, University of North Carolina, Chapel Hill, NC, USA, ^148^Department of Population Science and Experimental Medicine, Institute of Cardiovascular Science, University College London, London, UK, ^149^Department of Nutrition and Dietetics, School of Health Science and Education, Harokopio University of Athens, Athens, Greece, ^150^Department of Medicine Solna, Cardiovascular medicine, Stockholm, Sweden, ^151^Department of Epidemiology, Shanghai Cancer Institute, Shanghai, China, ^152^Division of Cardiovascular Medicine, Radcliffe Department of Medicine, University of Oxford, Oxford, UK, ^153^Department of Psychiatry, Interdisciplinary Center Psychopathy and Emotion Regulation, University of Groningen, University Medical Center Groningen, Groningen, The Netherlands, ^154^Institute for Clinical Diabetology, German Diabetes Center, Leibniz Center for Diabetes Research at Heinrich Heine University Düsseldorf, Düsseldorf, Germany, ^155^Division of Endocrinology and Diabetology, Medical Faculty, Heinrich Heine University Düsseldorf, Düsseldorf, Germany, ^156^German Center for Diabetes Research (DZD), Düsseldorf, Germany, ^157^Internal Medicine, Endocrine & Metabolism, Tri-Service General Hospital, Taipei, Taiwan, ^158^School of Medicine, National Defense Medical Center, Taipei, Taiwan, ^159^Internal Medicine, Endocrinology, Diabetes & Metabolism, Diabetes and Metabolism Research Center, The Ohio State University Wexner Medical Center, Columbus, OH, USA, ^160^Department of Environmental and Preventive Medicine, Jichi Medical University School of Medicine, Shimotsuke, Japan, ^161^Department of Anti-aging Medicine, Ehime University Graduate School of Medicine, Toon, Japan, ^162^National Institute of Public Health, University of Southern Denmark, Odense, Denmark, ^163^Department of Medicine, Endocrinology, Diabetes & Metabolism, Johns Hopkins University School of Medicine, Baltimore, MD, USA, ^164^Clinical Diabetes, Endocrinology & Metabolism, Translational Research & Cellular Therapeutics, Beckman Research Institute of the City of Hope, Duarte, CA, USA, ^165^Department of Clinical Gene Therapy, Osaka University Graduate School of Medicine, Suita, Japan, ^166^Department of Geriatric and General Medicine, Osaka University Graduate School of Medicine, Suita, Japan, ^167^Department of Public Health, University of Split School of Medicine, Split, Croatia, ^168^Institute of Biomedicine, Bioinformatics Center, Univeristy of Eastern Finland, Kuopio, Finland, ^169^Department of Medicine, University of Eastern Finland and Kuopio University Hospital, Kuopio, Finland, ^170^MRC Integrative Epidemiology Unit, University of Bristol, Bristol, Bristol, UK, ^171^USC-Office of Population Studies Foundation, University of San Carlos, Cebu City, Philippines, ^172^Department of Anthropology, Sociology and History, University of San Carlos, Cebu City, Philippines, ^173^State Key Laboratory of Oncogene and Related Genes & Department of Epidemiology, Shanghai Cancer Institute, Renji Hospital, Shanghai Jiaotong University School of Medicine, Shanghai, China, ^174^Internal Medicine, Endocrine & Metabolism, Taichung Veterans General Hospital, Taichung, Taiwan, ^175^Center for Geriatrics and Gerontology,, Taichung Veterans General Hospital, Taichung, Taiwan, ^176^National Defense Medical Center, National Yang-Ming University, Taipei, Taiwan, ^177^Diabetes Prevention Unit, National Institute for Health and Welfare, Helsinki, Finland, ^178^Center for Clinical Research and Prevention, Bispebjerg and Frederiksberg Hospital, Copenhagen, Denmark, ^179^Department of Clinical Medicine, Faculty of Health and Medical Sciences, University of Copenhagen, Copenhagen, Denmark, ^180^Yong Loo Lin School of Medicine, National University of Singapore and National University Health System, Singapore, Singapore, ^181^Department of Medicine, University of Texas Health Sciences Center, San Antonio, TX, USA, ^182^Department of Internal Medicine, Aichi Gakuin University School of Dentistry, Nagoya, Japan, ^183^Department of Diabetes, Diabetes, & Nutritional Sciences, James Black Centre, King's College London, London, UK, ^184^Department of Functional Pathology, Shimane University School of Medicine, Izumo, Japan, ^185^Department of Medicine and Pharmacology, New York Medical College School of Medicine, Valhalla, NY, USA, ^186^Oxford NIHR Biomedical Research Centre, Oxford University Hospitals NHS Foundation Trust, Oxford, UK, ^187^Colorado School of Public Health, University of Colorado Anschutz Medical Campus, Aurora, CO, USA, ^188^Department of Geriatric Medicine and Neurology, Ehime University Graduate School of Medicine, Toon, Japan, ^189^Institute of Epidemiology, Helmholtz Zentrum München Research Center for Environmental Health, Neuherberg, Bavaria, Germany, ^190^Institute for Medical Information Processing, Biometry, and Epidemiology, Ludwig-Maximilians University Munich, Munich, Bavaria, Germany, ^191^Gen-info LtD, Zagreb, Croatia, ^192^Department of Epidemiology and Population Health, Albert Einstein College of Medicine, Bronx, NY, USA, ^193^Genetics and Pharmacogenomics, Merck Sharp & Dohme Corp., Kenilworth, NJ, USA, ^194^Department of Public Health Sciences, Fred Hutchinson Cancer Research Center, Seattle, WA, USA, ^195^Department of Internal Medicine, Erasmus Medical Center, Rotterdam, The Netherlands, ^196^Centre for Global Health, The Usher Institute, University of Edinburgh, Edinburgh, UK, ^197^Ophthalmology & Visual Sciences Academic Clinical Program (Eye ACP), Duke-NUS Medical School, Singapore, Singapore, ^198^BHF Glasgow Cardiovascular Research Centre, Institute of Cardiovascular and Medical Sciences, University of Glasgow, Glasgow, UK, ^199^Department of Experimental Diabetology, German Institute of Human Nutrition Potsdam-Rehbruecke, Nuthetal, Germany, ^200^German Center for Diabetes Research (DZD e.V.), Neuherberg, Germany, ^201^Department of Genetics, Shanghai-MOST Key Laboratory of Health and Disease Genomics, Chinese National Human Genome Center at Shanghai (CHGC) and Shanghai Academy of Science & Technology (SAST), Shanghai, China, ^202^Sarepta Therapeutics, Cambridge, Massachusetts, USA, ^203^Medical Research Council Human Genetics Unit, Institute for Genetics and Cancer, University of Edinburgh, Edinburgh, UK, ^204^Department of Laboratory Medicine and Pathology, University of Minnesota, Minneapolis, MN, USA, ^205^Department of Nutrition, Harvard T.H. Chan School of Public Health, Boston, MA, USA, ^206^Institute of Clinical Medicine, Internal Medicine, University of Eastern Finland, Kuopio, Finland, ^207^Department of Medicine, Bornholms Hospital, Rønne, Denmark, ^208^Department of Internal Medicine, Division of Endocrinology, Leiden University Medical Center, Leiden, The Netherlands, ^209^Laboratory for Experimental Vascular Medicine, Leiden University Medical Center, Leiden, The Netherlands, ^210^Department of Human Genetics, Leiden University Medical Center, Leiden, The Netherlands, ^211^Department of Human Biology, University of Split School of Medicine, Split, Croatia, ^212^Carolina Population Center, University of North Carolina, Chapel Hill, NC, USA, ^213^Department of Endocrinology and Metabolism, Instituto Nacional de Ciencias Medicas y Nutricion, Mexico City, Mexico, ^214^Unidad de Investigación de Enfermedades Metabólicas, Instituto Nacional de Ciencias Médicas y Nutrición and Tec Salud, Mexico City, Mexico, ^215^Instituto Tecnológico y de Estudios Superiores de Monterrey Tec Salud, Mexico City, Mexico, ^216^Department of Medical Genomics, Pfizer/University of Granada/Andalusian Government Center for Genomics and Oncological Research (GENYO), Granada, Spain, ^217^Institute for Environmental Medicine, Chronic Inflammatory Diseases, Karolinska Institutet, Solna, Sweden, ^218^Department of Genetics, Division of Statistical Genomics, Washington University School of Medicine, St. Louis, MO, USA, ^219^Clinical and Health Services Research, National Institute on Minority Health and Health Disparities, Bethesda, MD, USA, ^220^Department of Medicine, General Internal Medicine, Johns Hopkins University School of Medicine, Baltimore, MD, USA, ^221^Medical School, Royal Perth Hospital Unit, University of Western Australia, Perth, WA, Australia, ^222^Department of Integrative Biomedical Sciences, University of Cape Town, Cape Town, South Africa, ^223^Aberdeen Centre for Health Data Science, 1:042 Polwarth Building,, School of Medicine, Medical, Science and Nutrition, University of Aberdeen, Foresterhill, Aberdeen, UK, ^224^Human Genetics Center, School of Public Health, The University of Texas Health Science Center at Houston, Houston, TX, USA, ^225^Human Genome Sequencing Center, Baylor College of Medicine, Houston, TX, USA, ^226^Division of Endocrinology and Diabetes, Graduate School of Molecular Endocrinology and Diabetes, University of Ulm, Ulm, Baden-Württemberg, Germany, ^227^LKC School of Medicine, Nanyang Technological University, Singapore and Imperial College London, UK, Singapore, Singapore, ^228^Hasso Plattner Institute for Digital Health at Mount Sinai, Icahn School of Medicine at Mount Sinai, New York, NY, USA, ^229^Digital Health Center, Hasso Plattner Institut, University Potsdam, Potsdam, Germany, ^230^Department of Medicine, Keck School of Medicine of USC, Los Angeles, CA, USA, ^231^Department of Physiology and Neuroscience, Keck School of Medicine of USC, Los Angeles, CA, USA, ^232^INSERM UMR 1283 / CNRS UMR 8199, European Institute for Diabetes (EGID), Université de Lille, Lille, France, ^233^INSERM UMR 1283 / CNRS UMR 8199, European Institute for Diabetes (EGID), Institut Pasteur de Lille, Lille, France, ^234^Imperial College Healthcare NHS Trust, Imperial College London, London, UK, ^235^MRC-PHE Centre for Environment and Health, Imperial College London, London, UK, ^236^Harvard Medical School, Boston, MA, USA, ^237^Department of Medicine, Jackson Heart Study, University of Mississippi Medical Center, Jackson, MS, USA, ^238^Department of Medicine, Faculty of Medicine, University of Kelaniya, Ragama, Sri Lanka, ^239^Department of Nutrition and Dietetics, School of Health Science and Education, Harokopio University of Athens, Kallithea, Greece, ^240^Department of Clinical Sciences, Lund University, Malmö, Sweden, ^241^Laboratory of Epidemiology and Population Sciences, National Institute on Aging Intramural Research Program, National Institutes of Health, Baltimore, MD, USA, ^242^CNR Institute of Clinical Physiology, Pisa, Italy, ^243^Intramural Research Program, National Institute of Aging, Baltimore, MD, USA, ^244^Diabetes Unit and Center for Genomic Medicine, Massachusetts General Hospital, Boston, MA, USA, ^245^Department of Medicine, Harvard Medical School, Boston, MA, USA, ^246^Department of Public Health and Clinical Medicine, Umeå University, Umeå, Sweeden, ^247^Department of Genomics of Common Disease, Imperial College London, London, UK, ^248^Department of Medicine, Cardiovascular medicine, Karolinska Institutet, Stockholm, Sweden, ^249^Department of Medicine, Division of Endocrinology, Diabetes & Metabolism, Cedars-Sinai Medical Center, Los Angeles, CA, USA, ^250^Diabetes Centre, Lund University, Sweden, ^251^Finnish Instituter of Molecular Medicine, Helsinki University, Helsinki, Finland, ^252^Faculty of Medicine, School of health sciences, University of Iceland, Reykjavik, Iceland, ^253^Department of Epidemiology, Cardiovascular Health Research Unit, University of Washington, Seattle, WA, USA, ^254^Department of Medicine, Division of Cardiovascular Medicine, Stanford University School of Medicine, Stanford University, Stanford, CA, USA, ^255^Division of Epidemiology and Community Health, University of Minnesota, Minneapolis, MN, USA, ^256^Department of Epidemiology and Biostatistics, MRC-PHE Centre for Environment and Health, School of Public Health, Imperial College London, London, UK, ^257^Center for Life Course Health Research, Faculty of Medicine, University of Oulu, Oulu, Finland, ^258^Unit of Primary Health Care, Oulu Univerisity Hospital, OYS, Oulu, Finland, ^259^Department of Life Sciences, College of Health and Life Sciences, Brunel University London, London, UK, ^260^Department of Ophthalmology, Medical Faculty Mannheim, Heidelberg University, Mannheim, Germany, ^261^Beijing Institute of Ophthalmology, Beijing Ophthalmology and Visual Science Key Lab, Beijing Tongren Eye Center, Beijing Tongren Hospital, Capital Medical University, Beijing, China, ^262^Institute of Molecular and Clinical Ophthalmology Basel IOB, Basel, Switzerland, ^263^Netherlands Heart Institute, Utrecht, The Netherlands, ^264^MRC/UVRI and LSHTM (Uganda Research Unit), Entebbe, Uganda, ^265^Faculty of Medicine, Institute of Health Sciences, University of Oulu, Oulu, Finland, ^266^Unit of General Practice, Oulu University Hospital, Oulu, Finland, ^267^Department of Epidemiology and Public Health, UCL, London, UK, ^268^Department of Public Health Solutions, Finnish Institute for Health and Welfare, Helsinki, Finland, ^269^Department of Medicine, University of Helsinki and Helsinki University Central Hospital, Helsinki, Finland, ^270^Minerva Foundation Institute for Medical Research, Helsinki, Finland, ^271^National Heart and Lung Institute, Imperial College London, London, UK, ^272^IFB Adiposity Diseases, University of Leipzig Medical Center, Leipzig, Germany, ^273^Institute for Social and Economic Research, University of Essex, Colchester, UK, ^274^University Institute of Primary Care and Public Health, Division of Biostatistics, University of Lausanne, Lausanne, Switzerland, ^275^Institute of Biomedicine, School of Medicine, University of Eastern Finland, Finland, ^276^Department of Clinical Physiology and Nuclear Medicine, Kuopio University Hospital, Kuopio, Finland, ^277^Foundation for Research in Health Exercise and Nutrition, Kuopio Research Institute of Exercise Medicine, Kuopio, Finland, ^278^Institute of Environmental Medicine, Cardiovascular and Nutritional Epidemiology, Karolinska Institutet, Stockholm, Sweden, ^279^Department of Medical Sciences, Uppsala, Sweden, ^280^Big Data Institute, Nuffield Department of Medicine, University of Oxford, Oxford, UK, ^281^Nuffield Department of Women's and Reproductive Health, University of Oxford, Oxford, UK, ^282^Department of Medical Epidemiology and Biostatistics and the Swedish Twin Registry, Karolinska Institutet, Stockholm, Sweden, ^283^Department of Public Health and Primary Care, Leiden University Medical Center, Leiden, The Netherlands, ^284^Institute of Cardiovascular and Medical Sciences, University of Glasgow, Glasgow, UK, ^285^Division of Population Health and Genomics, School of Medicine, University of Dundee, Ninewells Hospital and Medical School, Dundee, UK, ^286^Centre for Cognitive Ageing and Cognitive Epidemiology, University of Edinburgh, Edinburgh, UK, ^287^Department of Health Services, Cardiovascular Health Research Unit, University of Washington, Seattle, WA, USA, ^288^Department of Epidemiology, Tulane University School of Public Health and Tropical Medicine, New Orleans, LA, USA, ^289^Department of Pediatrics, Genetic and Genomic medicine, University of California, Irvine, Irvine, CA, USA, ^290^Havard Medical School, Boston, MA, USA, ^291^Tampere, Finnish Diabetes Association, Tampere, Finland, ^292^Pirkanmaa Hospital District, Tampere, Finland, ^293^Department of Medicine, University of Cambridge, Cambridge, UK, ^294^South Karelia Central Hospital, Lappeenranta, Finland, ^295^Department of Psychology, University of Miami, Miami, FL, USA, ^296^Paul Langerhans Institute Dresden of the Helmholtz Center Munich, University Hospital and Faculty of Medicine, Dresden, Germany, ^297^Division of Population Health and Genomics, Ninewells Hospital and Medical School, University of Dundee, Dundee, UK, ^298^Division of Sleep and Circadian Disorders, Brigham and Women's Hospital, Boston, MA, USA, ^299^Department of Public Health, Section of Epidemiology, Faculty of Health and Medical Sciences, University of Copenhagen, Copenhagen, Denmark, ^300^Departent of Molecular and Cellular Therapeutics, Royal College of Surgeons in Ireland, Dublin, Ireland, ^301^Department of Aging and Health, Guy’s and St Thomas’ Foundation Trust, London, UK, ^302^Cardiovascular and Metabolic Disease Signature Research Program, Duke-NUS Medical School, Singapore, Singapore, ^303^Department of Public Health Solutions, National Institute for Health and Welfare, Helsinki, Finland, ^304^Department of Public Health, University of Helsinki, Helsinki, Finland, ^305^Saudi Diabetes Research Group, King Abdulaziz University, Jeddah, Saudi Arabia, ^306^Department of Genomic Medicine and Environmental Toxicology, Instituto de Investigaciones Biomedicas, Universidad Nacional Autonoma de Mexico, Mexico City, Mexico, ^307^Department of Public Health and Clinical Nutrition, University of Eastern Finland, Finland, ^308^Department of Medicine, Internal Medicine, Lausanne University Hospital (CHUV), Lausanne, Switzerland, ^309^Department of Public Health Sciences, Wake Forest School of Medicine, Winston-Salem, NC, USA, ^310^Faculty of Medical Sciences, Newcastle University, Newcastle upon Tyne, UK, ^311^Beijing Tongren Eye Center, Beijing Key Laboratory of Intraocular Tumor Diagnosis and Treatment, Beijing Ophthalmology & Visual Sciences Key Lab, Beijing Tongren Hospital, Capital Medical University, Beijing, China, China, ^312^Department of Public Health, Faculty of Medicine, University of Kelaniya, Ragama, Sri Lanka, ^313^Department of Research and Evaluation, Kaiser Permanente of Southern California, Pasadena, CA, USA, ^314^Institute for Molecular Bioscience, The University of Queensland, Queensland, Australia, ^315^Kurume University School of Medicine, Japan, ^316^Wellcome Sanger Institute, Hinxton, UK, ^317^TUM School of Medicine, Technical University of Munich and Klinikum Rechts der Isar, Munich, Germany, ^318^Department of Pediatrics, Division of Endocrinology, Stanford School of Medicine, Stanford, CA, USA, ^319^Wellcome Centre for Human Genetics, Nuffield Department of Medicine, University of Oxford, Oxford, UK, ^320^Department of Medicine, Division of General Internal Medicine, Massachusetts General Hospital, Boston, MA, USA, ^321^Department of Medicine, General Internal Medicine, Massachusetts General Hospital, Boston, MA, USA, ^322^Department of Medicine, Diabetes Unit and Endocrine Unit, Massachusetts General Hospital, Boston, MA, USA, ^323^Department of Human Genetics, University of Michigan, Ann Arbor, MI, USA, ^324^Centre for Genetics and Genomics Versus Arthritis, Division of Musculoskeletal and Dermatological Sciences, The University of Manchester, Manchester, UK, ^325^Centre for Musculoskeletal Research, Division of Musculoskeletal and Dermatological Sciences, The University of Manchester, Manchester, UK, ^326^Department of Biostatistics, University of Liverpool, Liverpool, UK, ^327^University of Cambridge, Cambridge

^#^ Denote shared authorship contributions

^$^ Corresponding author
